# Transposon mutagenesis identifies cooperating genetic drivers during keratinocyte transformation and cutaneous squamous cell carcinoma progression

**DOI:** 10.1101/2019.12.24.887968

**Authors:** Aziz Aiderus, Justin Y. Newberg, Liliana Guzman-Rojas, Ana M. Contreras-Sandoval, Amanda L. Meshey, Devin J. Jones, Felipe Amaya-Manzanares, Roberto Rangel, Jerrold M. Ward, Song-Choon Lee, Kenneth Hon-Kim Ban, Keith Rogers, Susan M. Rogers, Luxmanan Selvanesan, Leslie A. McNoe, Neal G. Copeland, Nancy A. Jenkins, Kenneth Y. Tsai, Michael A. Black, Karen M. Mann, Michael B. Mann

**Author notes:** Correspondence to M.B.M. These authors contributed equally to this work (Aziz Aiderus, Justin Y. Newberg, Liliana Guzman-Rojas). Foundation Medicine, Inc., Cambridge, MA, USA (J.Y.N.); Houston Methodist Cancer Center, Houston Methodist Research Institute, Houston, TX, USA (L.G.-R.); Department of Genetics and Development, Columbia University, New York, NY, USA (D.J.J.); Monoclonal Antibody Core Facility, University of Texas M.D. Anderson Cancer Center, Houston, TX, USA (F.A.-M); Department of Head & Neck Surgery, University of Texas M.D. Anderson Cancer Center, Houston, TX, USA (R.R.); Global VetPathology, Montgomery Village, MD, USA (J.M.W.); Science Centre Singapore, Republic of Singapore (S.-C.L.); Department of Biochemistry, Yong Loo Lin School of Medicine, National University Singapore, Republic of Singapore (K.H.-K.B.); Pacific Edge Limited, Dunedin, Otago, New Zealand (L.S.); AgResearch Invermay Agricultural Centre, Mosgiel, Otago, New Zealand (L.A.M.); and Genetics Department, University of Texas M.D. Anderson Cancer Center, Houston, TX (N.G.C. and N.A.J.).

## Abstract

The systematic identification of genetic events driving cellular transformation and tumor progression in the absence of a highly recurrent oncogenic driver mutation is a challenge in cutaneous oncology. In cutaneous squamous cell carcinoma (cuSCC), the high UV-induced mutational burden poses a hurdle to achieve a complete molecular landscape of this disease. Here, we utilized the *Sleeping Beauty* transposon mutagenesis system to statistically define drivers of keratinocyte transformation and cuSCC progression *in vivo* in the absence of UV-IR, and identified both known tumor suppressor genes and novel oncogenic drivers of cuSCC. Functional analysis confirms an oncogenic role for the *ZMIZ* genes, and tumor suppressive roles for *KMT2C, CREBBP* and *NCOA2*, in the initiation or progression of human cuSCC. Taken together, our *in vivo* screen demonstrates an extremely heterogeneous genetic landscape of cuSCC initiation and progression, which can be harnessed to better understand skin oncogenic etiology and prioritize therapeutic candidates.

## INTRODUCTION

Cutaneous squamous cell carcinoma (cuSCC) is the second most common cancer in man, with approximately one million cases diagnosed annually in the United States. Although the majority of cuSCC are considered a low-risk neoplasm, up to 5% of high-risk cuSCCs are locally or distantly invasive and carry a poor prognosis due to a lack of biomarkers, therapeutic targets, or FDA-approved molecularly targeted therapies. This represents a substantial unmet need for approximately 50,000 patients per year with high-risk cuSCC, and an opportunity to identify new therapeutic modalities that could improve disease outcomes. All non-viral associated skin cancers are thought to require multiple cooperating mutations that deregulate distinct signaling pathways to initiate and progress the multi-step transformation of normal cells into a clinically significant neoplasm. Indeed, identifying cooperating mutations that drive malignant transformation is a prerequisite for developing better combinatorial therapies for managing and treating skin cancers. Most skin cancers, including cuSCC (Pickering et al., 2014; South et al., 2014), have the highest mutation rates among human cancers due to ultraviolet irradiation (UV-IR) induced damage from chronic, intermittent sun exposure. Thus, using human cancer sequencing data alone, with some of the highest mutational burdens of any cancer, poses challenges to identify cooperating, low-penetrant mutations that lead to cancer progression. This presents a need to develop *in vivo* model systems to help identify and prioritize novel cooperating candidate cancer drivers for keratinocyte transformation and subsequent progression to late-stage, invasive cuSCC.

*Sleeping Beauty* (SB) insertional mutagenesis (Ivics et al., 1997) is a powerful tool used to perform genome-wide forward genetic screens in laboratory mice for cancer gene discovery (Collier and Largaespada, 2007; Copeland and Jenkins, 2010; Dupuy et al., 2005; Dupuy et al., 2009; Mann et al., 2013; Mann et al., 2016b; Mann et al., 2012; Mann et al., 2015; Mann et al., 2014b; Rangel et al., 2016; Takeda et al., 2015) in animal models of both hematopoietic and solid tumors (Mann et al., 2014a; Mann et al., 2014b). SB transposons can identify early cancer progression drivers that cooperate to initiate tumors (Mann et al., 2015; Takeda et al., 2015), and potentially drive metastasis (Genovesi et al., 2013; Perez-Mancera et al., 2012). Importantly, SB insertions induce changes in gene expression, thus providing epigenetic information not easily obtained from carcinogenesis mouse models using chemical (Nassar et al., 2015) or chronic UV irradiation (Chitsazzadeh et al., 2016; Knatko et al., 2017) carcinogenesis mouse models or from limited patient samples. We demonstrate that SB mobilization of a low-copy T2/Onc3 transposon allele is sufficient to induce and progress a variety of cancers *in vivo*. Here, we report our efforts for cancer gene discovery in skin tumors. We focused on the analysis of skin tumors from these mice and identified several oncogenic and many tumor suppressor driver genes. Using high-throughput sequencing approaches (Mann et al., 2016b; Mann et al., 2015) to identify genome-wide SB mutations. Using our SB Driver Analysis (Newberg et al., 2018b) statistical framework, we profiled genome-wide SB mutations from early- and late-stage cuSCC and defined recurrently mutated, statistically significant candidate cancer drivers (CCDs) from bulk cuSCC tumors and from normal keratinocytes and early stage tumors, identifying both known tumor suppressor genes and novel oncogenic drivers. We further prioritized oncogenic and tumor suppressor candidates, and provide *in vitro* and *in vivo* functional evidence for the roles of these genes in the initiation and progression of cuSCC. Taken together, our efforts provide a systematic evolutionary landscape of cuSCC genesis, and highlights key pathways promoting disease that are potential therapeutic targets.

## RESULTS

### Candidate cancer driver discovery in SB-driven keratinocyte cancer models

We performed a forward genetic screen to define the tumor spectrum driven by the cooperation of SB with Trp53 mutant alleles and the underlying cooperating genes (**Fig. 1a** and **Supplementary Fig. 1a**). Whole-body mutagenesis was achieved using low-copy, bi-functional SB transposons from SB|Onc3 mice carrying an inducible SB transposase (Dupuy et al., 2009), Onc3 transposon (Dupuy et al., 2009), and *Actb-Cre* transgene (Lewandoski et al., 1997). The addition of either a conditional *Trp53* null (*Trp53*^*KO/+*^) (Jonkers et al., 2001) or recurrent point mutant allele (*Trp53*^*R172H/+*^) (Olive et al., 2004) accelerated tumor progression and significantly reduced tumor-free survival relative to *Trp53* wild-type littermate controls (*P*<0.0001, **Fig. 1a**). SB|Onc3 mice succumbed to a wide variety of solid and hematopoietic tumor types with variable penetrance (**Supplementary Table 1**). SB|*Trp53*^*KO/+*^ mice were the earliest to reach endpoint, but had the lowest overall tumor burden, presumably because these mice had the highest proportion of hematopoietic disease (**Supplementary Table 1**). SB|*Trp53*^*R172H/+*^ mice reached endpoint earlier than SB|*Trp53*^*+/+*^ mice and had the widest spectrum of solid tumors (**Supplementary Table 1**). This observation is consistent with previous studies showing that missense mutations in *Trp53* promote malignant transformation by abrogating both the tumor suppressive and oncogenic (gain-of-function and/or dominant negative) functions of TRP53 *in vivo* (Doyle et al., 2010; Hanel et al., 2013; Lang et al., 2004; Morton et al., 2010; Olive et al., 2004). The most prevalent of the more than 20 solid tumor types observed were early-stage cutaneous keratoacanthoma (cuKA) (**Fig. 1b**,**g**) and late-stage, well differentiated cutaneous squamous cell carcinoma (cuSCC) (**Fig. 1c-g**), with a disease penetrance and latency similar to Onc3 cohorts produced with constitutively active SBase (Dupuy et al., 2009). Importantly, these cuSCC/cuKAs were absent in mice with mutant *Trp53* lacking SB (**Fig. 1g**), strongly suggesting SB mutagenesis drives initiation of cellular transformation and subsequent cancer progression of a wide spectrum of tumor types *in vivo*.

**Table 1.**
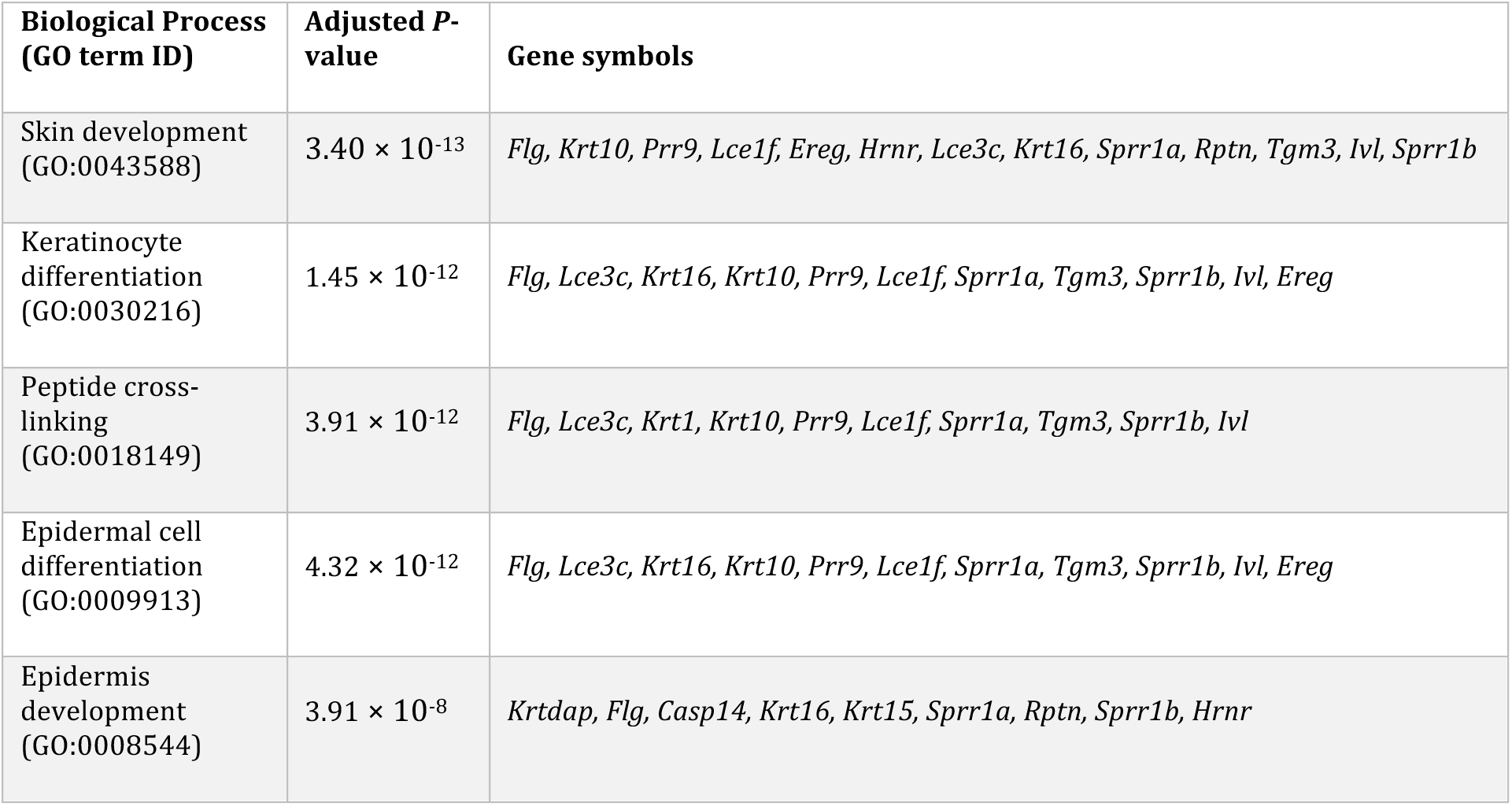
Gene Ontology biological processes enriched in SB-driven cuSCC tumors with *Zmiz1* insertions.

**Figure 1:**
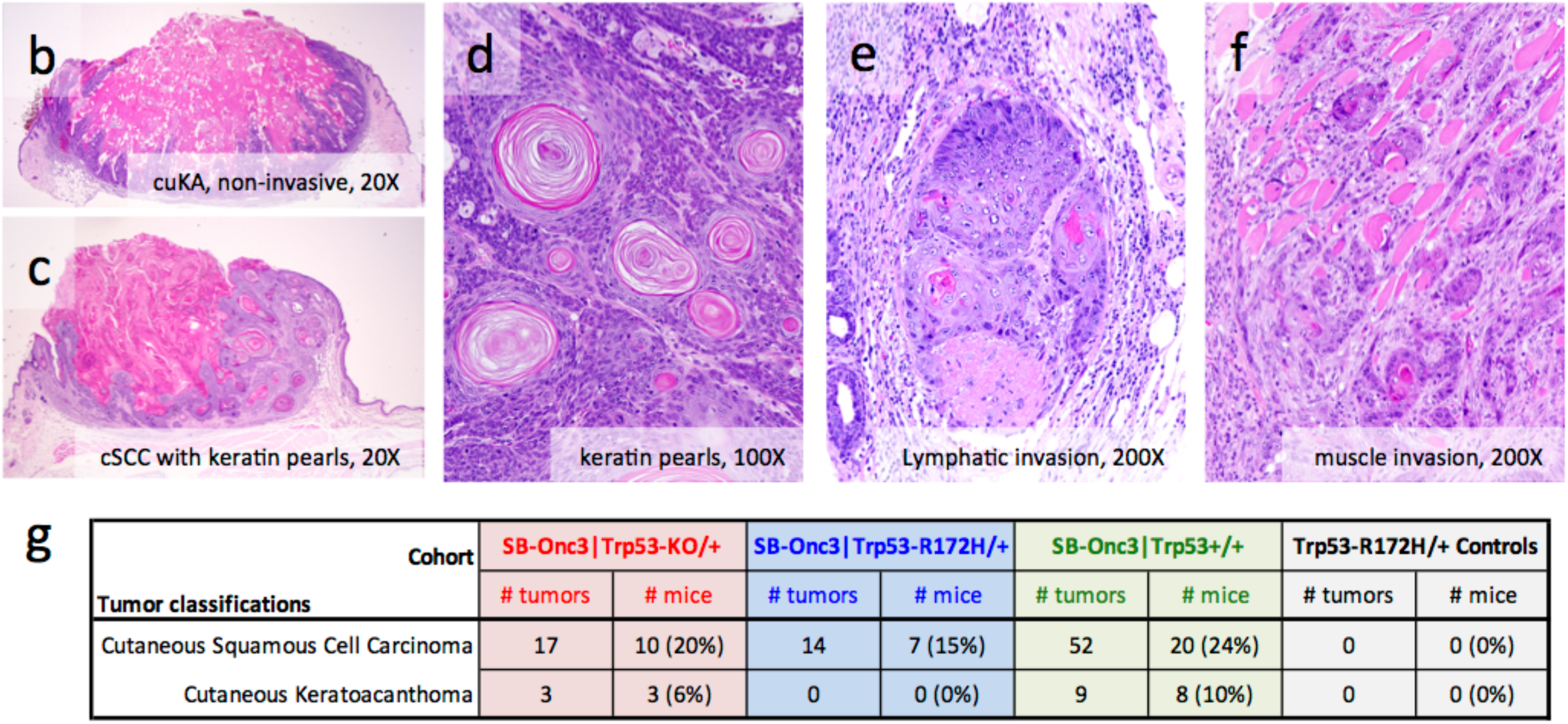
Whole-body SB transposon mutagenesis drives a diverse tumor spectrum in *Trp53* wild type and mutant mice. (**a**) Kaplan-Meier survival curves comparing experimental SB and non-SB control mouse cohorts (log-rank test, *P* < 0.0001, **Supplementary Figure 1**). Wild type *Trp53* cohorts with SB-Onc3 mobilized by a constitutively active *Rosa26*-SBase allele (Gray line, reported in (Dupuy et al., 2009)) do not differ significantly from SB-Onc3 mice mobilized by a conditionally activated *Rosa26*-LSL-SBase allele to the *Rosa26*-1lox-SBase allele by whole body *Cre* expression (log-rank test, *P*>0.05). Mice in all SB-Onc3 cohorts developed solid tumors, including cuSCC and HCA, but *Trp53*^+/−^ (red line) and *Trp53*^R172H/+^ (blue line) mice had significantly decreased survival compared to the WT cohort (Green line, *P*<0.0001, log-rank). Non-SB control Trp53 R172H/+ (broken black line) mice had significantly decreased survival compared to the WT cohort (solid black line, *P*<0.0001, log-rank). (**b-c**) Histology and tumor classification from sections of skin masses stained with hematoxylin and eosin. (**c**) Early-stage, non-invasive mass with cutaneous keratoacanthoma-like morphology in SB-Onc3|*Trp53*^+/+^ mouse (20×). (**d**) Late-stage, invasive mass with cuSCC displaying keratin pearl morphology in SB-Onc3|*Trp53*^KO/+^ mouse (20×). (**f**) cuSCC keratin pearls in SB-Onc3|*Trp53*^+/+^ mouse (100×). (**e**) cuSCC displaying lymphatic invasion in SB-Onc3|*Trp53*^R172H/+^ mouse (200×). (**e**) cuSCC displaying muscle invasion in SB-Onc3|*Trp53*^+/+^ mouse (200×). (**g**) Cutaneous tumor spectrum in SB-Onc3 mice. *Trp53* mutation shortens tumor latency, broadens solid tumor spectrum, and reduces overall tumor burden/multiplicity induced by whole body SB transposon mutagenesis compared to wild type *Trp53* cohorts (**Supplementary Table 1**).

To investigate SB-driven keratinocyte transformation and cuSCC progression in wildtype and Trp53 mutant mice, we sequenced SB insertions sites using the 454_splink method (Mann et al., 2016b) from 83 individual skin masses isolated from 65 SB|Onc3 mice (**Supplementary Table 2–3**), that were histologically confirmed as late-stage cutaneous squamous cell carcinoma (cuSCC). Using gene-guided SB Driver Analysis statistical framework (Newberg et al., 2018b), we identified 31,266 non-redundant transposon insertion sites from the individual (**Supplementary Table 1**) or combined analysis of Trp53 mutant and wild type cuSCC cohorts (**Supplementary Table 4**) which were used to statistically define 428 candidate cancer drivers (CCDs, **Supplementary Table 5**; see **Methods** and **Supplementary Figure 2**). Nearly all CCDs were identified across all cohorts, suggesting that their role in driving cuSCC is independent of *Trp53* mutation (**Supplementary Figure 3a**). For this reason, we focused subsequent data analysis on the superset consisting of all SB|cuSCC masses with all *Trp53* alleles.

Previous studies suggest that most CCDs identified by SB mutagenesis function during tumor progression rather than tumor initiation (Mann et al., 2015). We previously reported a workflow that allow us to distinguish initiating and progression drivers within bulk tumor and single cell sequencing in an SB-driven myeloid leukemia model (Mann et al., 2016b). Stringent filtering of the SB insertions to include only events with the highest sequencing read depths defines trunk driver alterations that are predicted to be under positive-selection within the clonally expanding tumors, where they participate in either initiation or early progression of cellular transformation (Mann et al., 2016b; Mann et al., 2015). To identify CCDs that drive keratinocyte transformation, we selected the SB insertion sites with the highest number of sequencing reads to define trunk drivers (see **Online Methods**), reasoning that tumor initiating insertions would have among the highest frequency in tumor cells. Using SB insertions sites by the 454_splink method, we identified 142 trunk drivers of initiation and/or early progression of keratinocyte transformation *in vivo* (subsequently referred to as trunk drivers) (**Supplementary Table 6** and **Supplementary Figure 3b**). SB insertion maps for representative cuSCC trunk drivers show the locations of mapped SB insertions and define candidate cancer drivers as putative proto-oncogenes (**Supplementary Figure 3c**) or tumor suppressor (**Supplementary Figure 3d**) candidate cuSCC driver genes.

Next, we investigated keratinocyte transformation using keratinocyte-specific SB mutagenesis in *Pten*-sensitized mice. The screen originally was designed to activate an inducible allele of the SB transposase (Rosa26-LSL-SB11) using *Keratin5*-Cre (K5-Cre, Tg(KRT5-cre)5132Jlj, MGI:3050065) (Ramirez et al., 2004) to drive SB insertional mutagenesis in stratified epithelium of the mammary ducts in a haploinsufficient *Pten*^*floxed/+*^ sensitized background (Rangel et al., 2016). However, K5 also induces cre expression in cells of the basal epidermis. Using a two-generation mating scheme, we first introduced K5-Cre onto the background of a Cre-LoxP inducible *Pten* mutant strain, *Pten*^*CKO/+*^ (*Pten*^tm1Hwu^; MGI:2156086). Compound heterozygous progeny (K5-Cre; *Pten*^*CKO/+*^) were mated with double homozygous SB strains containing either a high-copy (T2/Onc2-TG.6113) or a low-copy (T2/Onc3-TG.12740) transposon donor concatemer and the inducible SB transposase (Rosa26-LSL-SBase or SBase^LSL^) to generate experimental and control cohorts (Rangel et al., 2016) (**Supplementary Figure 1a**). Cutaneous keratoacanthomas (cuKA) were diagnosed from routine H&E sections and found to occur in 43% (13 of 30) of quadruple mutant mice (**Supplementary Table 1**), either K5-Cre; *Pten*^*CKO/+*^; TG.6113; SBase^LSL^ or K5-Cre; *Pten*^*CKO/+*^; TG.12740; SBase^LSL^. Four of the mice with cuKA also developed a breast tumor and none of the mice in the control groups developed skin masses before reaching a predetermined experimental end-point at 450 days of age.

To identify SB drivers of cuKA in *Pten*-sensitized mice, we used 45 flash frozen cuKA specimens from 13 quadruple mutant mice (**Supplementary Table 2–3**) collected at necropsy to create genomic DNA and sequenced using the 454-Splink method to sequencing SB insertion events. Using the 454 sequencing data (**Supplementary Table 7**), we performed gene-guided SB Driver Analysis (Newberg et al., 2018b) to define 302 statistically significant candidate cancer drivers (**Supplementary Table 8**) and 29 Trunk drivers (**Supplementary Table 9**). Our top ranked genes were *Chuk* (*IKKA*/*IKK1*), *Notch1*, and *Kctd15* (**Supplementary Figure 4a-b**). All genes were found to have inactivating patterns of SB insertion events into each of their gene-coding regions, strongly suggesting that cooperating tumor suppressors drive early keratinocyte transformation and cuKA initiation *in vivo*. However, three genomes lacking *Chuk* insertions were found to have activating SB insertions into *Zmiz1* that were identical to those observed in the cuSCC cohorts (**Supplementary Figure 4a**). This may suggest that the altered function(s) of *Chuk* in cuKAs may be overlapping with those of the SB-induced *Zmiz1*^ΔN185^ (Rogers et al., 2013) allele.

### SBCapSeq enhances transposon driver detection in normal, premalignant, and tumor skin cell genomes

To confirm and extend our SB analysis from 454 sequencing and define genetic drivers acquired during the evolution of keratinocyte transformation, we developed an enhanced version of SB Capture Sequencing (SBCapSeq) (Mann et al., 2016b) that improves transposon-based, liquid-phase capture experimental workflow optimized for Ion Torrent sequencing from normal and tumor specimen genomes (Mann, *et al*., 2020, manuscript in preparation). We used SBCapSeq to investigate keratinocyte mutational heterogeneity by histological stage.

We generated SBCapSeq libraries using genomic DNA isolated from the necropsy collected flash frozen specimens of 60 cuSCC, 11 cuKA and 32 normal skin genomes (**Fig. 2** and **Supplementary Table 2**). All cuSCC specimens were from independent mice and 59 of 60 were previously sequenced using the Splink_454 method (**Supplementary Table 10**). Comparisons between the sequencing methods allowed for platform-independent sequence validation of clonal drivers and demonstrated the superiority of the SBCapSeq method to identify clonal and sub-clonal insertions based on the total number and read-depth enrichment of genome-wide SB insertion events with biological and technical reproducibility (**Supplementary Figure 5**). Using the SBCapSeq bioinformatics workflow (Mann et al., 2016b) and gene-guided SB Driver Analysis statistical framework (Newberg et al., 2018b), we identified 11,113,694 reads at 59,337 non-redundant transposon insertion sites from cuSCCs (**Supplementary Table 11**) which were used to statistically define 1,333 candidate cancer driver genes (**Supplementary Table 12**). Additionally, we identified 86 trunk drivers (**Supplementary Figure 6a** and **Supplementary Table 13**). Using the same methodology, we next analyzed cuKA specimens harvested either from mice with cuSCC or from an independent age- and genotype-matched cohort (**Supplementary Table 10**) and identified 2,085,280 reads at 6,727 non-redundant transposon insertion sites (**Supplementary Table 14**) defining 192 candidate cancer driver genes (**Supplementary Table 15**) and 10 trunk drivers (**Supplementary Figure 6b** and **Supplementary Table 16**). Finally, we identified 2,910,945 reads at 47,537 non-redundant transposon insertion sites (**Supplementary Table 17**) from normal skin specimens (**Supplementary Table 4**) that defined 547 candidate cancer drivers (**Supplementary Table 18**). No trunk drivers were identified.

**Figure 2:**
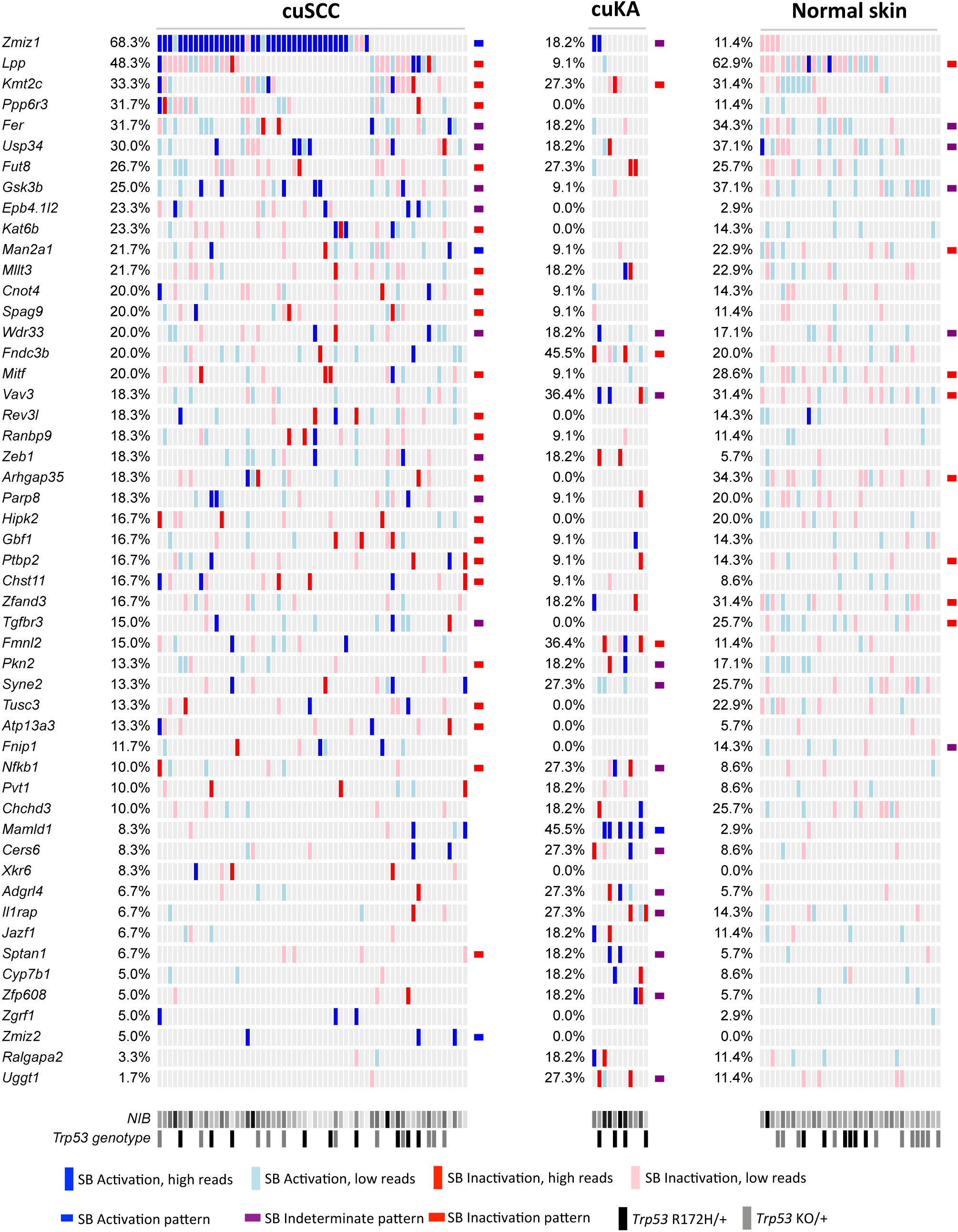
Landscape of trunk drivers mutated during SB-induced keratinocyte transformation and cuSCC progression using SBCapSeq. SB Trunk Driver Analysis, with read depth cutoff of 300, was performed using SBCapSeq insertion data from the SB|cuSCC and SB|cuKA tumor cohorts (the normal skin cohort did not identify any significant trunk drivers). Drivers with significant family-wise error rate (FWER) adjusted *P*-values were merged into a single list and plotted from each of the cuSCC, cuKA and normal skin specimens by occurrence within the cuSCC cohort. Vertical colored bars represent genomes (columns) with SB insertion events that occur within the same (sense, blue) or opposite (anti-sense, red) DNA strand relative to transcription of a driver gene (row). High read depth sites (>299 reads supporting an SB insertion event, blue or red) are distinguished from low read depth sites (<300 reads supporting an SB insertion event, light blue or pink) to denote clonal and sub-clonal SB insertion events, respectively. Driver gene classifications (indeterminate, activating or Inactivating) are shown as a bar at the end of a gene on a cohort specific manner where calculation was possible. NIB, normalized insertion burden from highest (black) to lowest (light gray). *Trp53* genotypes: *Trp53* ^+/+^ (white), *Trp53* ^R172H/+^ (black), and *Trp53* ^KO/+^ (gray).

The histologically normal skin genomes from mice with SB mobilization had a mean of 1,320 SBCapSeq detected insertion events (range 1–3,857; **Supplementary Table 2**), although only 0.82% (389/47,537) of the normal skin insertions had read depths >300 reads per site (range 1–2,297; **Supplementary Table 17**). This is one order of magnitude less than the 23.7% (range 1-19,798) of cuKA genomes or 8.5% (5,014/57,337) of cuSCC genomes with SBCapSeq detected insertion events with >300 reads per specimen (cuKA range 1-19,798; **Supplementary Table 14** and cuSCC range 1–35,809; **Supplementary Table 11**). Clonal selection of recurrent, high-read depth SB insertion sites is featured prominently within the cuSCC and cuKA genomes and almost entirely missing within the normal skin genomes (**Fig. 2**), strongly suggesting that SBCapSeq read depths below ∼300 represent background events and those at 300 and substantially higher represent clonally selected foreground mutation events. Further evidence for clonal selection associated with disease progression was provided by technical replication of SBCapSeq libraries prepared from two independent cuSCC or normal skin specimens (**Supplementary Figure 7**). Independent SBCapSeq libraries demonstrated excellent biological/technical reproducibility in clonal cuSCC genomes (**Supplementary Figure 7a-b**) and no reproducibility in non-clonal normal skin genomes (**Supplementary Figure 7c-d**). Importantly, sequence analysis of the same normal skin SBCapSeq library from independent Ion Torrent runs demonstrated excellent technical reproducibility (**Supplementary Figure 7e-f**), further supporting the robustness of our methodology to detect clonal selection and validating our observation that normal skin does not exhibit clonal selection of SB insertions, despite the high insertion burden. These observations are consistent with the large numbers of mutations observed during chronic UV-IR exposure in human eyelid (Martincorena et al., 2015) and SKH hairless mouse (Chitsazzadeh et al., 2016; Knatko et al., 2017) skin and, together with our SB data, indicates that keratinocytes within phenotypically normal skin can tolerate high background mutagenic insults that may act as a mutational reservoir prior to clonal selection of cooperating driver mutations that permit frank outgrown of cuSCC *in vivo*. It is for this reason that no trunk drivers could be identified from the normal skin specimen genomes. The quantitative, stage-dependent insertional mutation burdens derived from SBCapSeq in cutaneous specimens reveal incredible levels of tumor heterogeneity that has not previously been observed for any SB-driven solid tumor.

### cuKA progression to late-stage cuSCC via clonal and sub-clonal intratumor heterogeneity

We identified four skin masses with histologically continuous but distinct regions of cuKA and cuSCC, suggesting for the first time that cuKA likely progresses directly to late-stage invasive cuSCC *in vivo* (**Supplementary Table 19**). We performed SBCapSeq on representative, histologically distinct regions to investigate intratumor heterogeneity and gain insight into the drivers likely to mediate cuKA to cuSCC progression (**Fig. 3a-c, Supplementary Figure 8a-f**, and **Supplementary Table 20**). Here, the read-depth threshold was adjusted to 200 reads to reflect the lower sequencing depth of the tumor regions. For several loci, we identified insertions at the same nucleotide address across tumor regions from both cuKA and cuSCC in the same mouse, indicating clonal origins of these lesions. Invariably, clonal insertions into *Zmiz1* were identified across each section of all 4 masses (**Fig. 3a**), at three different TA-dinucleotide addresses (**Supplementary Figure 8a-d**), suggesting that *Zmiz1* was under positive selection within the nascent transformed cells that gave rise to the distinct cuKA and cuSCC features. Additional tumor-specific clonal loci include *Fer, Rere*, and *Usp34*, all of which were defined in the population analysis. We also found evidence for sub-clonal insertion events identified at the same nucleotide address in some but not all regions within the same tumor, demonstrating the evolution of the tumor. Some genes with sub-clonal insertion events with read depths greater than 200 were defined as drivers in the bulk population, including *Tcf4* and *Pak2, while other genes, such as Egr2* and *Rab3gap1*, were not (**Fig. 3b-c** and **Supplementary Figure 8e-f**). These data provide evidence for a common keratinocyte progenitor that acquires SB insertions in genes that promote cuKA and are maintained in cuSCC. This analysis also highlights the intra- and inter-tumor heterogeneity of insertions in SB cuSCC tumors and importantly demonstrates that SB insertion profiles can be used to trace the evolution of cuKA to cuSCC using both the insertion nucleotide address and read depth to define clonal relationships. Finally, this analysis lends insight into potential cooperating drivers in keratinocyte transformation progression.

**Figure 3:**
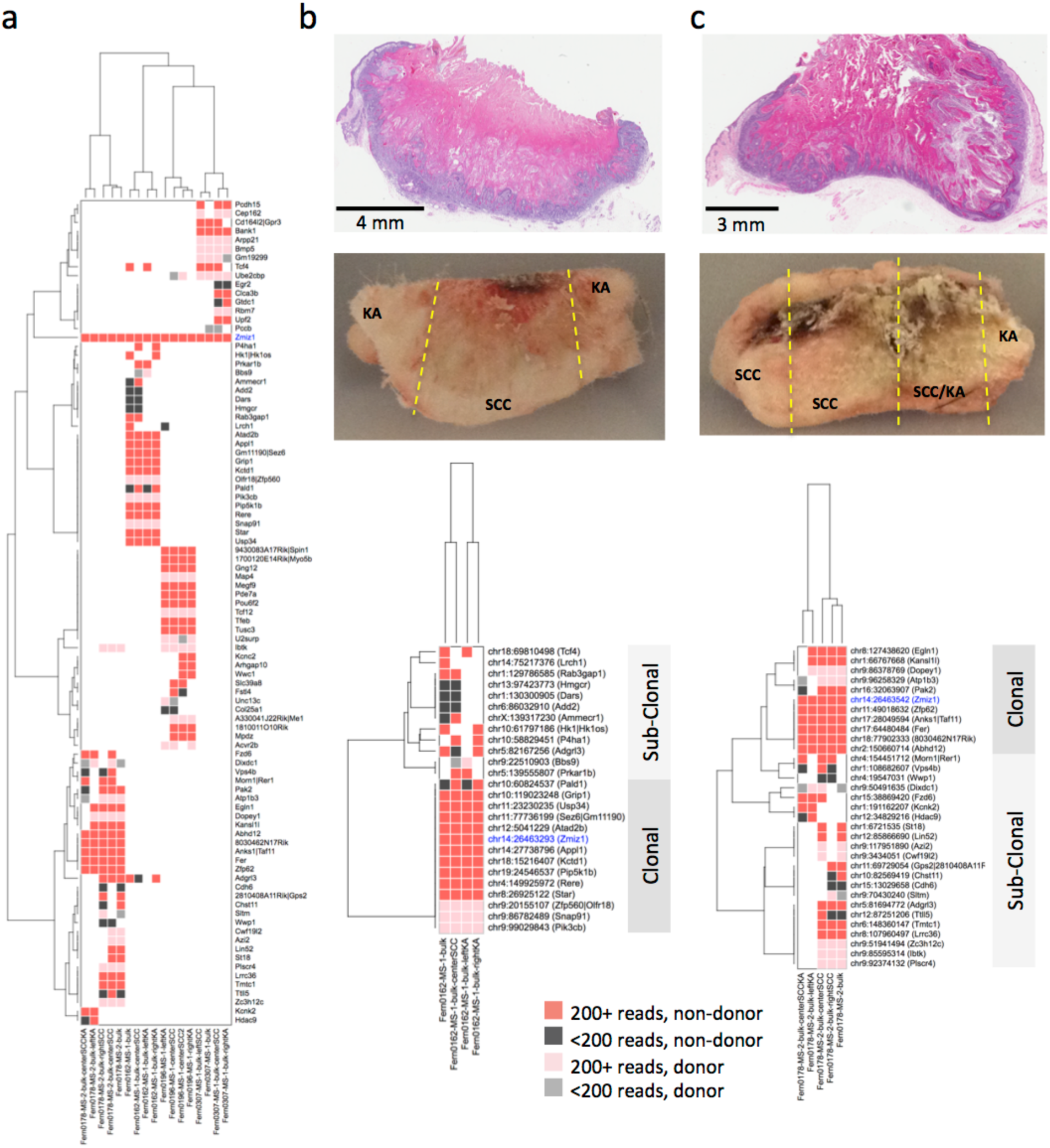
Multi-region SBCapSeq analysis of dual histology keratinocyte-derived skin masses containing distinct cuKA and cuSCC histological regions. (**a**) Hierarchical two-dimensional clustering (Hamming distance with the Ward method of agglomeration) of recurrent genic SBCapSeq insertion data from four skin masses containing distinct cuKA and cuSCC regions. Four distinct specimen (*x*-axis) and gene (*y*-axis) clades, each pertaining to a single bi-lesional mass, were detected demonstrating clonal identities. *Zmiz1* was the only gene recurrently mutated across all samples with high read depths. Genomic DNA was isolated from adjacent serial sections and included a “bulk” reference consisting of a cross-section of entire mass, and three or four regions dissected from histologically distinct regions (see **panels b-c** and **Supplementary Figure 8**). (**b**) Two masses with Aperio slide scans of H&E stained FFPE specimens showing distinct cuKA and cuSCC histologies suggestive of clonal evolution of cuSCC differentiation within cuKA masses and the adjacent flash frozen tissue prior to sectioning for genomic DNA isolation. Hierarchical two-dimensional clustering (Hamming distance with the Ward method) of recurrent genic SBCapSeq insertion data shown below each specimen demonstrates trunk (common to all samples) and sub-clonal, lesion-specific SB insertions.

#### Pten cooperation with Chuk in early-stage cuKA

In a previously published study, a surprising insight into genotype-phenotype relationships during keratinocyte transformation was found by comparing the sequencing of early-stage cuKA masses that arose spontaneously in a breast cancer cohort consisting of quadruple heterozygous mice carrying alleles for K5-Cre, Pten-floxed, Onc3(TG.12740), ^LSL^ SBase to phenotypically indistinguishable cuKA masses observed in the SB|P53 cohorts (Rangel et al., 2016; Stransky et al., 2011). In keratinocytes with SB mobilization and heterozygous loss of *Pten*, activating SB insertions into *Zmiz1, Zmiz2*, and *Mamld1* were rarely, if ever, observed. Instead, inactivating insertions into *Chuk* were statistically over-represented and, when they occurred, were mutually exclusive with *Zmiz1* or *Mamld1* insertions, suggesting they may be within the same genetic pathway. Surprisingly, SBCapSeq analysis of normal skin from 9 separate mice with the same quadruple heterozygous genotype revealed many low-read depth insertions into the *Pten* locus, but no insertions in *Chuk* (**Supplementary Table 21-22**), suggesting that selection for *Chuk* inactivation may be dependent upon haploinsufficient loss of *Pten*. In addition, using the datasets within the recently published SBCDDB (Newberg et al., 2017), we found statistically significant evidence for co-occurring loss of *Pten* (either via SB insertion or by breeding in a heterozygous null allele) and SB inactivation of *Chuk* across solid tumor models (Fisher’s exact test, *P*=2.36 × 10^−7^). Statistical significance for co-occurring mutations in *Pten* and *Chuk* in solid cancers persisted even after removing all cuKA tumors from the analysis (Fisher’s exact test, *P*=3.78 × 10^−16^), suggesting this relationship may be generally observed in non-keratinocyte tumor lineages. Remarkably, querying for co-occurrence of alterations in *PTEN* and *CHUK* via cBioPortal, across 41,834 patient samples from 170 studies, revealed significant co-occurrence for these two genes in human cancers (Chi-square with Yates correction, χ^2^=406, *P*<0.001; **Supplementary figure 4c-d**). Collectively, these data suggest that coordinated loss of both *Pten* and *Chuk* may have a role in initiating and progressing oncogenic transformation in keratinocyte as well as non-keratinocyte lineages. Given the recurring inactivating SB insertions in *Chuk* and activating *Zmiz1* observed in our *Pten*-loss sensitized mice (**Supplementary Figure 4a**), it is curious that a recent separate *Pten*-sensitized SB screen for cooperating TSGs failed to identify *Chuk* or *Zmiz1* entirely (de la Rosa et al., 2017). One possibility for this difference is that the reported screen used a single-copy SB allele, while our screen used multi-copy SB alleles with multiple independent genic insertion events per tumor, and the lack of *Chuk* or *Zmiz1* insertion events may suggest that positive selection for variant *Chuk* or *Zmiz1* alleles in haploinsufficient *Pten*-mutant skin cells may require one or more additional cooperating drivers.

### Comparative oncogenomic meta-analysis

One hundred-seven of the SB candidate Trunk driver genes are direct orthologs of human cancer genes found in the Cancer Gene Census (GRCh38 COSMICv86) (Futreal et al., 2004), a growing catalogue of mutations causally implicated in cancer, which is an enrichment greater than expected by chance (χ^2^=60.75 with Yates correction, *P*<0.0001; **Supplementary Tables 23–27**). Another 64 SB candidate driver genes, including *ZMIZ1, ZMIZ2*, and *KMT2C*, have human orthologs with recurrent non-silent mutations identified by exome sequencing (Lee et al., 2014; Pickering et al., 2014; South et al., 2014) of 68 cuSCC genomes (χ^2^=23.70 with Yates correction, *P*<0.0001). We conclude from these findings that SB identifies the same genes mutated by the diversity of mutational processes operating within end-stage human cuSCC genomes.

To gain insight into the biological functions of the candidate discovery drivers, we used a curated subset of the Enrichr analysis platform (Chen et al., 2013; Kuleshov et al., 2016) gene-set libraries for pathways and gene ontologies (see **Methods**) to perform biological pathway and process enrichment analysis from the cutaneous specimen cohorts. Here we integrated the drivers identified by 454 and SBCapSeq for cuSCC in SB|Onc3 mice and derived a composite list of cuKA drivers from two different cuKA datasets, one from cuKAs derived from SB|Onc3 mice sequenced by SBCapSeq and the second from cuKAs isolated from a previously published mouse model with K5-Cre; Pten^CKO/+^ and SB|Onc2 or SB|Onc3 mice (Rangel et al., 2016; Stransky et al., 2011) sequenced by 454. Using these integrated driver lists, we discovered statistically significant enrichment of recurrent SB candidate drivers in many signaling pathways and biological processes previously associated with human SCC, including EGFR, MAPK, NOTCH, and WNT-TGFβ (**Supplementary Figure 9a-i** and **Supplementary Table 28**).

Several of the top 30 most frequent drivers in SB cuSCC are key players in multiple pathways, suggesting that SB perturbation of multi-functional genes drives cuSCC. An oncoprint aligns the conserved drivers between SB cuSCC, SB cuKA and mutated human homologs in cuSCC that map to enriched pathways (**Fig.4**). Further, the SB|cuSCC Trunk drivers encode proteins with significantly more known protein-protein interactions than expected by chance (*P*=8.55 × 10^−6^, STRING enrichment analysis; number of nodes: 144, number of edges: 124, expected number of edges: 82) as do the SB|cuKA trunk drivers (*P*=6.93 × 10^−5^, STRING enrichment analysis; number of nodes: 37, number of edges: 14, expected number of edges: 4; **Supplementary Figure 10**). PPIs among drivers identified in SB-driven cuKAs suggests that perturbation of NOTCH signaling may be an initiating or acquired event in cuKA. Notably, we also observed that all SB|cuSCC tumor genomes have at least one defined driver involved in chromatin modification (consisting of chromatin and histone modification enzymes) and/or Hippo pathway genes (**Fig. 4** and **Supplementary Table 28**). Comparative genomic integration of the SB defined cuSCC and cuKA drivers with human cuSCC genomes reveals that while differences in the incidence of individual genes across cohorts can vary substantially, the biological pathways are conserved (**Supplementary Figure 9a**). These differences may be due in part to the types of alterations produced by SB transposition (gene activation/inactivation) versus UV-IR (silent/non-silent mutations) or, alternatively, may reflect the complexity of the systems of genetic networks (Nadeau and Dudley, 2011) that exists between driver genes and keratinocyte transformation phenotypes across species.

**Figure 4:**
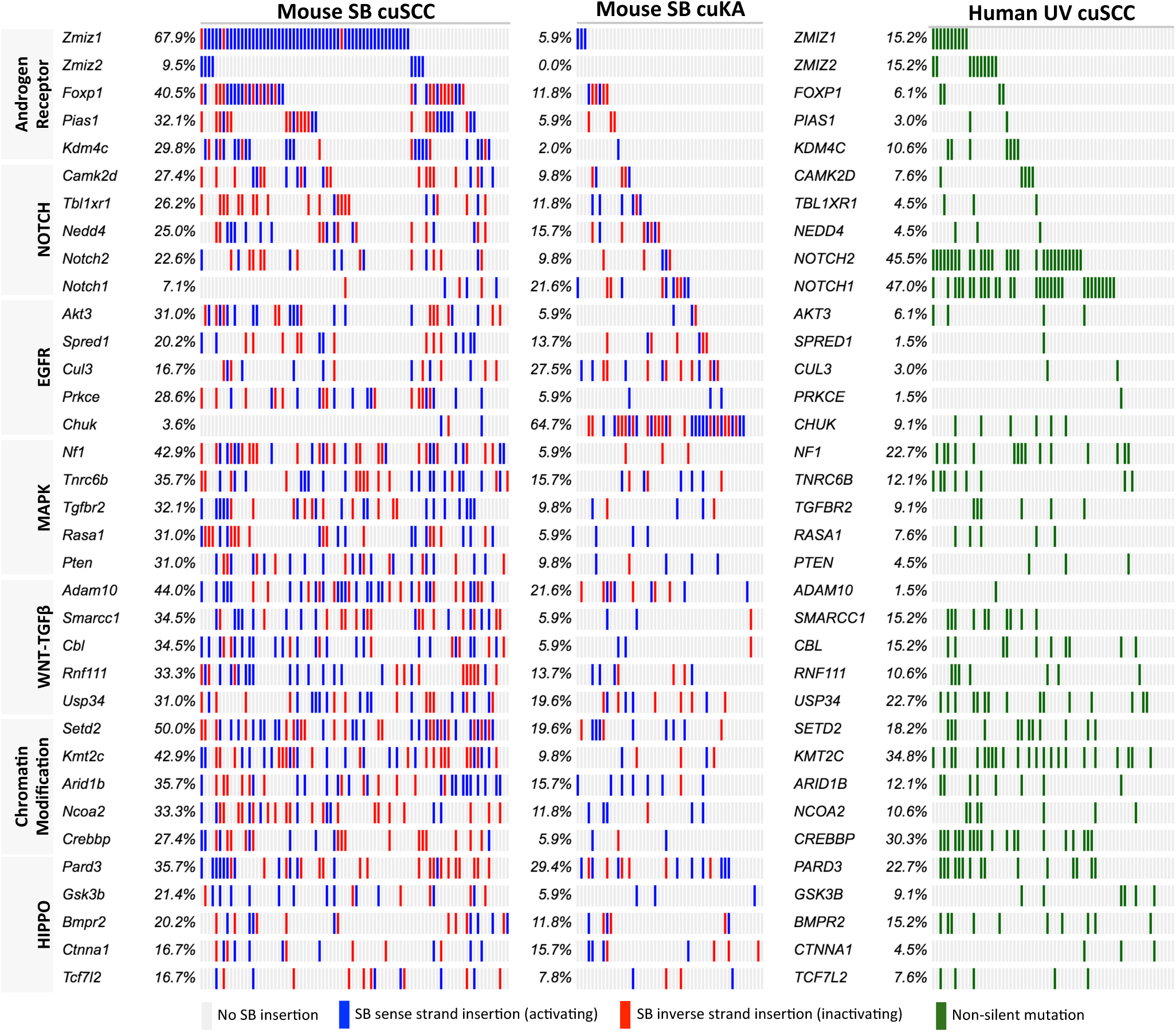
Comparative genomic landscape of candidate driver mutations and common altered pathways in mouse and human cuSCC genomes. Integrated Oncoprints sorted by biological pathway or process discovery significance within the SB|cuSCC dataset (**Supplementary Table 28**) and displaying the mutation burden across SB-induced mouse cuSCC (n=84) or cuKA (n=62) and human UV-IR-induced cuSCC genomes.

### SB-driven gene-expression alterations in cuSCC genomes

The most highly mutated oncogenic drivers in cuSCC genomes were the transcriptional co-factors *Zmiz1* and *Zmiz2*, which are collectively and mutually exclusively mutated in 71% of end-stage SB|cuSCC genomes. High read depth insertions are clustered into a single intron of each gene and occur in the same transcriptional orientation, where they are predicted to drive expression of a truncated chimeric transcript containing the oncogenic MIZ domain (**Fig. 5a**). We confirmed this prediction by RT-PCR (not shown) and normalized microarray data (**Fig. 5b**) analysis, where cuSCC tumors with *Zmiz1* insertions were found to have significantly higher mRNA abundance than tumors without *Zmiz1* insertions (**Fig. 5b**). We also observed a positive correlation between high-read-depth SB insertions among the 11 tumors harboring trunk *Zmiz1* insertions and *Zmiz1* gene expression (**Fig. 5c**).

**Figure 5:**
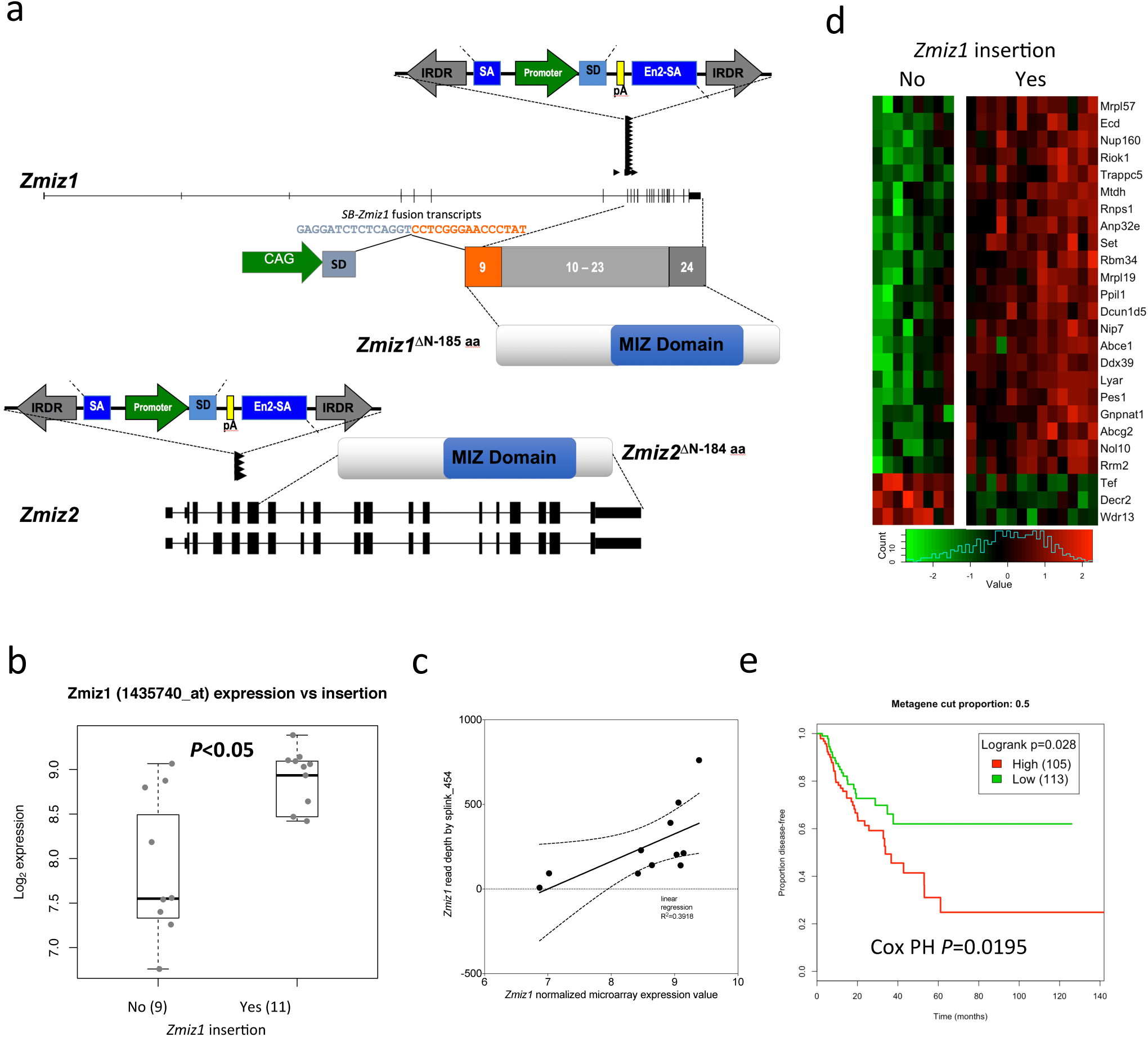
SB-mediated mutagenesis promotes N-terminal truncated ZMIZ paralog activation during cuSCC progression in mice. (**a**) Recurrent TA-dinucleotide SB insertion events from individual cuSCC genomes reveal activating sense strand SB insertions into Zmiz paralogs Zmiz1 and Zmiz2. Chimeric fusion between the SB transposon splice donor (SD) and splice acceptor sites at exon 9 of *Zmiz1* or exon 6 of *Zmiz2* resulting in a CAG-promoter driven transcription of a truncated mRNA that is predicted to encode N-terminally truncated proteins *Zmiz1*^ΔN-185 aa^ or *Zmiz1*^ΔN-184 aa^ that contain functional MIZ-type zinc finger domains, containing a Siz/PIAS RING finger (SP-RING). (**b**) Microarray gene expression analysis reveal that tumors with *Zmiz1* SB insertions (yes) had significantly higher expression of *Zmiz1*, compared to tumors without insertions (no). Box boundaries indicate interquartile range; whiskers indicate maximum and minimum values; center lines indicate medians. (**c**) High positive correlation between Splink_454 read depth and *Zmiz1* expression by microarray analysis. (**d**) Top 25 differentially expressed genes between genomes with (yes) or without (no) *Zmiz1* SB insertions (see **Supplementary Figure 11a**). (**e**) Survival plots for patients with head and neck type SCC (hnSCC) based on the expression of a *ZMIZ1*–centric metagene constructed using singular value decomposition of human hnSCC RNA-seq dataset from TCGA. These reveal poor patient survival outcomes predicted by increasing expression of *ZMIZ1* and closely regulated genes consisting of a 23-gene signature (**Supplementary Figures 14-16**).

Since ZMIZ1^ΔN185^ retains the DNA binding and transactivation domains, we reasoned that the *ZMIZ1*^ΔN185^ isoform may drive cuSCC via alterations in gene expression. To identify genes that were differentially expressed between SB cuSCC tumors with and without insertions in *Zmiz1*, we used Rstudio to create a Shiny app that used the normalized microarray data (see **Supplementary Table 29-30**), to generate a 25-(**Fig. 6d** and **Supplementary Fig. 11a**), 50-(**Supplementary Fig. 11b**), and 96-gene (**Supplementary Table 31**) differential expression signature (*P*<0.05, *P*_*FDR-corrected*_ <0.12) from the top differentially expressed genes. These genes demonstrated enrichment for biological pathways and processes related to gene expression, RNA processing and metabolism (**Supplementary Table 32-33**).

**Figure 6:**
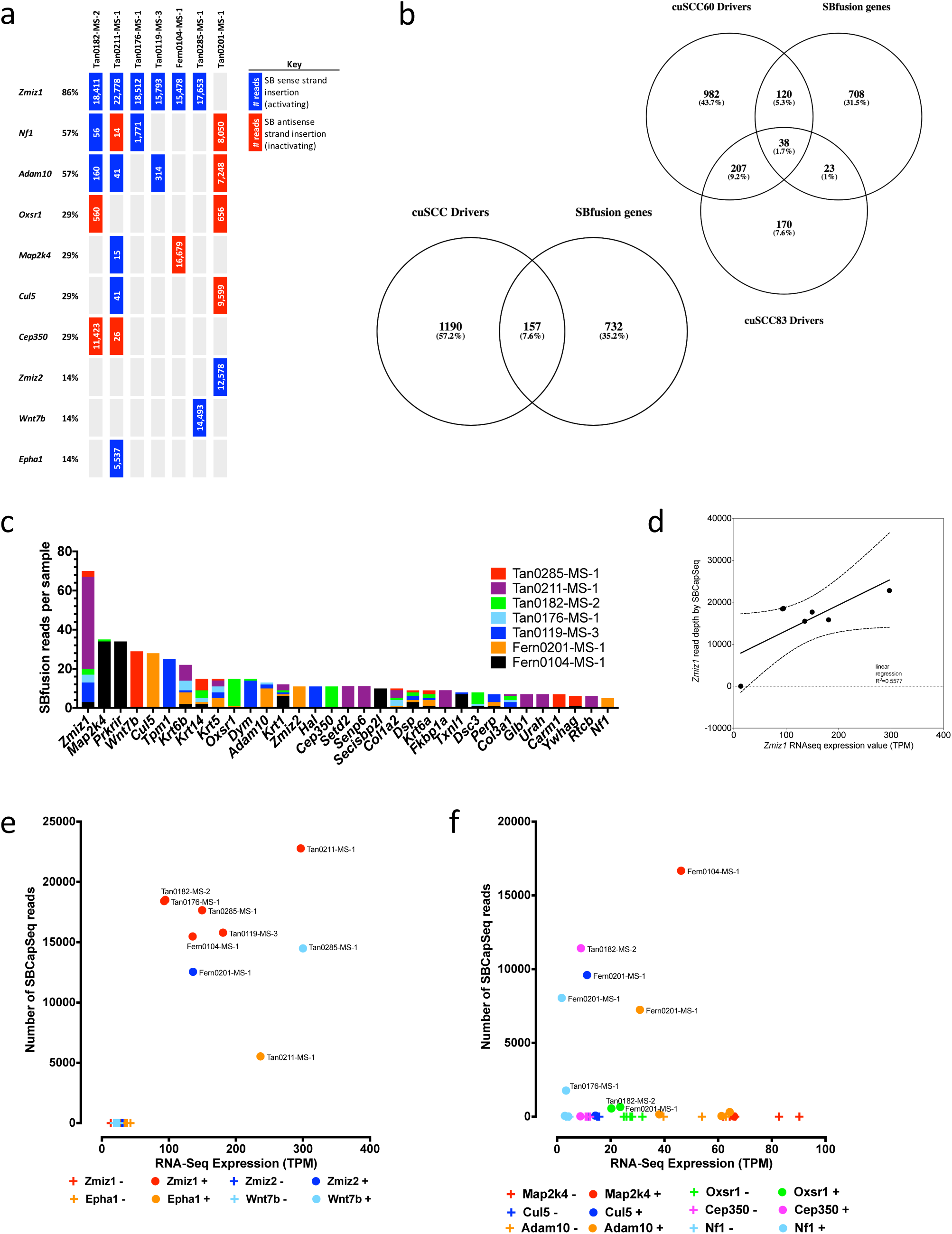
Clonally selected SB insertion events affect trunk driver gene expression in SB-induced cuSCC genomes. (**a**) Seven late-stage cuSCC SB-Onc3 genomes with mutually exclusive high-read depth *Zmiz1* or *Zmiz2* SB insertions selected for RNA-seq analysis of SB-fusion transcripts and global gene expression levels. (**b**) Venn diagrams comparing the overlap between cuSCC driver genes and detection of SBfusion transcripts by whole transcriptome RNA sequencing (wtRNA-seq) performed on bulk populations of cells isolated from 7 late-stage cuSCC genomes. (**c**) Genes with the most abundant SBfusion reads detected by wtRNA-seq analysis performed on bulk populations of cells isolated across the 7 late-stage cuSCC genomes. (**d**) High positive correlation between SBCapSeq read depth and *Zmiz1* expression by wtRNA-seq analysis. (**e**) Multifold induction of gene expression in cuSCC masses with (**+**) high read depth activating SB insertion events among 4 candidate oncogenic drivers compared with normal gene expression levels in cuSCC tumors without (**–**) SB insertions. Individual gene plots may be found in (**Supplementary Tables 39-40**). (**f**) Reduced gene expression in cuSCC masses with (**+**) high read depth inactivating SB insertion events among 6 candidate tumor suppressor drivers compared with normal gene expression levels in cuSCC tumors without (**–**) SB insertions. Individual gene plots may be found in (**Supplementary Tables 39-40**).

We also conducted a secondary differential analysis in R and identified 289 genes that were differentially expressed (*P*<0.05, *P*_*FDR-corrected*_ <0.2) between the *Zmiz1*-positive and *Zmiz1*-negative SB-driven cuSCC tumor groups (**Supplementary Table 34**). As expected, *Zmiz1* expression was significantly higher in tumors with activating insertions. Among the differentially expressed genes, the expression levels of *Serpinb2, Spink12*, and *Stfa3* were individually statistically significantly higher in cuSCC genomes with activating *Zmiz1* insertions than in cuSCC genomes without *Zmiz1* insertions (**Supplementary Fig. 11c**), and *Mtdh, Anp32e, Ecd, Dcun1d5*, and *Kdm7a* were also independently identified as cuSCC Progression Drivers. These statistically defined co-driver expression relationships indicate that these cooperating mutagenic insertions may drive keratinocyte transformation and/or progression in this mouse model of human cuSCC.

To identify potential oncogenic pathways and processes that were enriched among the 289 significantly differentially expressed genes (DEGs), we conducted enrichment analysis using Enrichr analysis platform. Based on the KEGG database, only the interleukin-17 signaling was significantly enriched among the DEGs, suggesting that *ZMIZ1*^ΔN185^ isoform may not alter oncogenic pathways at the transcript level. However, gene ontology analysis revealed a significant enrichment for biological processes involved in epidermis differentiation and development, suggesting that *ZMIZ1*^ΔN185^ isoform drives the dysregulation of keratinocyte development during cuSCC development. We observed enrichment of several paralog gene families, including keratin (*Krt1/10/15/16* and *Krtdap*), stratum corneum-associated structural late cornified envelope (*Lce1f* and *Lce3c*), and keratinocyte envelop cross-linking small proline rich (*Sprr1a* and *Sprr1b*) proteins, and key components of the epidermal differentiation complex (EDC) (Kypriotou et al., 2012), including *S100a8, S100a9, Ivl, Flg* (**Table 1**). Based on these finding, we asked whether these EDC genes are altered in human cuSCC. Using a published gene expression dataset (GSE32628 (Hameetman et al., 2013)), we demonstrate that EDC gene transcripts *IVL, LCE3C, S100A8, S100A9, SPRR1B*, and *SPRR2B* were significantly upregulated in human AK and cuSCC, relative to matched normal epidermis (**Supplementary Figure 12**). Importantly, increased expression of EDC genes has been reported in human cuSCC (Hudson et al., 2010). Taken together, our microarray gene expression analysis demonstrates that the *ZMIZ1*^ΔN185^ isoform is a critical mediator of gene expression changes essential for early keratinocyte transformation and sustained during cuSCC progression to end-stage tumors.

To further analyze the impact of clonally selected trunk SB insertions on gene-level and transcript-level expression, we sequenced ribo-depleted, whole-transcriptome RNA-seq (wtRNA-seq) libraries created from total RNA isolated from 7 cuSCC genomes (**Supplementary Table 35**) with high-read depth activating SB insertions into either *Zmiz1* (n=6) or *Zmiz2* (n=1) to identify SBfusion reads (Mann et al., 2016b). The wtRNA-seq reads contained spliced chimeric transposon splice-donor or splice-acceptor sequences fused to adjacent known exons in the mouse genome. In total, we identified 2,906 SBfusion wtRNA-seq reads from 889 unique RefSeq genes (**Supplementary Table 35**), including 154 progression and 27 trunk cuSCC drivers, respectively (**Supplementary Table 37-38**), which is substantially more than expected by chance (Chi-square test with Yates correction, χ^2^= 159, *P*<0.0001). Thirty-three genes with SBfusion events supported by ≥5 SBfusion reads, include our top trunk drivers *Zmiz1* and *Zmiz2* and several keratins (**Fig. 6a-c**). We also observed a positive correlation between high-read-depth SB insertions among the 6 cuSCC tumors harboring trunk *Zmiz1* insertions and exhibiting high *ZMIZ1* gene expression (**Fig. 6d**). We next selected 10 genes with high-read depth SBCapSeq insertion sites and SBfusion reads detected by RNA-seq within the 7 cuSCCs to investigate potential alterations in gene expression due to clonal SB insertion events, including 4 activating (*Zmiz1, Zmiz2, Wnt7b*, and *Epha1*) and 6 inactivating (*Nf1, Adam10, Oxsr1, Map2k4, Cul5*, and *Cep350*) SB insertion events (**Fig. 6e-f, Supplementary Figure 13a-f**, and **Supplementary Figure 14a-f**).

Importantly, in each tumor, SBfusion reads containing either *SB-Zmiz1* or *SB-Zmiz2* spliced-exons were represented in the top 2% of SBfusion transcripts and were supported by multiple independent SBfusion reads (**Supplementary Table 39-40**). *Zmiz1* or *Zmiz2* gene expression was significantly increased within each of the 7 cuSCC genomes (⪆100 and up to 300 times normal expression levels) and correlated with a highest SBCapSeq read depth sites with activating SB insertions and activating SBfusion reads (**Figure 6e** and **Supplementary Figure 13a-d**). This strongly suggests that the SB insertion events driving either *Zmiz1* or *Zmiz2* have been clonally selected to increase oncogenic expression of the N-terminal truncated forms during keratinocyte transformation and cuSCC progression *in vivo*. Similarly, *Epha1* and *Wnt7b* had high SBCapSeq read depth activating SB insertions and wtRNA-seq SBfusion reads and significantly higher gene expression compared to basal expression levels (⪆100 times) observed in tumors without insertions or SBfusion reads (**Figure 6e** and **Supplementary Figure 13e-f**). Since SB insertions into these genes were not recurrent at the population level, *Epha1* and *Wnt7b* were not identified as statistically significant drivers in cuSCC. However, the integrated genomic and transcriptomic analysis strongly suggests that each of these genes may have a ‘private’ role in driving tumor progression in the genomes in which they occur with elevated expression levels. Both genes encode proteins that are linked to cancer hallmarks — *Epha1* is an ephrin receptor subfamily of the protein-tyrosine kinase family (Potla et al., 2002) and *Wnt7b* is a WNT family member (Huguet et al., 1994). For the 6 predicted inactivated genes, we observed reduced gene expression that correlated with high SBCapSeq read depth of inactivating SB insertions and truncated SBfusion reads (**Figure 6f** and **Supplementary Figures 14a-f**). *Adam10, Cul5*, and *Nf1* are Trunk Drivers and *Map2k4* is a Progression Driver in cuSCC cohorts, further suggesting these events contribute to cuSCC progression *in vivo*. *Oxsr1* and *Cep350* were private selected events in individual tumors that may operate in cuSCC progression within the tumors that harbor clonal insertion events. Taken together, these data demonstrate that integrated genomic and transcriptome analyses may provide a means to discover significant cooperating drivers in cuSCC genomes from bulk specimens.

### Clinical relevance of SB|cuSCC driver orthologs in human SCC

The mammalian paralogous genes *Zmiz1* and *Zmiz2* are mutually exclusive trunk drivers activated in three quarters of SB-driven cuSCC genomes in an SB-driven cuSCCs, suggesting that *Zmiz* proteins are proto-oncogenes in cutaneous squamous keratinocyte transformation *in vivo*. *Zmiz1* and *Zmiz2* encode transcription factor proteins that share core c-terminal features, including androgen receptor (AR) binding domain, proline-rich binding domain, and MIZ-type zinc finger PIAS (protein inhibitor of activated STAT) domains, that coordinately act to both interact with and regulate the activity of various cancer-associated proteins (Beliakoff and Sun, 2006; Huang et al., 2005; Kahyo et al., 2001; Lee et al., 2013; Megidish et al., 2002; Schmidt and Muller, 2002, 2003; Sharma et al., 2003; Soler et al., 2008). SB activating trunk insertions in *Zmiz1* or *Zmiz2* are unique to SB-induced cuSCC tumors and are predicted to activate gene expression by driving expression of specific N-terminal truncated forms of ZMIZ1^ΔN185^ and ZMIZ2^ΔN184^, a closely related paralog (**Figure 5a**). The likelihood of achieving unique, sense-strand oriented SB insertion events into two related genes at the same protein coding region on two different chromosomes in dozens of tumors from independent mice by chance is incredibly small (*P*<0.0001), suggesting both trunk driver ZMIZ oncoproteins contribute directly to keratinocyte initiation and tumor progression *in vivo*. The ZMIZ1^ΔN185^ and ZMIZ2^ΔN184^ onco-proteins share 62% sequence identity, retain the zinc finger PIAS, proline-rich transactivation, and AR binding domains shared by both peptides. Importantly, both oncoproteins lack the N-terminus in the full-length protein, which is thought to inhibit the intrinsic transcriptional activity of a C-terminal proline-rich transactivation domain (Sharma et al., 2003). *ZMIZ1 (RAI17/ZIMP10)* and *ZMIZ2 (ZIMP7)* are collectively mutated or amplified in one third of human cuSCC (Pickering et al., 2014; South et al., 2014) and 5% of hnSCC (Cancer Genome Atlas, 2015) genomes. To investigate their oncogenic potential in human keratinocyte transformation, we performed a metagene expression analysis of *ZMIZ1* and *ZMIZ2* in human TCGA head and neck SCC (hnSCC) datasets for which gene expression (RNA-seq) and patient survival outcomes are available. A *ZMIZ1* metagene, with a correlation threshold of 0.65, identifies a 23-gene signature that links high expression with poor outcomes in human hnSCC patients (Cox Proportional Hazards Regression, *P*=0.0195; Log-rank Test, *P*=0.028 at 50% quintile; **Figure 5e; Supplementary Figure 15a–c**).

Ectopic over-expression of N-terminus truncated ZMIZ1 in mouse keratinocytes leads to epithelial hyper-proliferation but is not sufficient for progression to late-stage cuSCC (Rogers et al., 2013), suggesting that cooperating mutations are required for frank cuSCC progression *in vivo*. Biological pathway enrichment in the *SB* cuSCC genomes revealed several cancer-associated pathways, including the EGFR, NOTCH, and WNT signaling pathways. However, we noted that all *Zmiz*-mutated cuSCC genomes contained one or more inactivating SB mutations in chromatin-remodeling genes (**Supplementary Figure 9g**), suggesting this class of mutated tumor suppressor genes may contribute significantly in driving cuSCC progression. Four of the top-ranked chromatin remodeling trunk driver orthologs were also found to be significantly mutated in >15% of human cuSCC genomes (Pickering et al., 2014; South et al., 2014). Importantly, as observed in mouse tumors, mutations in the orthologs *KMT2C (MLL3), ARID1B (BAF250B), NCOA2 (KAT13C)*, and *CREBBP (CBP/KAT3A)* were also found in the same cuSCC genomes harboring *ZMIZ* mutations (**Figure 2**), suggesting they might cooperate during human keratinocyte transformation. Previously, *KMT2C* (*MLL3*) mutations were found to be associated with poor outcomes in aggressive cuSCC (Pickering et al., 2014), suggesting mutations in this gene may be a biomarker for advanced stage disease. To extend this result, we used survival information and RNA-seq expression from a hnSCC dataset, and observed that a *KMT2C* metagene, with a correlation threshold of 0.66, identifies a 176-gene signature that links high expression with poor outcomes in human hnSCC patients (Cox Proportional Hazards Regression, *P*=0.0498; Log-rank Test, *P*=0.0093 at 75% quintile; **Supplementary Figure 16**). Similarly, consistent with hippo signaling in cuSCC, we identified a 20-gene signature consisting of Hippo pathway orthologs predictive of patient outcome (Cox Proportional Hazards Regression, *P*<0.0001; Log-rank Test, *P*=0.0002 at 50% quintile; **Supplementary Figure 17**).

### Chromatin modifying tumor suppressors drive human keratinocyte initiation

Chromatin remodeling is one of the most frequently inactivated biological processes in SCC, and has been demonstrated to be involved in either tumor initiation or progression in multiple cancer types (Andricovich et al., 2018; Cho et al., 2018; Dhar et al., 2018; Jia et al., 2018; Mathur et al., 2017; Yu et al., 2016). In our SB cuSCC screen, chromatin remodeling was the top biological process that was significantly enriched (**Supplementary Table 28**). However, it remains unclear how expression alteration of chromatin remodeling genes feature in the cuSCC disease trajectory remains unclear. We prioritized three tumor suppressor genes involved in histone modification (*KMT2C* and *CREBBP*) or transcriptional regulation (*NCOA2*) with high concordance between SB-cuSCC and human cuSCC to study their roles in keratinocyte transformation using immortalized and minimally transformed HaCaT keratinocyte cell line (Boukamp et al., 1988). First, we confirmed expression of each gene within HaCaT cells (data not shown) and established a baseline colony formation assay using enforced expression of oncogenic KRAS-G12D in HaCaT cells (**Supplementary Figure 18**). Next, we used shRNAs targeting these genes to assess cellular transformation *in vitro* as a surrogate of tumor initiation, and disease progression *in vitro* and *in vivo* (**Fig. 7**, 1-factor ANOVA, *P*<0.05). To investigate whether knockdown of *KMT2C, CREBBP* or *NCOA2* can transform keratinocytes, we used HaCaT cells to stably knock down expression of target genes. Forced depletion of *CREBBP* significantly increased cellular proliferation relative to control, and a similar trend was observed with *KMT2C* knockdown (**Figure 7a**). No significant difference in proliferation was observed between *NCOA2* knockdown and control (**Figure 7a**). To more stringently investigate transformation, we assessed anchorage-independent growth, a hallmark of cellular transformation, using a soft agar colony formation assay. Knockdown of *CREBBP* or *KMT2C* expression resulted in significantly larger size and increased number of colonies, relative to control (**Figure 7b**), while no significant difference was observed between *NCOA2* knockdown and control (**Figure 7b**). Taken together, these data suggest that knockdown of *CREBBP* or *KMT2C* can transform the immortalized HaCaT cell system *in vitro*. Finally, we tested whether *KMT2C, CREBBP*, or *NCOA2* single gene knockdown in HaCaT cells can initiate the formation of masses *in vivo*. Cells were injected into flanks of immunodeficient *NSG* mice, and mass formation was monitored for 9 weeks. At endpoint, no significant difference in tumor volume was observed between control and *shKMT2C, shCREBBP* or *shNCOA2* (**Figure 7c-d**), suggesting that expression changes in more than one driver gene may be needed for *in vivo* transformation.

**Figure 7:**
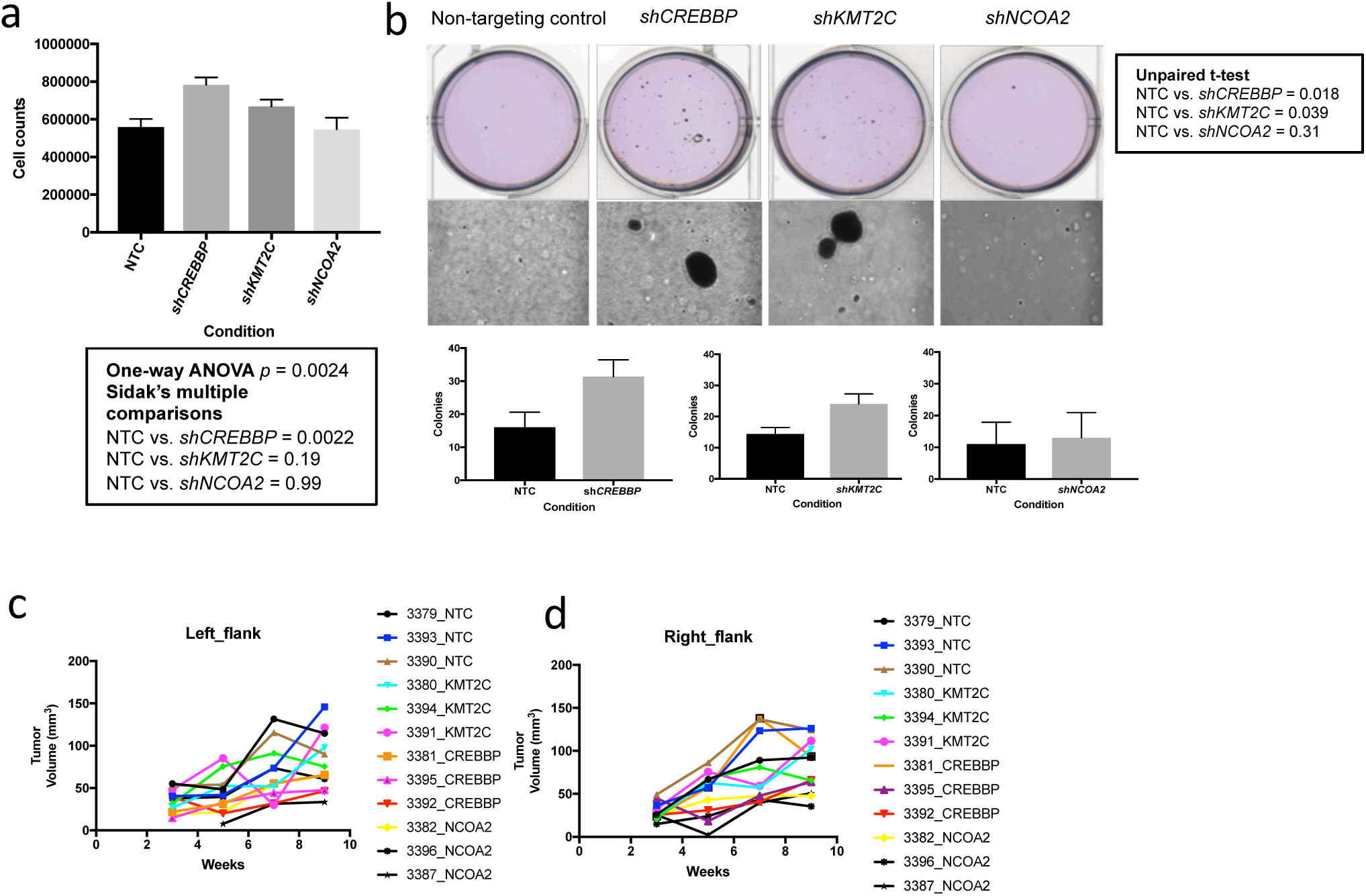
*In vitro* and *in vivo* functional validation of chromatin remodeler in immortal keratinocytes. (**a**-**b**) Assessment of keratinocyte transformation in response to knockdown of *CREBBP, KMT2C* and *NCOA2* by proliferation assay or anchorage-independent soft agar assay. For proliferation assay (n=6), statistical significance was tested by one-way ANOVA followed by Sidak’s multiple comparisons adjustment. For soft agar assay (n=3), statistical significance was tested by unpaired t-test. (**c-d**) In vivo assessment of mass formation in HaCaT cells with knockdown of KMT2C, CREBBP, or NCOA2. Three million cells were injected into left and right flanks of NSG immunodeficient mice (n=3 per condition), and growth monitored up to nine weeks.

### Functional validation of SB cutaneous candidate drivers in human SCC cell lines

To investigate the role of SB-driven cuSCC drivers to promote cuSCC progression and provide functional evidence that the SB candidate drivers are important for human SCC progression, we prioritized 12 top-ranked SB driver orthologs from cuKA or cuSCC datasets (*CHUK, CUL1, EHMT1, FBXO11, KMT2C, LARP4B, MAMLD1, NCOA2, NCOR2, NOTCH1, PTEN*, and *ZMIZ1*) for in vitro validation. Ten are predicted tumor suppressor drivers (*CHUK, CUL1, EHMT1, FBXO11, KMT2C, LARP4B, NCOA2, NCOR2, NOTCH1*, and *PTEN*) that are mutated in human cuSCC and commonly identified as drivers in other SB mutagenesis screens reported in the SBCDDB (Newberg et al., 2018c) and 2 predicted oncogene drivers, *ZMIZ1* and *MAMLD1*, that are mutated in human cuSCC but are not oncogenic drivers in other SB mouse models of human cancers (Newberg et al., 2018c). We performed short-term siRNA-mediated gene knockdown (3 siRNAs per gene, **Supplementary Table 41**) in two well-characterized SCC cell lines (A431 and SCC13) and one lung SCC cell line (SW900) that each harbor *TP53* mutations. Cell lines with confirmed knockdown of driver target gene expression were assayed for changes in proliferation rates. All 12 drivers tested had at least one siRNA that significantly altered cellular proliferation compared to a scrambled control siRNA in at least one cell line (one-factor ANOVA with multiple testing correction, FDR adjusted-*P*<0.1; **Supplementary Figure 19** and **Supplementary Table 42**). We observed a modest but significant decrease in cellular proliferation with ZMIZ1 depletion and a dramatic decrease in cell proliferation in A431 cells with ZMIZ2 depletion. Likewise, knockdown of the predicted tumor suppressive drivers consistently increased cell proliferation, supporting the role of these genes in human SCCs (**Supplementary Table 42**). *CHUK, NOTCH1*, and *PTEN* validated in all three cell lines. Increased proliferation in A431 cells, consistent with previously reported data (Leonard et al., 2011).

*NCOA2* validated in both cuSCC cell lines, and *FBXO11, KMT2C, LARP4B, MAMLD1* and *ZMIZ1* validated in one cuSCC and the lung cell line. *CUL1, EHMT1, NCOR2* were significant only in the SW900 lung cell line. si*PTEN* knockdown Additionally, 3 independent siRNAs targeting *ZMIZ2*, an independent driver and paralog of *ZMIZ1*, were tested and found to grow significantly slower compared to a scrambled control siRNA in the A431 cell line (**Supplementary Figure 19**).

To both confirm and extend these findings, we next stably depleted each of these 12 drivers in in A431 cells using lentiviral pools containing three non-overlapping, short hairpin RNAs (shRNAs) to individually knockdown each of the 13 target candidate drivers and one non-targeting control delivered to silence target gene expression in A431 cells (**Supplementary Figure 20**) which was verified by qPCR (**Figure 8a**). Seven of the drivers displayed statistically significant changes in proliferation of A431 cells: *shZMIZ1, shZMIZ2*, and *shMAMLD1* decreased proliferation; and *shKMTC2, shNCOA2, shFBXO11* and *shCHUK*, accelerated proliferation, consistent with their predicted roles as proto-oncogenes, while shRNA-mediated knockdown of tumor suppressors KMTC2, NCOA2, FBXO11 and CHUK, accelerated proliferation (**Figure 8a-c**, two-factor ANOVA with multiple comparisons, FDR-adjusted-*P*<0.05). While knockdown of the *ZMIZ* paralogs in the A431 cell line demonstrated significant reduction in proliferation, we sought to verify that this observation was not cell-line specific. Using the COLO16 cell line, we achieved stable knockdown of *ZMIZ* expression, and observed decreased proliferation, relative to control. Taken together, our functional shRNA screen has yielded strong evidence for the *ZMIZ* oncogenes identified from our SB screen to be involved in human cuSCC progression *in vitro*, and prioritized several tumor suppressor genes for further characterization.

**Figure 8:**
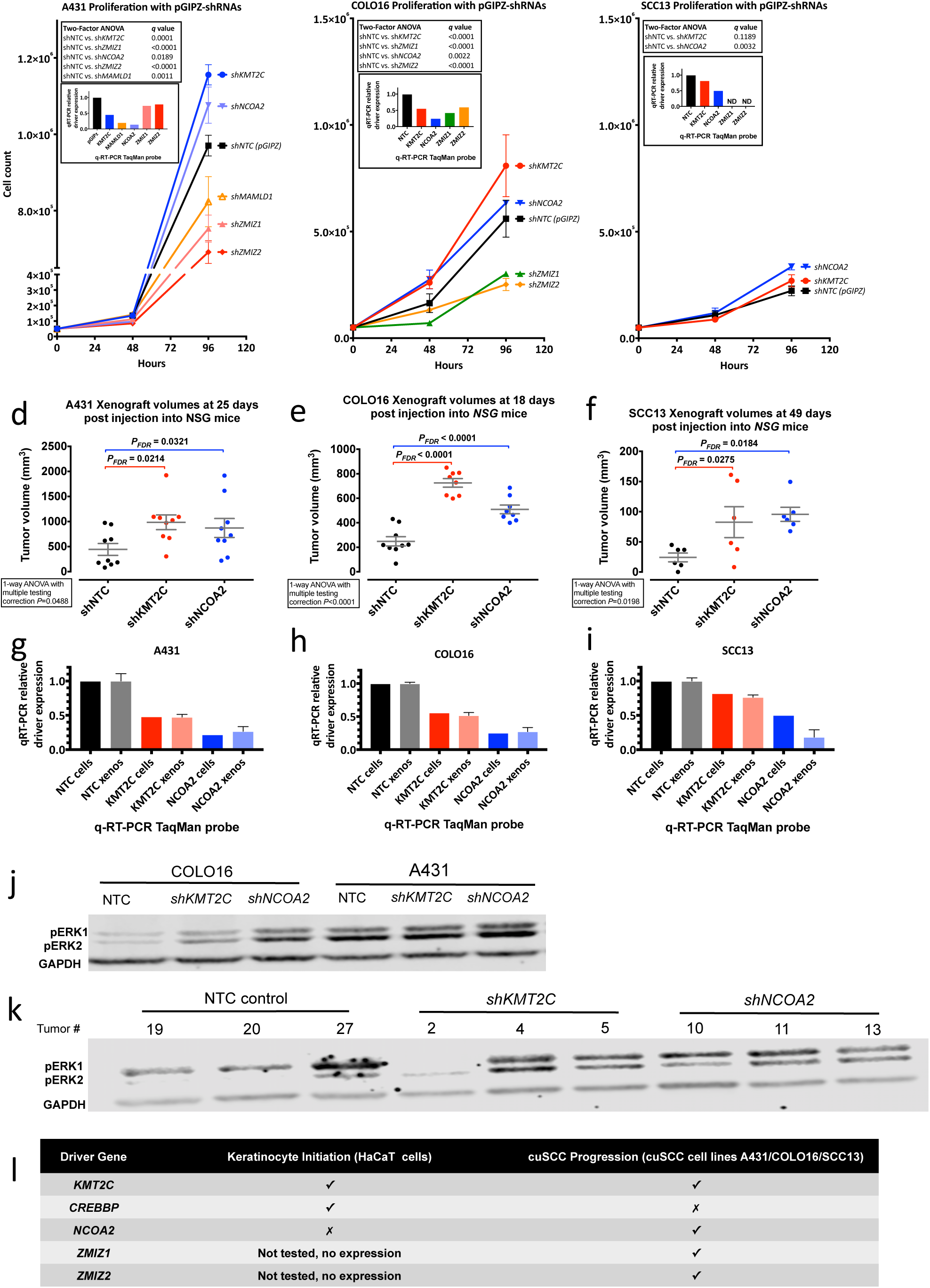
*In vitro* and *in vivo* functional validation of chromatin remodeler and ZMIZ drivers in cuSCC cell lines. (**a**-**c**) Knockdown of *KMT2C, NCOA2* or *ZMIZ* genes alter proliferation rates of human cuSCC cell lines (n=3; error bars, SEM; statistical significance measured by one-way ANOVA followed by multiple pairwise comparisons and FDR-adjusted *P-*values). (**d-f**) *KMT2C* and *NCOA2* knockdown accelerates cuSCC xenograft progression *in vivo*. (**g-h**) qRT-PCR values in ‘cells’ at the time of injection into NSG mice and in ‘xenos’ from specimens collected at necropsy. N=1 for ‘cells’ and N=6 or 8 (duplicates from 3 (COLO16) or 4 (A431, SCC13) independent masses) from xeno specimens. (**j**) Immunoblot for phospho-ERK expression in COLO16 and A431 cell lines with *KMT2C* or *NCOA2* knockdown. (**k**) Immunoblot for phospho-ERK expression from three largest tumors for each condition from the COLO16 xenograft assay shown in (**d**). (**l**) Summary of genes investigated with regards to their knockdown effects in human immortal keratinocyte and cuSCC cell line phenotypes.

### Chromatin modifying tumor suppressors drive human keratinocyte initiation and cuSCC progression

Since we demonstrated that knockdown of *KMT2C* or *NCOA2* increases the proliferation rates of A431 cell line *in vitro*, we asked whether this finding could be extended to other cell lines. Consistent with the observation made in A431 cells, we observed increased proliferation in COLO16 and SCC13 cells with decreased expression of *KMT2C or NCOA2*, relative to control (**Figure 8**, *P*<0.05, 1-factor ANOVA). We then determined whether *KMT2C and NCOA2* are cuSCC TSGs *in vivo*, and performed xenograft assays by subcutaneous injection of cells for each of the three cuSCC cell lines into *NSG* immunodeficient mice. All three cuSCC cell lines demonstrated enhanced *in vivo* tumor growth with *KMT2C, NCOA2*, or *CREBBP* depletion compared to cells with a non-targeting control shRNA (**Figure 8d-i**, 1-factor ANOVA, *q*<0.05 and **Supplementary Figure 21-22**), providing functional evidence that *KMT2C and NCOA2*, but not *CREBBP*, are human TSGs that promote *in vivo* cuSCC progression.

Finally, we explored the mechanistic basis of how decreased expression of *KMT2C* or *NCOA2* may progress cuSCC. The MAPK pathway has been shown to be operational in cuSCC (Lambert et al., 2014), and melanoma patients receiving BRAF inhibitor can develop cuSCC via hyperactivation of the MAPK pathway (Zhang et al., 2015). Since *KMT2C* and *NCOA2* can dynamically regulate global changes in gene expression, we hypothesized that knockdown of these two genes progress cuSCC via altering the MAPK flux. To test this, we immunoblotted for total and phosphorylated ERK (phospho-ERK) expression in our stable cell systems. In A431 and SCC13 cell lines with high basal phospho-ERK expression, knockdown of *KMT2C* or *NCOA2* did not alter phospho-ERK levels. In contrast, in COLO16 cell line with low phospho-ERK expression in control, knockdown of either *KMT2C* or *NCOA2* resulted increased ERK activation (**Figure 8j**). We pursued this finding further and used the flash-frozen tissues from our xenograft assay. Whole cell lysates from the three largest tumors for each condition (i.e., NTC, *shKMT2C, shNCOA2*) were harvested, and probed for phospho-ERK expression (**Figure 8k**). Consistent with our *in vitro* data, we observed a trend of increased phospho-ERK expression in tissues with *KMT2C* or *NCOA2* knockdown. Taken together, these data suggest that increased MAPK signaling may be one mechanism through which *KMT2C* or *NCOA2* knockdown drives to cuSCC progression.

## DISCUSSION

In this study, we sought to define the cooperating genetic events required for keratinocyte transformation and progression to frank cuSCC using the *SB* transposon mutagenesis system *in vivo*. Using an *Actb-Cre* transgene to activate systemic expression of the transposon, we observed tumor lesions in multiple organs. The kinetics of tumorigenesis were further accelerated in mice harboring either null (*Trp53*^*KO/+*^) (Jonkers et al., 2001) or recurrent point mutant (*Trp53*^*R172H/+*^) (Olive et al., 2004) alleles. This observation is consistent with previous studies showing that missense mutations in *Trp53* promote malignant transformation by abrogating both the tumor suppressive and oncogenic (gain-of-function and/or dominant negative) functions of Trp53 *in vivo* (Doyle et al., 2010; Hanel et al., 2013; Lang et al., 2004; Morton et al., 2010; Olive et al., 2004). While the SB|Onc3 mice developed a spectrum of tumor types, including of the hematopoietic lineage, we observed an enrichment for late-stage, well-differentiated cuSCC.

Despite the high curative rate for cuSCC, there is a significant population of patients who develop advanced disease, for which there are no defined therapeutic targets. Due to the high UV-induced mutation burden, combined with the scarcity of clinical samples, it is challenging to define the recurrent and druggable targets within these tumors. To address this need, we isolated early and advanced lesions from a SB mouse model of cuSCC to define recurrent drivers and potential new avenues for therapeutic intervention. From our in-depth sequencing analysis of SB insertions in advanced cuSCC genomes, we identified clonal and sub clonal drivers involved in the progression and maintenance of cuSCC. 64 drivers have human orthologs with non-silent mutations in human cuSCC genomes. Intriguingly, the majority of drivers in SB-driven cuSCC lesions were predicted tumor suppressor genes, suggesting that cumulative loss of these genes is a central feature of cuSCC. We showed that a few key TSGs, including *KMT2C* are differentially expressed in cuSCC with prognostic implications for patients with hnSCC.

We subsequently focused on understanding the genetic events required for keratinocyte transformation. Our SBCapSeq and SB Driver analysis identified 349 CCDs, regardless of *Trp53* status, of which 107 have been implicated in human cancers. Intriguingly, we observed that the SB-driven cuSCC lesions were driven almost exclusively by SB inactivating insertions in tumor suppressor genes, suggesting that cumulative loss of these TSG drivers is a central feature of cuSCC. Indeed, recent analysis from our lab, using a complimentary promoterless SB transposon forward genetic screen, revealed that tumor suppressor drivers alone can lead to keratinocyte initiation and cuSCC progression *in vivo* (Aiderus et al., 2020). Furthermore, in cuSCC lesions that arose from K5-Cre; *Pten*^*CKO/+*^ model previously reported, our SB Driver Analysis workflow revealed inactivating insertion patterns in all 300 CCDs, which corroborates the Actin-Cre driven model utilized in our study. Notably, in the single allele *Pten* loss model, we identified inactivation in *Chuk* (also known as *Ikka*) as a significant driver of cuSCC, and this gene has previously been reported to have a tumor suppressive role in cuSCC via regulating the EGFR signaling axis (Liu et al., 2008). However, since *Chuk* inactivation only occurs in a haploinsufficient *Pten* background, these data suggest that inactivation of these genes alters different pathways that cooperate to drive cuSCC. Tantalizingly, PanCancer analysis of over 40,000 tumors from multiple tissue types using cBioPortal revealed a significant co-occurrence of both genes, suggesting that the association between *Pten* and *Chuk* may be extended to cancer derived from different cellular lineages. Our data provide a rich resource to rationally identify, prioritize, and credential likely cooperating mutational events in cuSCC (**Fig. 3**).

Our report is the first to identify *Zmiz2* as a significant candidate cancer gene in any SB study and the first to discover mutual exclusivity among activated *Zmiz1, Zmiz2*, and *Mamld1* in early keratinocyte transformation and cuSCC progression. Insertions into *Zmiz1* (Dupuy et al., 2009; Friedel et al., 2013; Rogers et al., 2013) and *Mamld1* (Friedel et al., 2013) have been previously observed in skin tumors induced by transposon insertional mutagenesis. The possible redundancy of ZMIZ oncoproteins in the keratinocyte transformation program, by mutual exclusivity of trunk insertions within cuSCC genomes, strongly suggests that the ZMIZ1^ΔN185^ and ZMIZ2^ΔN184^ oncoproteins may have common neomorphic properties that contribute to the hallmarks of skin cancer. The work of Rogers *et al*. (Rogers et al., 2013), and more recently Mathios *et al*. (Mathios et al., 2019), on the dynamic regulation of *ZMIZ1* expression in human cancer, together with our finding that *ZMIZ1* and *ZMIZ2* are mutated with at least on non-silent alteration in at least one third of published human cuSCC genomes (Pickering et al., 2014; South et al., 2014) and, suggest that this gene may be an important initiating trunk mutation in human keratinocyte initiation and cuSCC progression. Transcriptome analysis on SB-driven cuSCCs with high read depth insertions in *Zmiz1* or *Zmiz2* to detected SB transposon fusions to both drivers. Of note, the N-terminal truncated versions of both *Zmiz* genes were selected in a mutually exclusive manner. The Zmiz1^ΔN185^ and Zmiz2^ΔN184^ onco-proteins share 62% sequence identity, and share the zinc finger PIAS, proline-rich transactivation, and AR binding domains. Importantly, both oncoproteins lack the N-terminus in the full-length protein, which is thought to inhibit the intrinsic transcriptional activity C-terminal proline-rich transactivation domain (Sharma et al., 2003). *ZMIZ1* and *ZMIZ2* are mutated in one third of human cuSCC genomes sequenced to date (Pickering et al., 2014; South et al., 2014), however, its functional significance remains unclear. *Zmiz1* has been shown to be involved in T cell development and leukemogenesis via interacting with NOTCH1 (Pinnell et al., 2015), but how it is involved in the tumorigenesis of other cell lineages is unclear. We have provided two preliminary lines of evidence with regards to how *ZMIZ* may be involved in squamous cancers. First, shRNA-mediated knockdown of *ZMIZ1* and *ZMIZ2* in A431 and COLO16 human cuSCC cell lines significantly decreased their growth rates, compared to non-targeting controls. While ZMIZ1 overexpression has been shown to sufficient for disease initiation, our cell proliferation data suggests that pathways regulated by ZMIZ1/2 continue to be active in advanced disease. Further, the association of high expression of a *ZMIZ*-centric metagene derived from TCGA hnSCC cohort, with poor patient survival supports an oncogenic role for *ZMIZ* in tumors of squamous lineage.

We identified potential signaling pathways, such as EGFR, NOTCH and WNT, that may cooperate with ZMIZ to drive cuSCC *in vivo*. Intriguingly, all SB cuSCC tumors with *Zmiz1/2* insertions had inactivating insertions in at least one gene involved in chromatin remodeling, which suggests that changes in gene expression via epigenetic regulation are necessary to progress a benign lesion to frank cuSCC. Some examples of the chromatin remodeler genes that were recurrently altered in our screen include members of the complex of proteins associated with Set1 (COMPASS) such as *Kmt2c* and *Kdm6a*, and the switch/sucrose non-fermenting (SWI/SNF) such as *Arid1b, Arid1a* and *Pbrm1*. Although the COMPASS and SWI/SNF pathway members have been reported to be mutated in various cancers (Gozdecka et al., 2018; Kadoch et al., 2013; Wang et al., 2018), whether alterations in one or cooperation of both pathways is required for the etiology of the disease remains unclear. Our functional data showing that knockdown of *KMT2C*, a key molecule in the COMPASS complex, in human cuSCC cell lines accelerated *in vitro* proliferation and *in vivo* xenograft growth, supporting a role for this gene as a tumor suppressor in advanced disease. Of clinical significance, several recent studies have highlighted potential dependencies amenable to pharmacologic inhibition in cancer cells with aberrations in COMPASS members (Wang et al., 2018).

While mutations in chromatin remodelers have been reported in the literature, only recently have studies provided functional evidence that this family of genes can function as tumor suppressor in the initiation and progression of different cancers. Our *in vivo* screen identified inactivation of at least one gene involved in chromatin remodeling in every tumor, suggesting that this family of genes may have an important role at some stage during cuSCC genesis. Our functional data suggests that *CREBBP* is involved in the earlier stages of keratinocyte transformation, while decreased *NCOA2* expression can drive disease progression. Interestingly, *KMT2C* knockdown drives keratinocyte transformation or cuSCC progression in different cell systems utilized. Finally, while ongoing work is focused on elucidating the mechanistic basis of how alterations in chromatin remodeler gene expression affect cuSCC biology, we demonstrate that one route of cuSCC progression via decreased *NCOA2* or *KMT2C* expression is activation of the MAPK pathway. Notably, this was observed in one of three cell lines assessed, suggesting that depending on the genetic background, chromatin remodelers influence disease progression via different mechanisms. In a similar genetic screen for myeloid leukemia drivers (Mann et al., 2016b; Newberg et al., 2018a), we identified *Ncoa2* as an oncogene and *Zmiz1* as a tumor suppressor, in contrast to the tumor suppressor of *Ncoa2* and oncogenic *Zmiz1* in the cuSCC model described in this study. These two examples, and several other from our various studies cataloged in the SBCDDB (Newberg et al., 2018c), support the dual roles many candidate cancer driver genes have in both promoting and inhibiting various cancer hallmark features depending upon the complexities of network of interactions operating to control genotype-phenotype outcomes in disease. In a related example, it was surprising that the frequency of *Zmiz1/2* insertions were low within the K5-cre|Pten cuKA masses given that our study in SB|Onc3 mice, and several previous transposon studies, have found activating *Zmiz1* insertions with high frequency in skin tumors. This may result from the use of the *Krt5*-cre, as the *Keratin5* promoter is principally active within basal skin cells but not keratinocytes. This raises the possibility that the tumor cell of origin is another important factor for skin cancer progression (as it is in breast cancer), and suggests that masses arising from basal cells can readily form frank cuKAs but not cuSCCs for unknown reasons. Additional experiments are needed and warranted to explore this possibility.

Our experimental approach for driving high mutational burdens by SB mutagenesis, recapitulates the sporadic, stepwise evolutionary selection of cooperating drivers in non-melanoma skin cancers in humans exposed to solar UV. Despite these divergent mutagens, we observed remarkable overlap of driver genes, pathways, and networks between the mouse and human cuSCCs, suggesting a deep mutual biology between these shared drivers and cutaneous oncogenesis. We hope that by defining the cooperative oncogenic and tumor suppressor networks that operate during keratinocyte transformation and subsequent cuSCC progression, the results of our screen will provide a foundation for exploring new therapeutic strategies.

## Author Contributions

M.B.M., K.M.M. N.G.C., and N.A.J. conceived the project and provided laboratory resources and personnel for animal husbandry, specimen archiving, sequencing and compute management. M.B.M. and K.M.M. designed the studies, directed the research, interpreted the data, performed experimental work, designed the SBCapSeq oligos, coordinated sequencing efforts and analyzed the data. A.A., L.G.-R., A.M.C.-S., A.L.M., D.J.J., F.A.-M., R.R., S.-C.L., K.H.-K.B., and K.Y.T. provided reagents, performed and/or analyzed experimental datasets. A.A., J.Y.N., M.A.B., and M.B.M. performed statistical and bioinformatics analyses. K.R. and S.M.R. performed and directed necropsy and histotechnology services. J.M.W. performed and directed veterinary pathology analysis, including tumor grading and diagnosis. L.M. and L.S. isolated RNA and performed microarray hybridizations. L.G.-R., A.M.C.-S., D.J.J., and F.A.-M. performed and optimized capture hybridizations for the SBCapSeq method and/or performed Ion Torrent sequencing for SBCapSeq, RNA-Seq experiments. M.B.M., L.G.-R., A.A., and A.L.M performed *in vitro* experiments. M.B.M. and A.L.M performed *in vivo* xenograft experiments. A.A., K.M.M., and M.B.M. wrote the manuscript with contributions from N.G.C., N.A.J., K.Y.T., and M.A.B. All co-authors approved the manuscript prior to submission.

## Competing financial interests

The authors declare no competing financial interests.

**Supplementary Figure 1:**
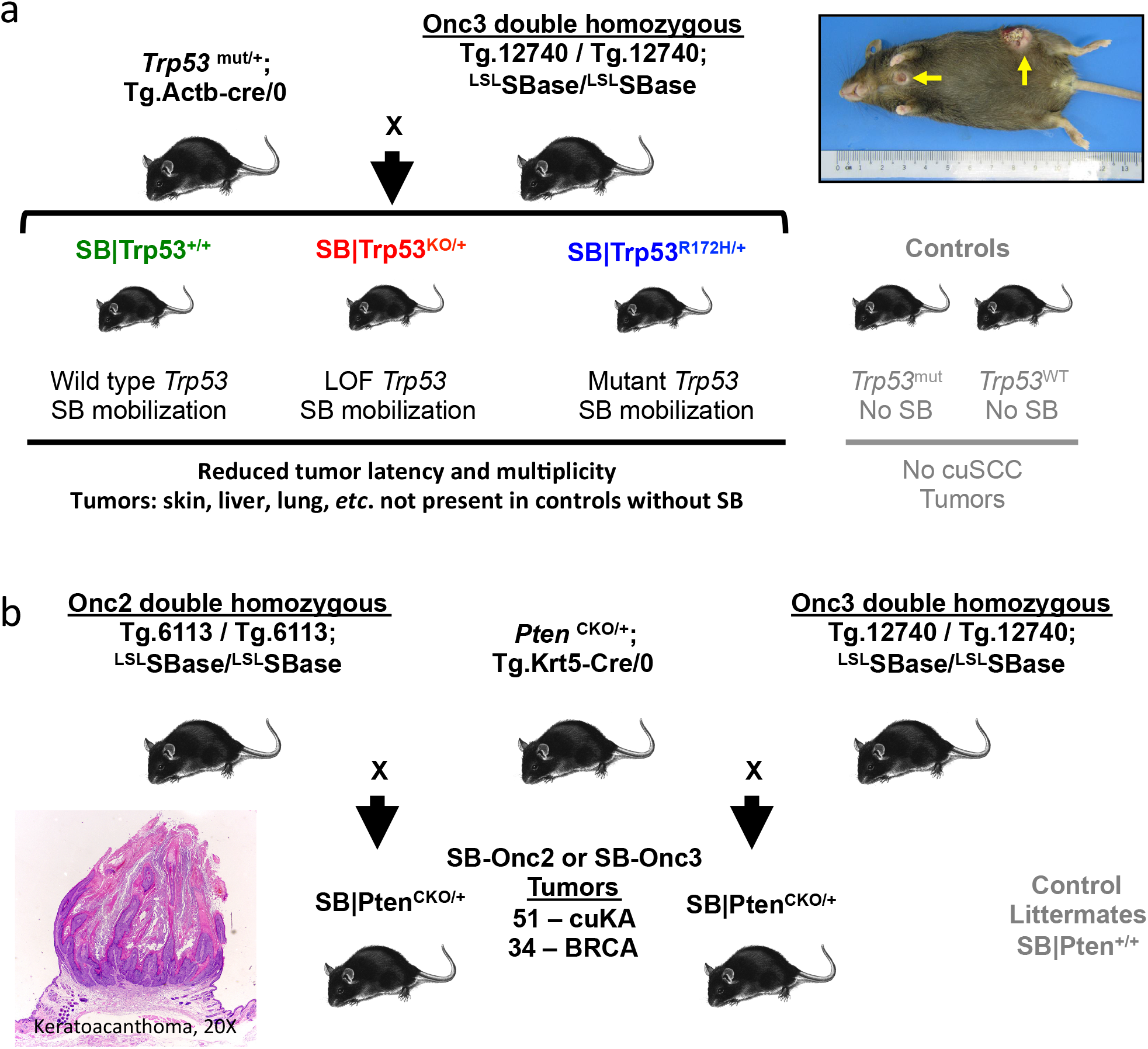
Overview of SB-induced tumor studies. (**a**) SB|Trp53|Onc3, (**b**) SB|Pten|Onc2, and (**c**) SB|Pten|Onc3 cohorts, from (Rangel et al., 2016), used in this work. Yellow arrow indicates cuSCC masses at necropsy.

**Supplementary Figure 2:**
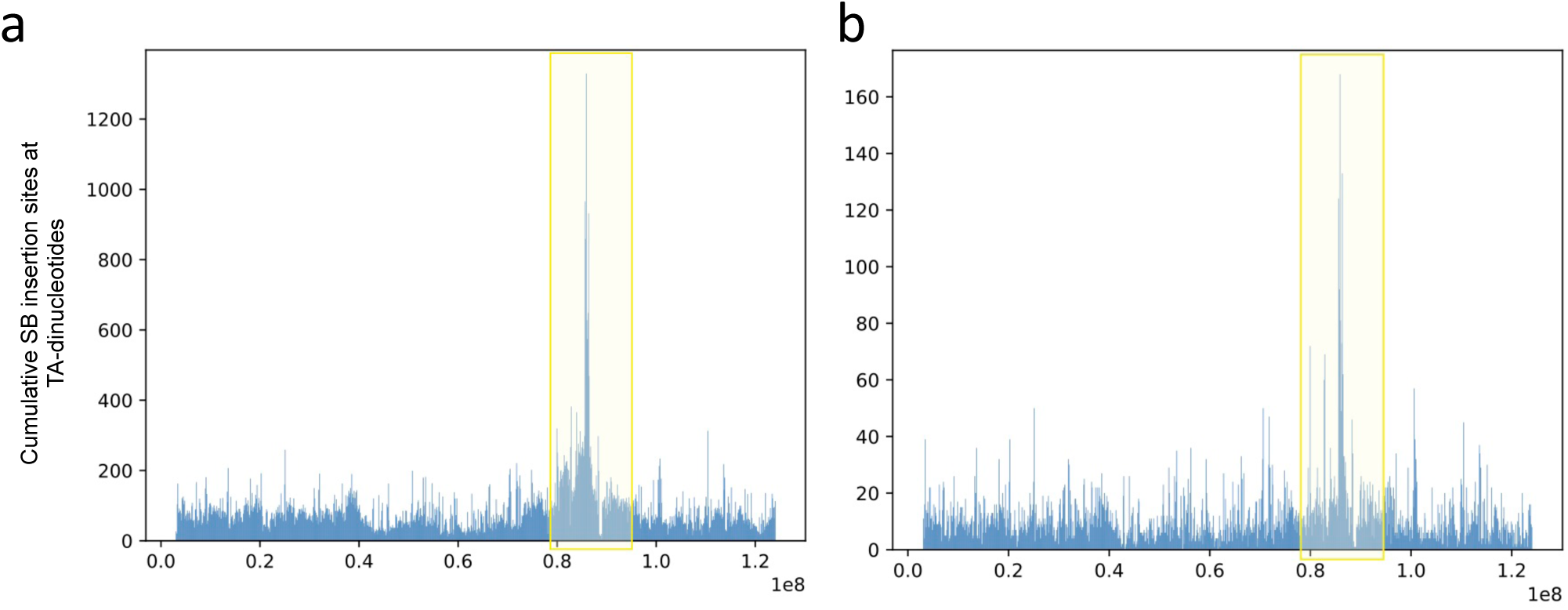
SB T2/Onc3 TG. 12740 allele donor position mapping and exclusion for SB Driver Analysis. Cumulative SB insertion sites at TA-dinucleotides from (a) 454_Splink and (b) Ion_SBCapSeq datasets showing the likely TG.12740 donor site at 87,000,000 bp and the exclusion region (yellow box) between 79,000,000 and 95,000,000 bp applied when SB Driver Analysis was run on chromosome 9. The 81 genes between *Filip1* at chr9:79663368-79825689 and *Tfdp2* at chr9:96096693-96224065 were excluded from SB Driver Analysis. The complete list of 81 genes excluded from SB Driver Analysis from tumors harboring the SB T2/Onc3 TG.12740 donor allele: *1190002N15Rik, 1700034K08Rik, 1700057G04Rik, 1700065D16Rik, 4930524O08Rik, 4930554C24Rik, 4933400C23Rik, 9330159M07Rik, 9430037G07Rik, A330041J22Rik, Adamts7, AF529169, Ankrd34c, Atr, B430319G15Rik, Bckdhb, Bcl2a1a, Bcl2a1b, Bcl2a1d, Cep162, Chst2, Ctsh, Cyb5r4, D430036J16Rik, Dopey1, Elovl4, Fam46a, Gk5, Hmgn3, Htr1b, Ibtk, Impg1, Irak1bp1, Lca5, Me1, Mei4, Mir184, Mir6386, Mir7656, Morf4l1, Mrap2, Mthfs, Mthfsl, Myo6, Nt5e, Paqr9, Pcolce2, Pgm3, Phip, Plod2, Pls1, Plscr1, Plscr2, Plscr4, Plscr5, Prss35, Rasgrf1, Ripply2, Rwdd2a, Senp6, Sh3bgrl2, Slc9a9, Snap91, Snhg5, Snx14, Syncrip, Tbc1d2b, Tbx18, Tmed3, Tpbg, Trim43a, Trim43b, Trim43c, Trpc1, Ttk, U2surp, Ube2cbp, Xrn1, Zfp949, Zic1, Zic4*.

**Supplementary Figure 3:**
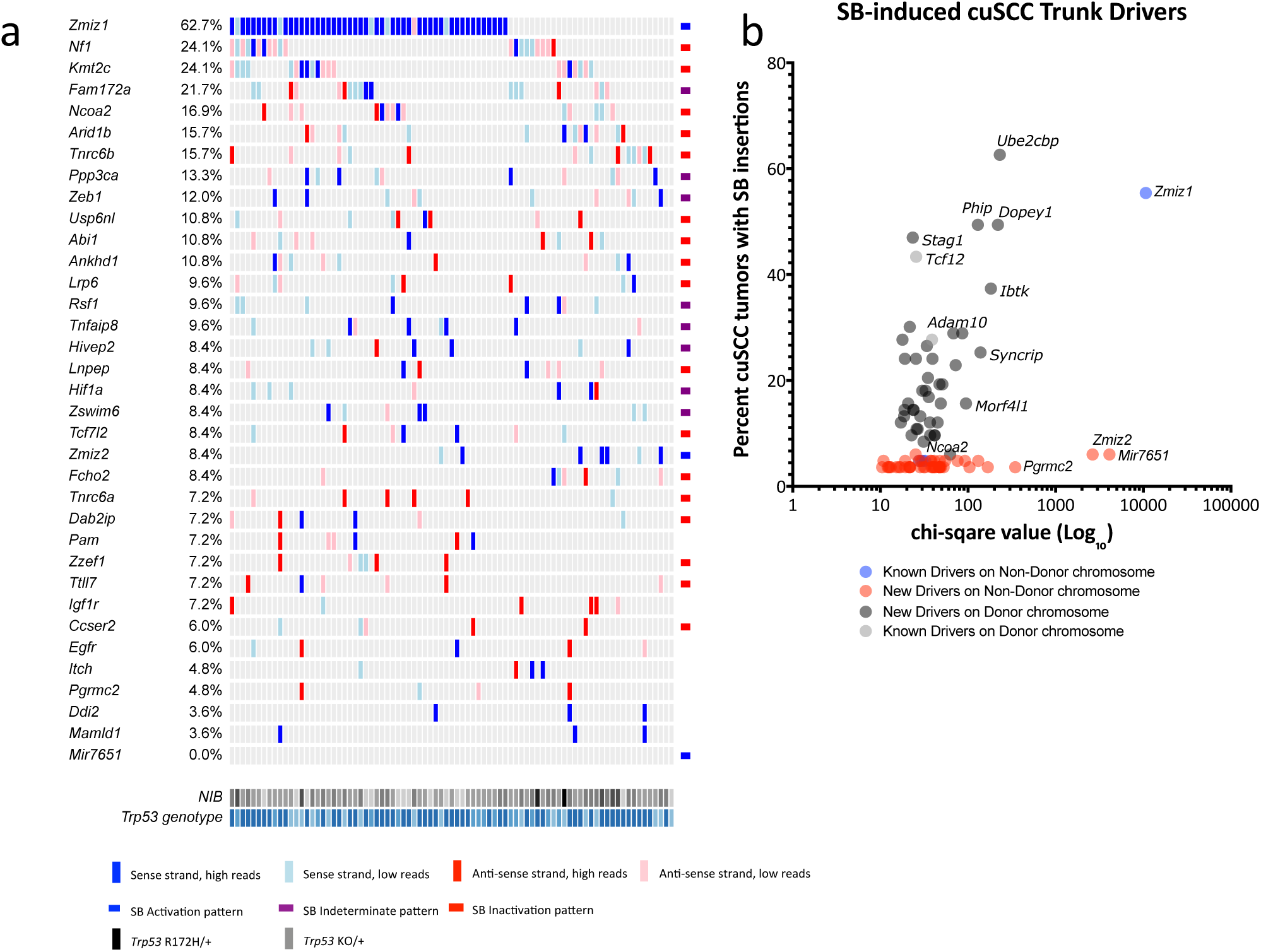
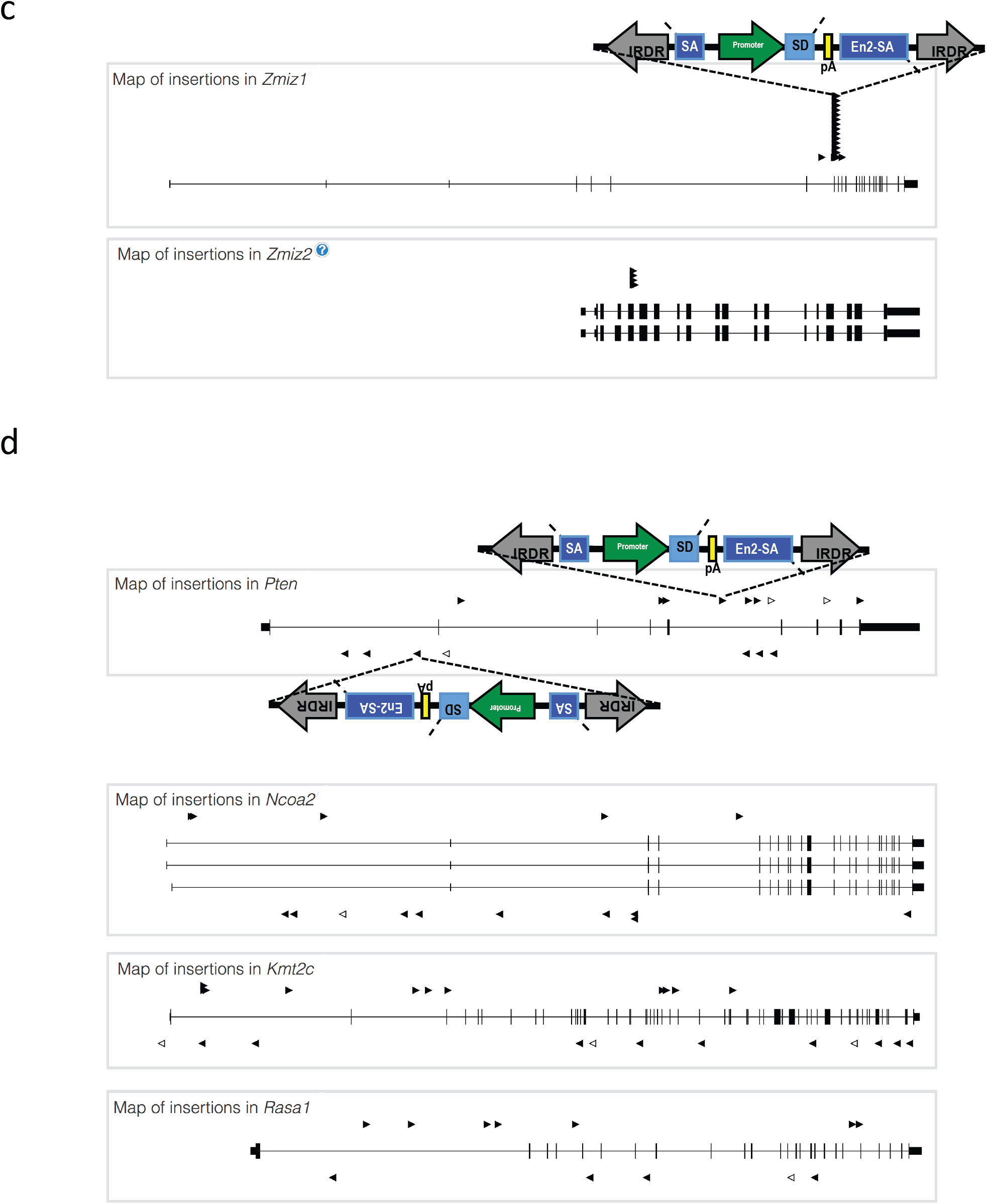
Landscape of candidate trunk drivers mutated in SB-induced cuSCC with Splink_454 sequencing. SB Driver Analysis applied to Splink_454 SB insertion data (**Supplementary Table 4**) with read depth cutoff of 12 on 83 late-stage cuSCC genomes. (**a**) FWER significant driver genes (rows) were sorted by occurrence within the cohort genomes (columns). Waterfall plots were generated on all insertions, and insertion patterns determined by progression driver analysis are shown to the side. Normalized insertion burden (NIB) from lowest (light gray) to highest (black). *Trp53* genotype shown from Trp53 ^+/+^ (light blue), Trp53 ^R172H/+^ (blue) and Trp53 ^KO/+^ (dark blue). (**b**) Trunk cuSCC Driver genes containing SB insertion events with 10 or more 454 sequence reads per tumor. (**c-d**) Representative SB insertion maps in cuSCC trunk drivers showing the locations of mapped SB insertions (triangles) within (**c**) activating (oncogenic) pattern SB insertions into *Zmiz1* and *Zmiz2* or (**d**) Inactivating (tumor suppressive) pattern of SB insertions into *Pten, Ncoa2, Kmt2c*, and *Rasa1* from SB|cuSCC genomes. *(panels* ***c*** *and* ***d*** *continue on next page)*

**Supplementary Figure 4:**
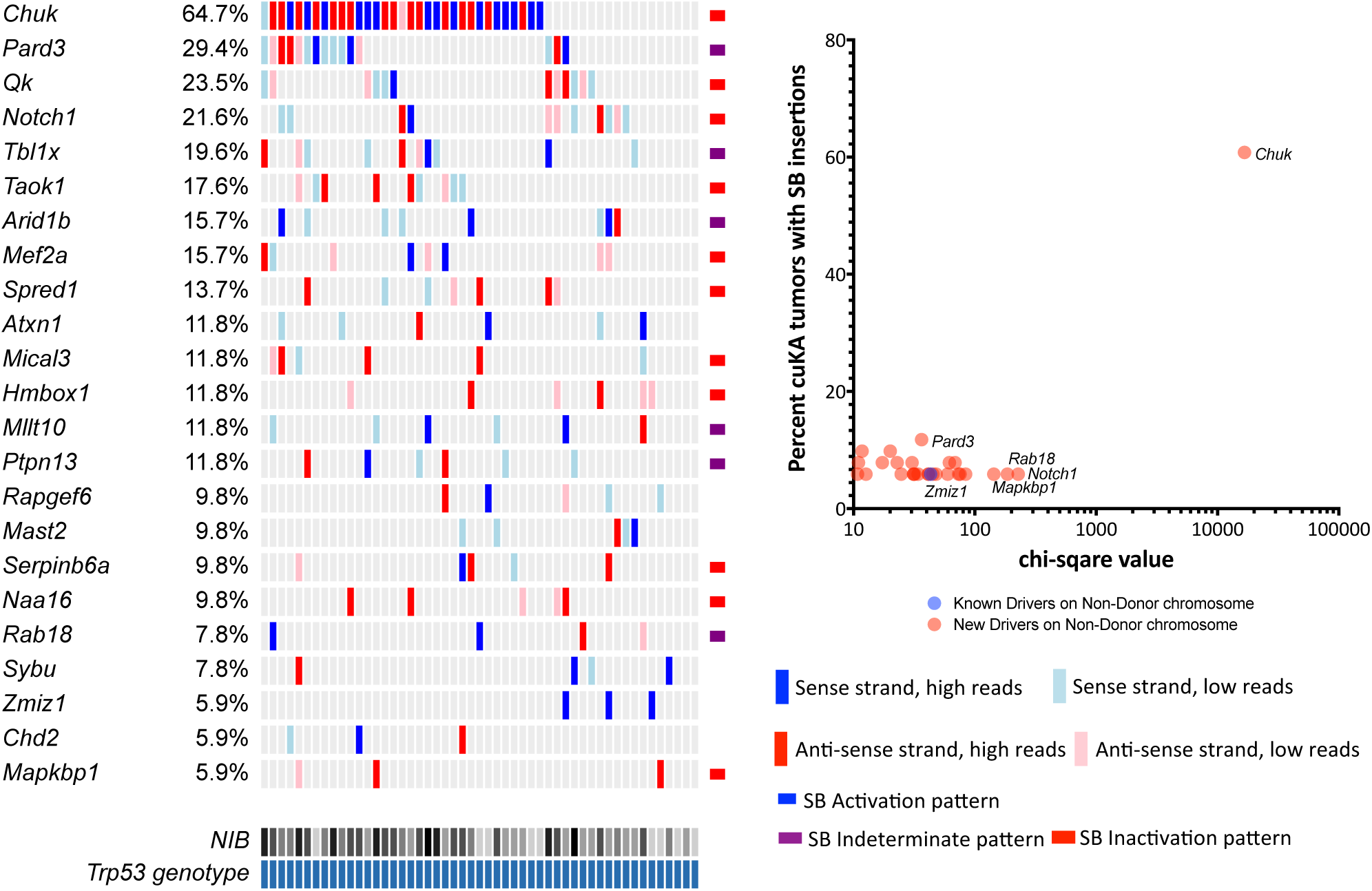
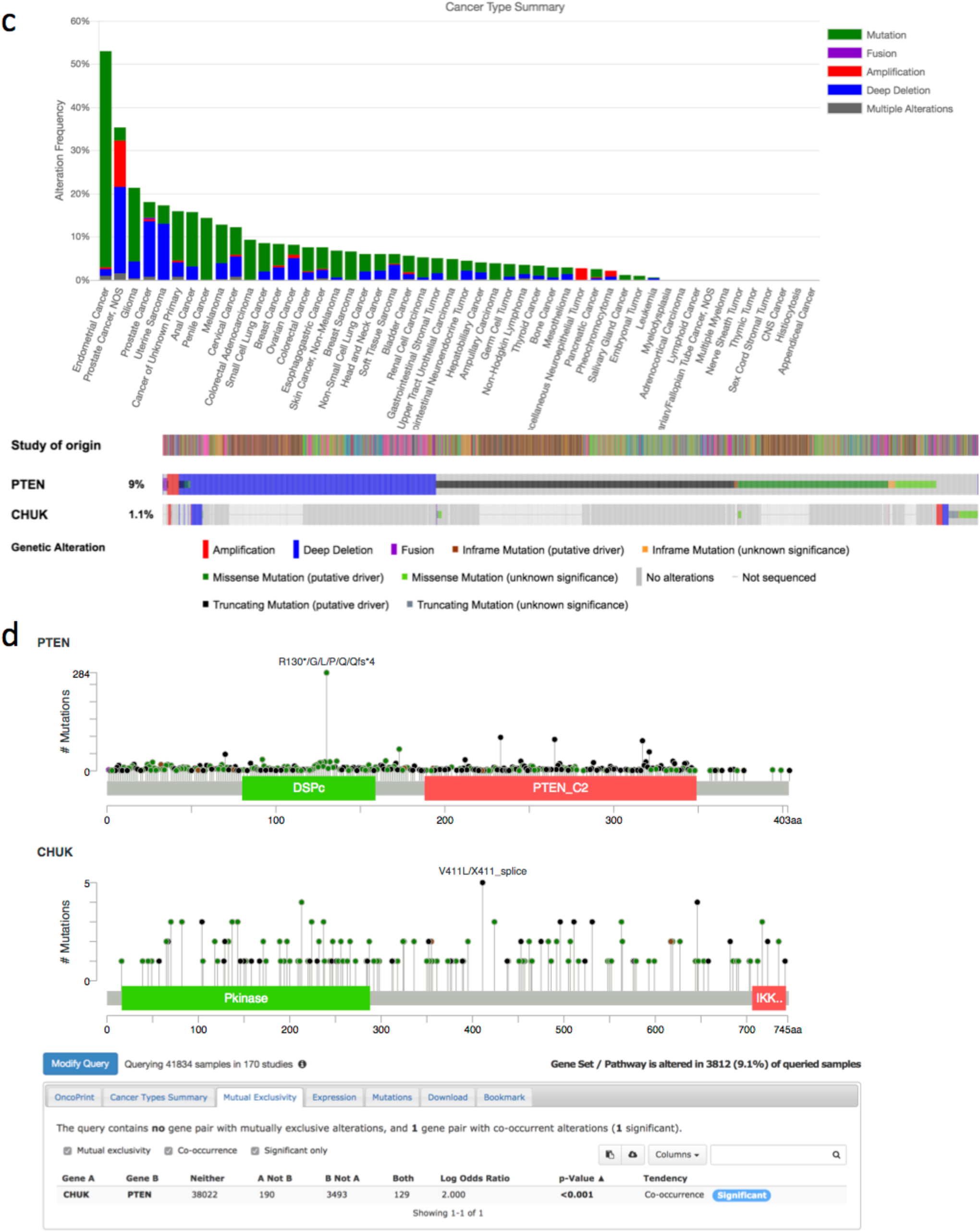
Landscape of candidate trunk drivers mutated in SB-induced cuKA arising from the SB|*Krt5*-Cre|*Pten*^floxed/+^cohort with Splink_454 sequencing. SB Driver Analysis applied to Splink_454 SB insertion data (**Supplementary Table 7**) with read depth cutoff of 20 on 83 late-stage cuSCC genomes. (**a**) FWER significant driver genes (rows) were sorted by occurrence within the cohort genomes (columns). Waterfall plots were generated on all insertions, and insertion patterns determined by progression driver analysis are shown to the side. Normalized insertion burden (NIB) from lowest (light gray) to highest (black). *Trp53* genotype shown from *Trp53* ^+/+^ (light blue), *Trp53* ^R172H/+^ (blue) and *Trp53* ^KO/+^ (dark blue). (**b**) Trunk cuKA Drivers genes containing SB insertions events with 10 or more 454 sequence reads per tumor. (**c**) PTEN and CHUK alteration occurrences and Oncoprint across bBioPortal datasets. (**d**) PTEN and CHUK protein coding alterations and co-occurrence values from cBioPortal. *(panels* ***c*** *and* ***d*** *continue on next page)*

**Supplementary Figure 5:**
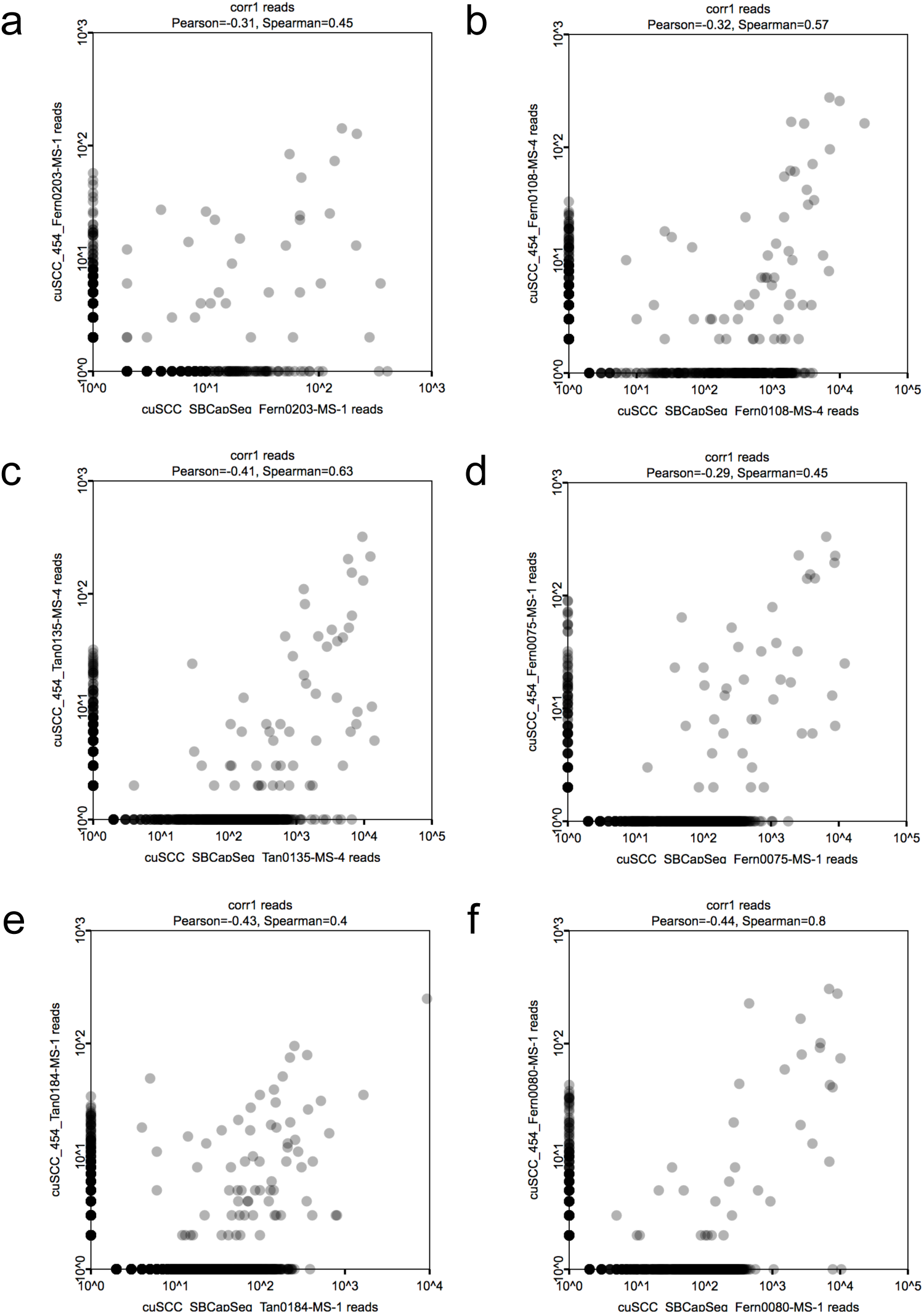
Comparison of SB insertion results using SBCapSeq versus splink_454 methods. Representative plots of the individual SB insertion sites based on read depth from six individual cuSCC genomes (**a-f**) from genomic DNAs created from bulk tumor specimens from biological replicate libraries (independent library workflows applied to the same biological specimen isolate) comparing SBCapSeq (*x*-axis) versus splink_454 (*y*-axis) methods. Compared to Splink_454, the SBCapSeq method identifies more SB insertion events and overall higher read depth sites, however many of the high read depth sites were highly reproducible. Across the individual cuSCC genomes in the cohort, we observe highly variable reproducibility (from low to moderate) between the sequencing methods, supported by both Pearson’s and Spearman’s correlation metrics (**a-f**). We find SBCapSeq to be universally superior to 454 sequencing with little difference between the various SBCapSeq protocol variants tested, including SBCapSeq v1 (**a-b**) or SBCapSeq v2 (**c-d**) with individual captured libraries or SBCapSeq v2 (**e-f**) with multiplex pooled captured libraries.

**Supplementary Figure 6:**
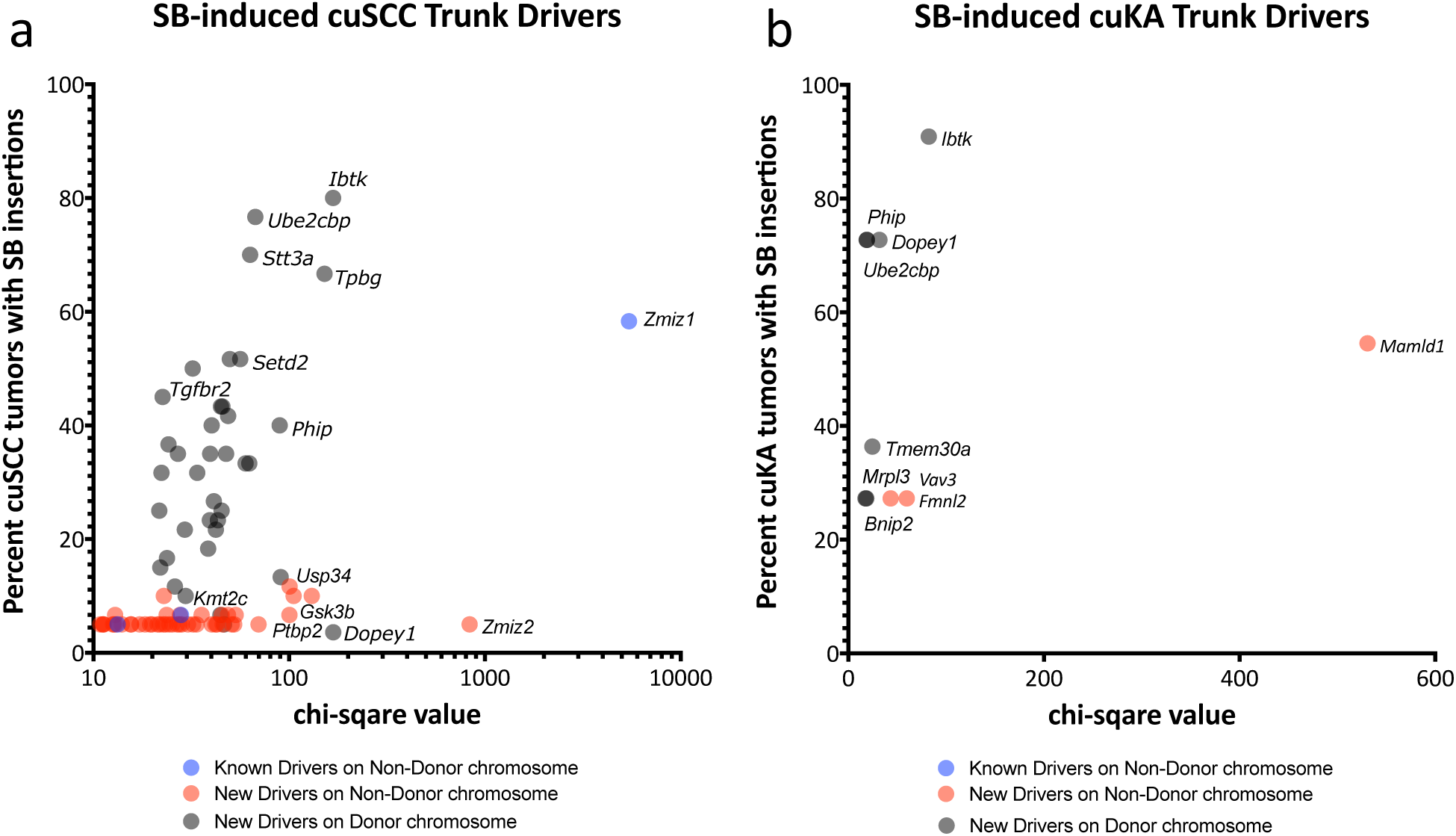
Candidate trunk drivers mutated in SB-induced cuSCC with Splink_454 sequencing. (**a**) Trunk cuSCC Drivers genes from SB Driver Analysis applied to SBCapSeq insertion data (**Supplementary Table 4**) with read depth cutoff of 300 on 84 late-stage cuSCC genomes. Trunk cuSCC Driver genes. (**b**) Trunk cuKA Drivers genes from SB Driver Analysis applied to SBCapSeq insertion data (**Supplementary Table 7**) with read depth cutoff of 300 on 11 early-stage cuKA genomes.

**Supplementary Figure 7:**
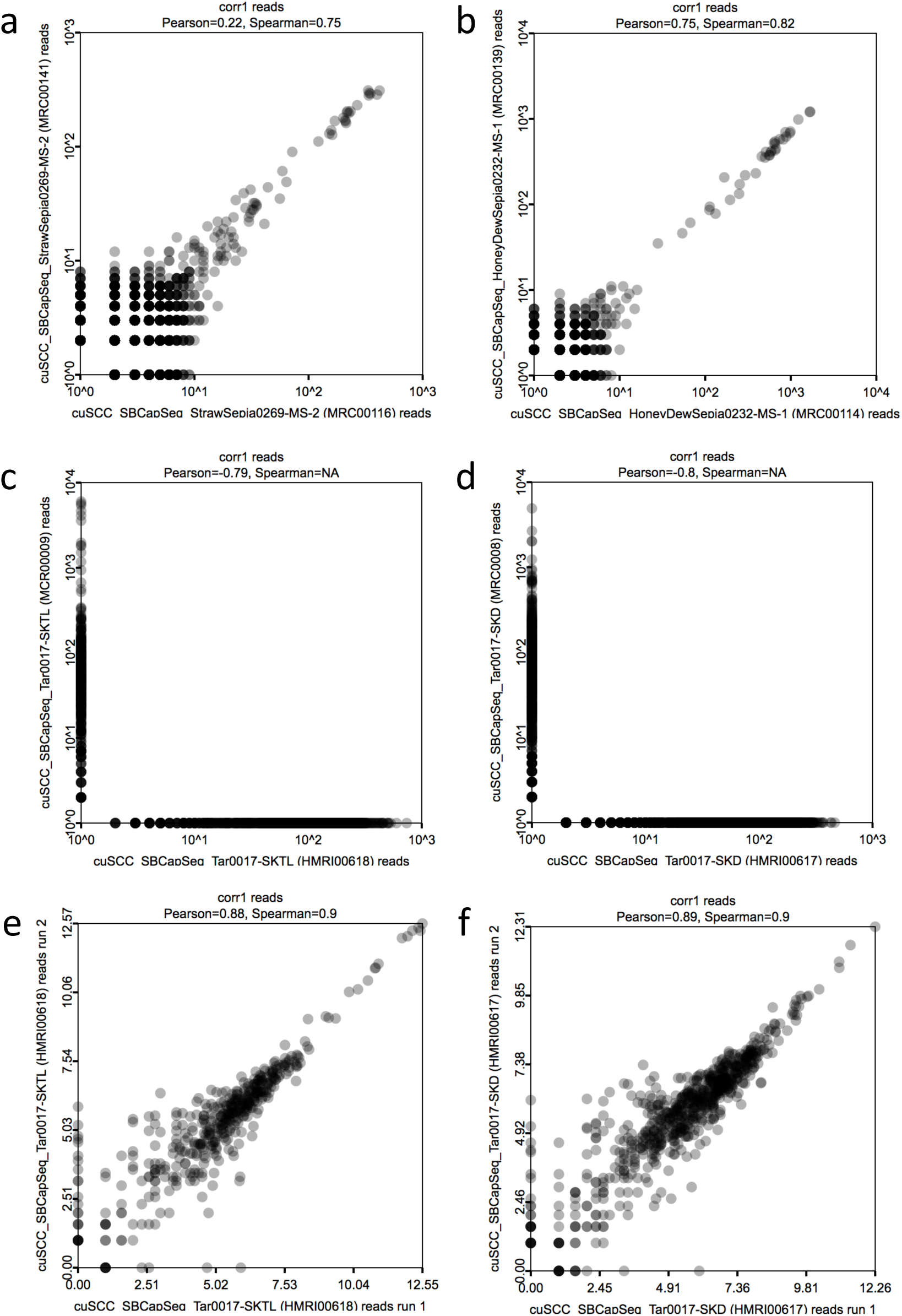
Evaluating the reproducibility of SBCapSeq results from bulk cuSCC and normal skin specimens. Representative plots of the individual SB insertion sites based on read depth from two individual cuSCC genomes (**a-b**) and two unselected skin cell genomes from histologically normal skin (**c-d**) from genomic DNAs created from bulk tumor specimens from biological replicate libraries (independent library workflows applied to the same biological specimen isolate) comparing library 1 (*x*-axis) to library 2 (*y*-axis). Biological reproducibility of cuSCC specimen libraries is high, supported by both Pearson’s and Spearman’s correlation metrics, using all of SB insertion sites with read depths of 2 or higher (**a-b**). In contrast, biological reproducibility of normal skin specimen libraries is exceptionally low, supported by both Pearson’s and Spearman’s correlation metrics, and fails entirely to identify the same SB insertion sites (**c-d**), indicating that these are likely background passenger insertion events that are not reproducibly sequenced. However, plotting of the individual SB insertion sites based on read depth from two technical replicates (the same library preparation) from two separate sequencing runs confirms very high reproducibility between individual libraries at high-to-moderate read depths, supported by both Pearson’s (r= 0.48) and Spearman’s (ρ=0.49 or undetermined) correlation metrics (**e-f**), further supporting the lack of biological replication is most likely due to non-clonal background passenger insertion events and do not result from flaws in the SBCapSeq protocol *per se*.

**Supplementary Figure 8:**
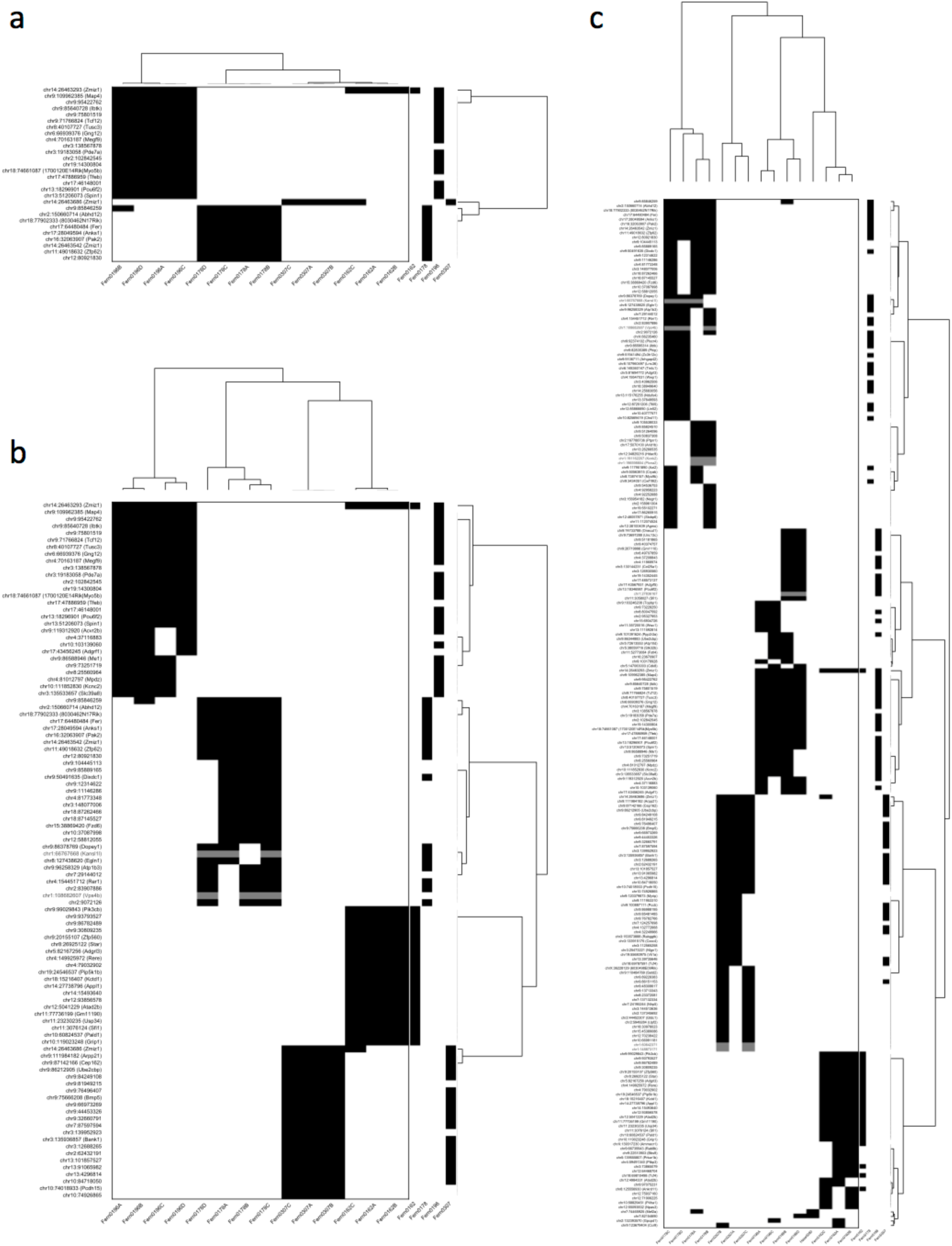
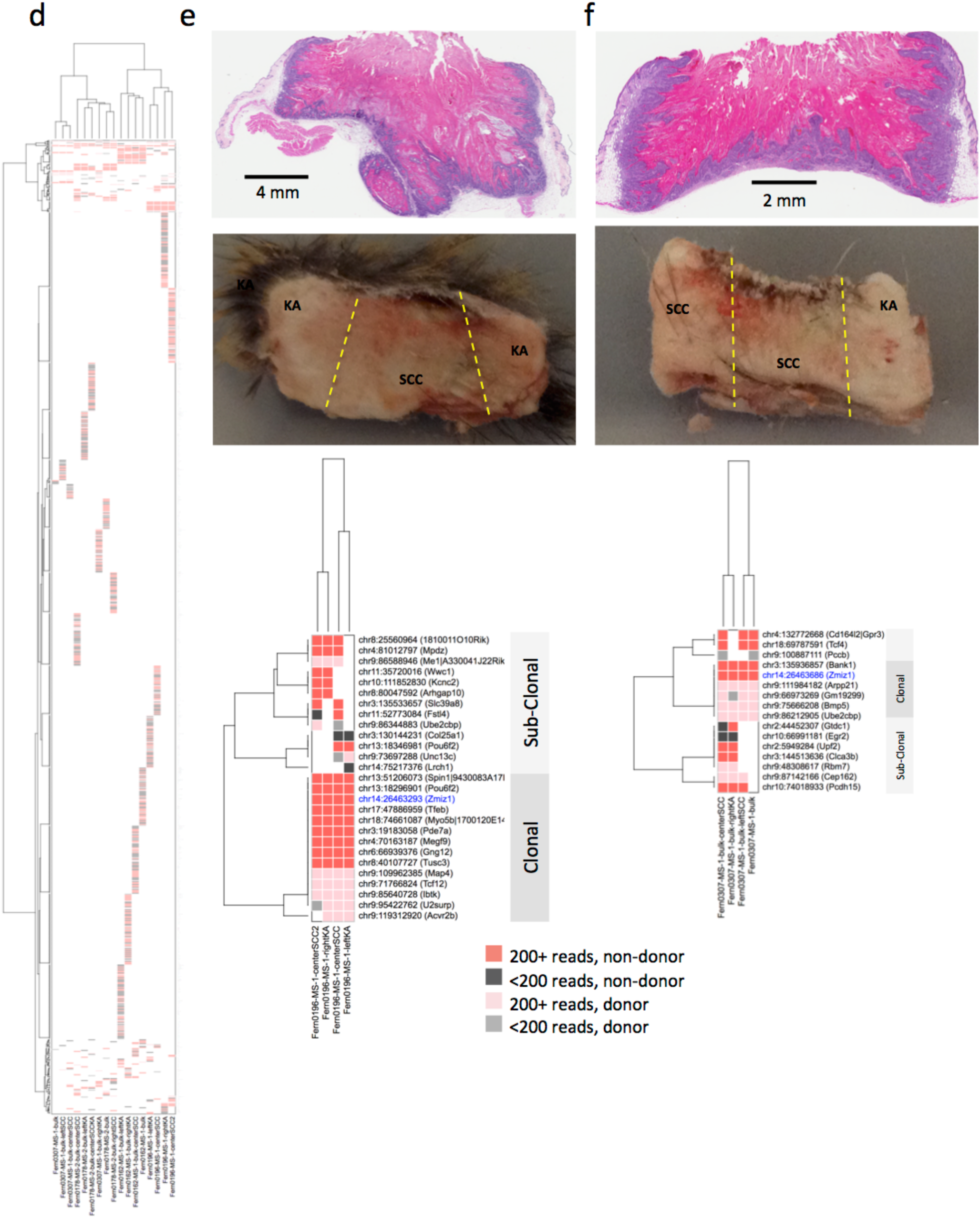
Multi-region SBCapSeq analysis of bi-lesional keratinocyte-derived skin masses containing distinct cuKA and cuSCC histologies. Hierarchical two-dimensional clustering (Hamming distance with the Ward method) of recurrent genic SBCapSeq insertion data from four skin masses containing distinct cuKA and cuSCC regions with ≥4 (**a**), ≥3 or more (**b**), or ≥2 (**c**), recurrent SB insertions. At each recurrent threshold, four distinct specimen (*x*-axis) and gene (*y*-axis) clades, each pertaining to a single bi-lesional mass, were detected, thus demonstrating clonal outgrowth. Genomic DNA was isolated from adjacent, non-overlapping serial sections and three or four regions dissected from histologically distinct regions; the tumor reference, consisting of a cross-section of each mass (including all regions), is plotted as independent tracks on the right of each graph (see **panels e-f** and **Fig. 3 a-b**). (**d**) Hierarchical two-dimensional clustering (Hamming distance with the Ward method) of recurrent genic SBCapSeq insertion data from four skin masses containing distinct cuKA and cuSCC regions. Four distinct specimen (*x*-axis) and gene (*y*-axis) clades, each pertaining to a single bi-lesional mass, were detected, demonstrating clonal identities. *Zmiz1* was the only gene recurrently mutated across all samples with high read depths; most SB insertions were private to a single sample and consistently had lower read depths suggesting these are unselected passenger insertions. Genomic DNA was isolated from adjacent serial sections and included a “bulk” reference consisting of a cross-section of the entire mass, and three or four regions dissected from histologically distinct regions (see **panels e-f**). (**e-f**) Two masses with Aperio slide scans of H&E stained FFPE specimens showing distinct cuKA and cuSCC histologies suggestive of clonal evolution of cuSCC differentiation within cuKA masses and the adjacent flash frozen tissue prior to sectioning for genomic DNA isolation. Hierarchical two-dimensional clustering (Hamming distance with the Ward method) of recurrent genic SBCapSeq insertion data shown below each specimen demonstrates trunk (common to all samples) and sub-clonal, lesion-specific SB insertions.

**Supplementary Figure 9:**
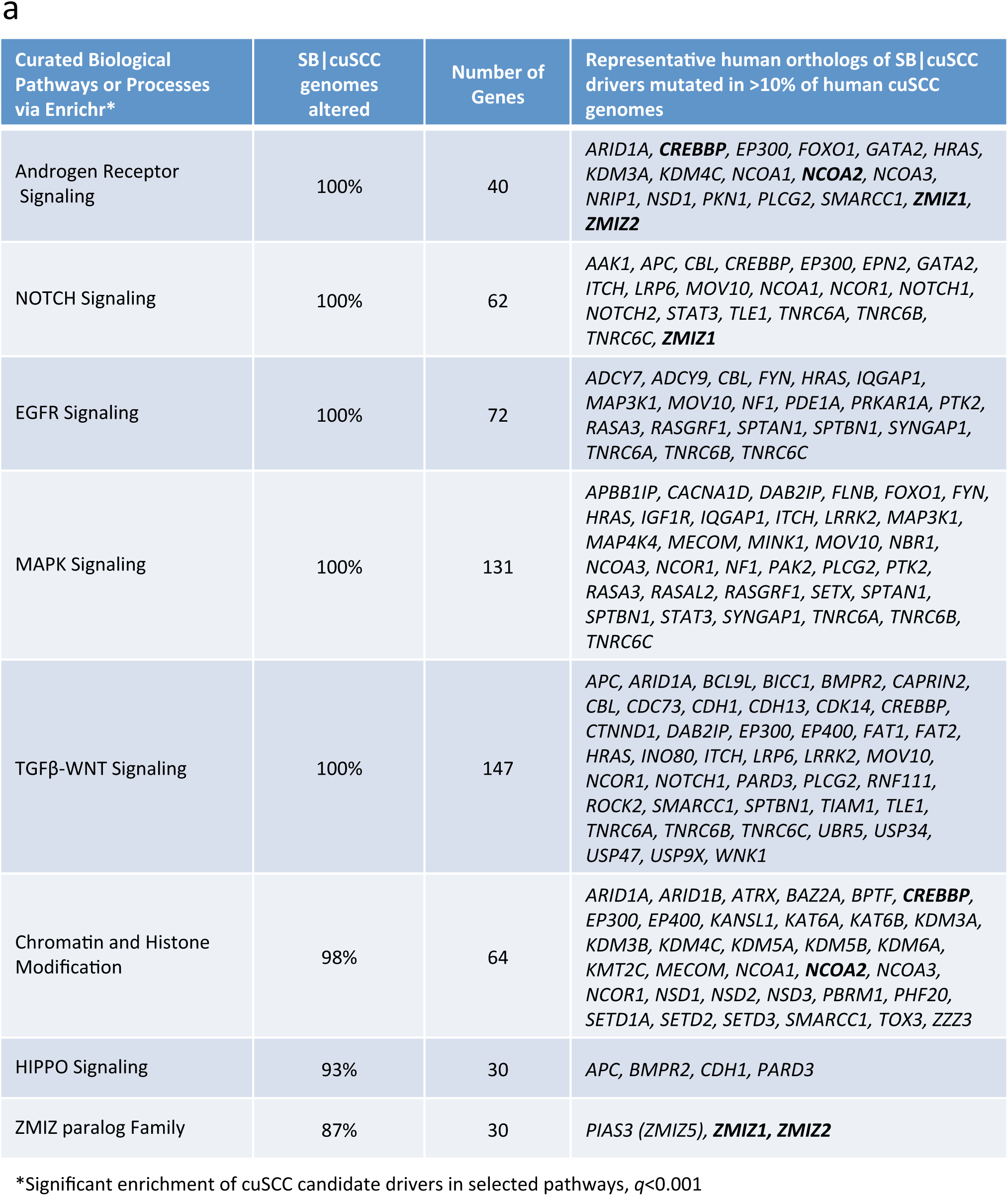
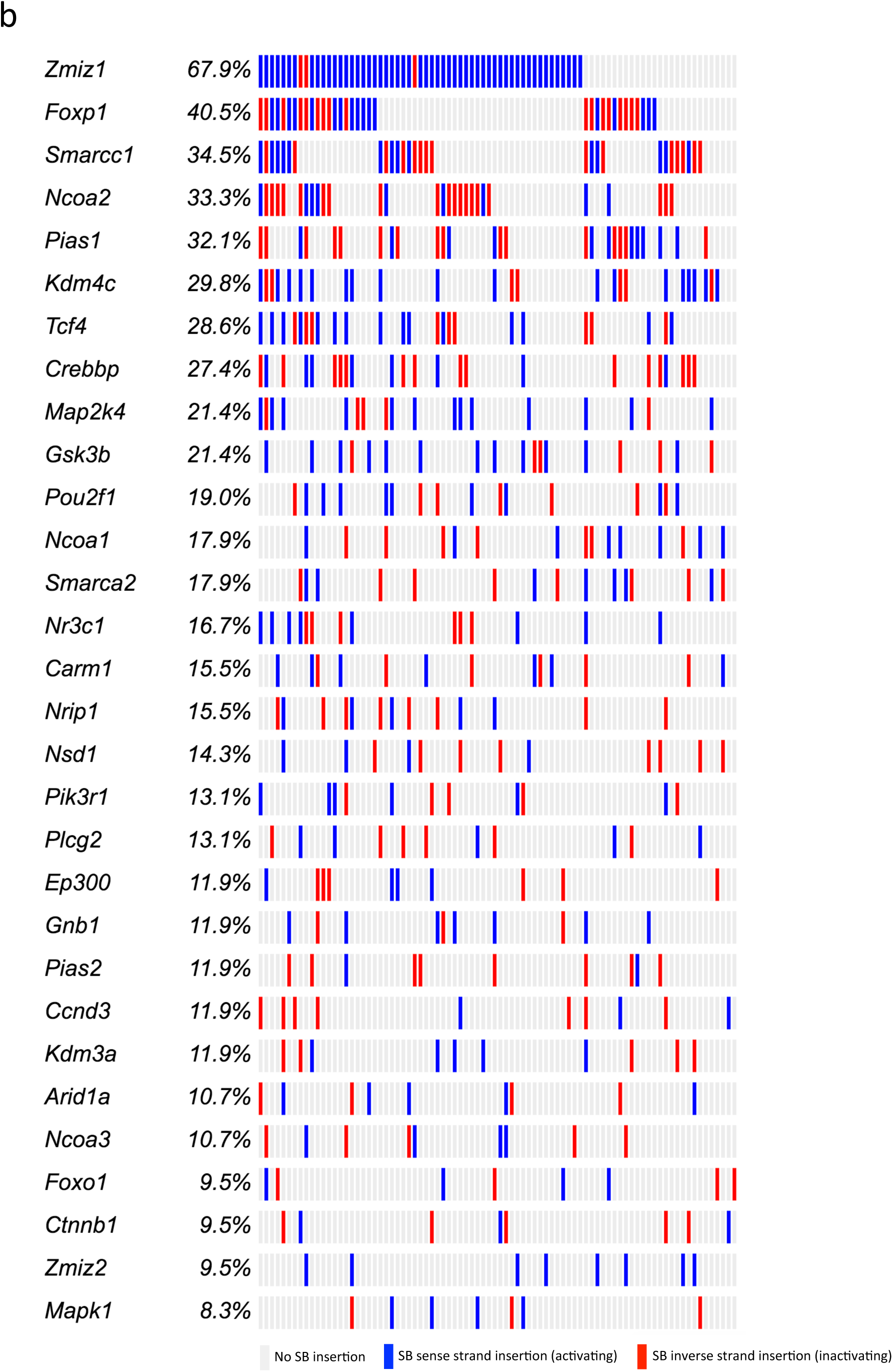
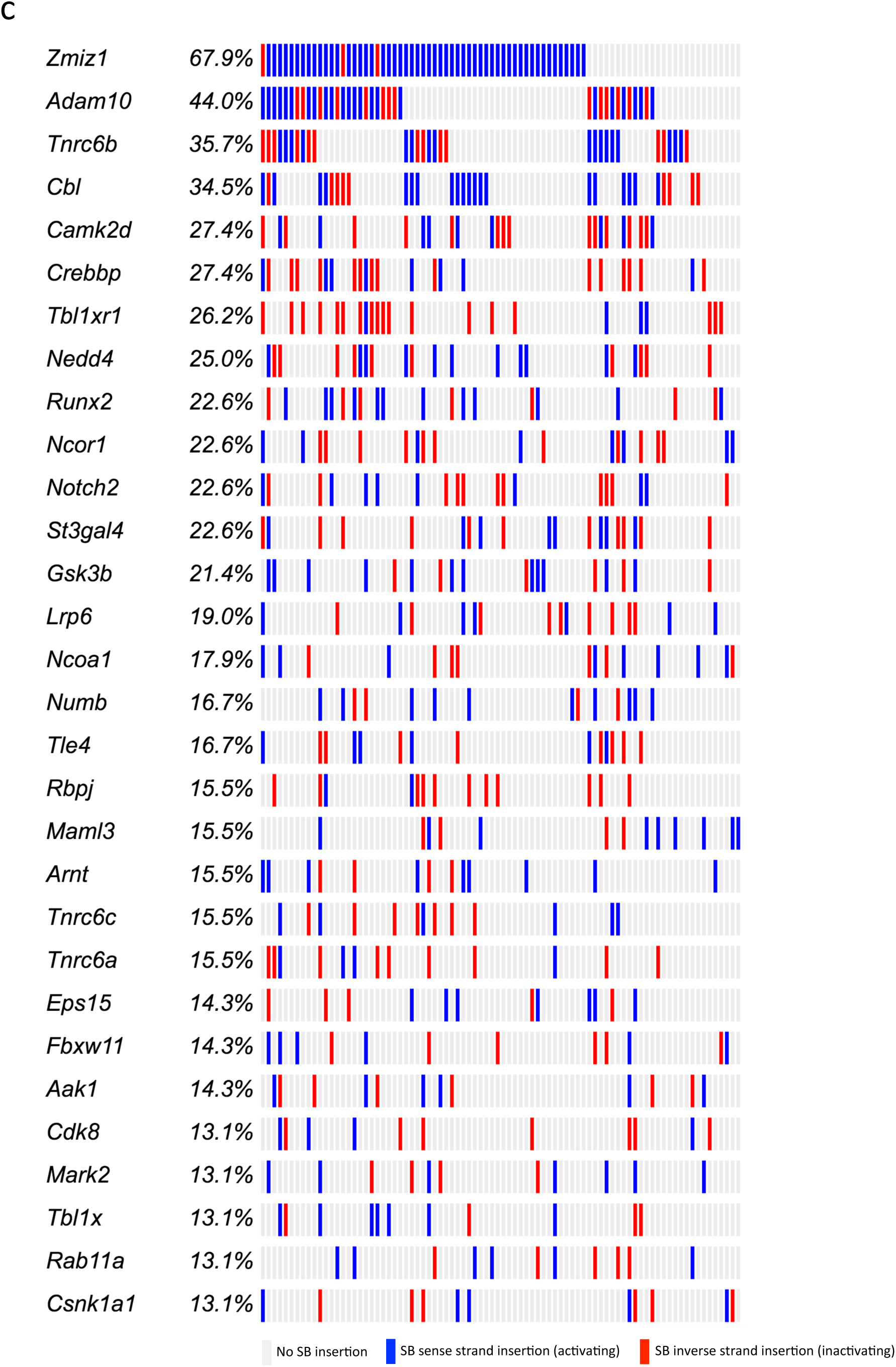
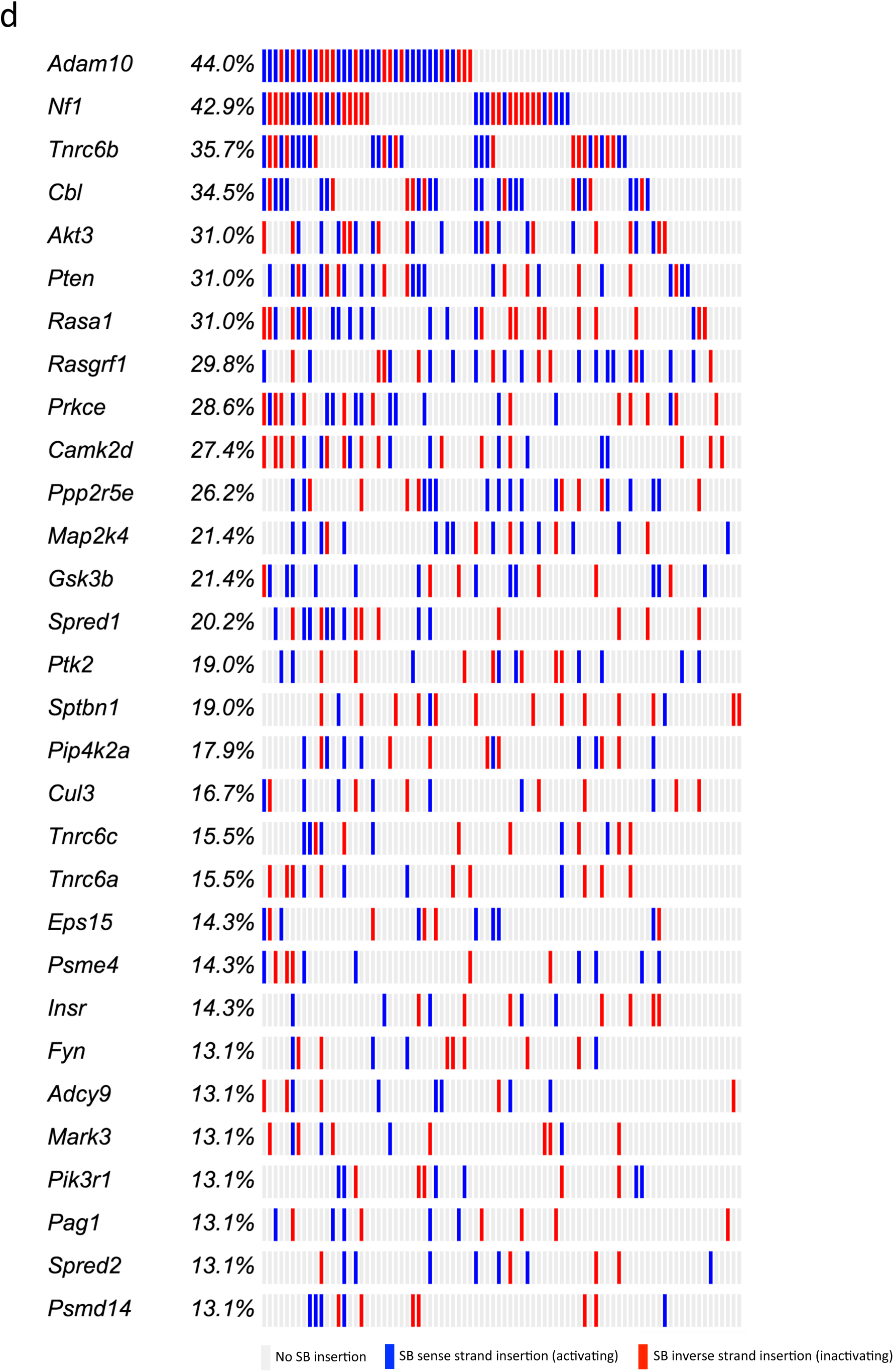
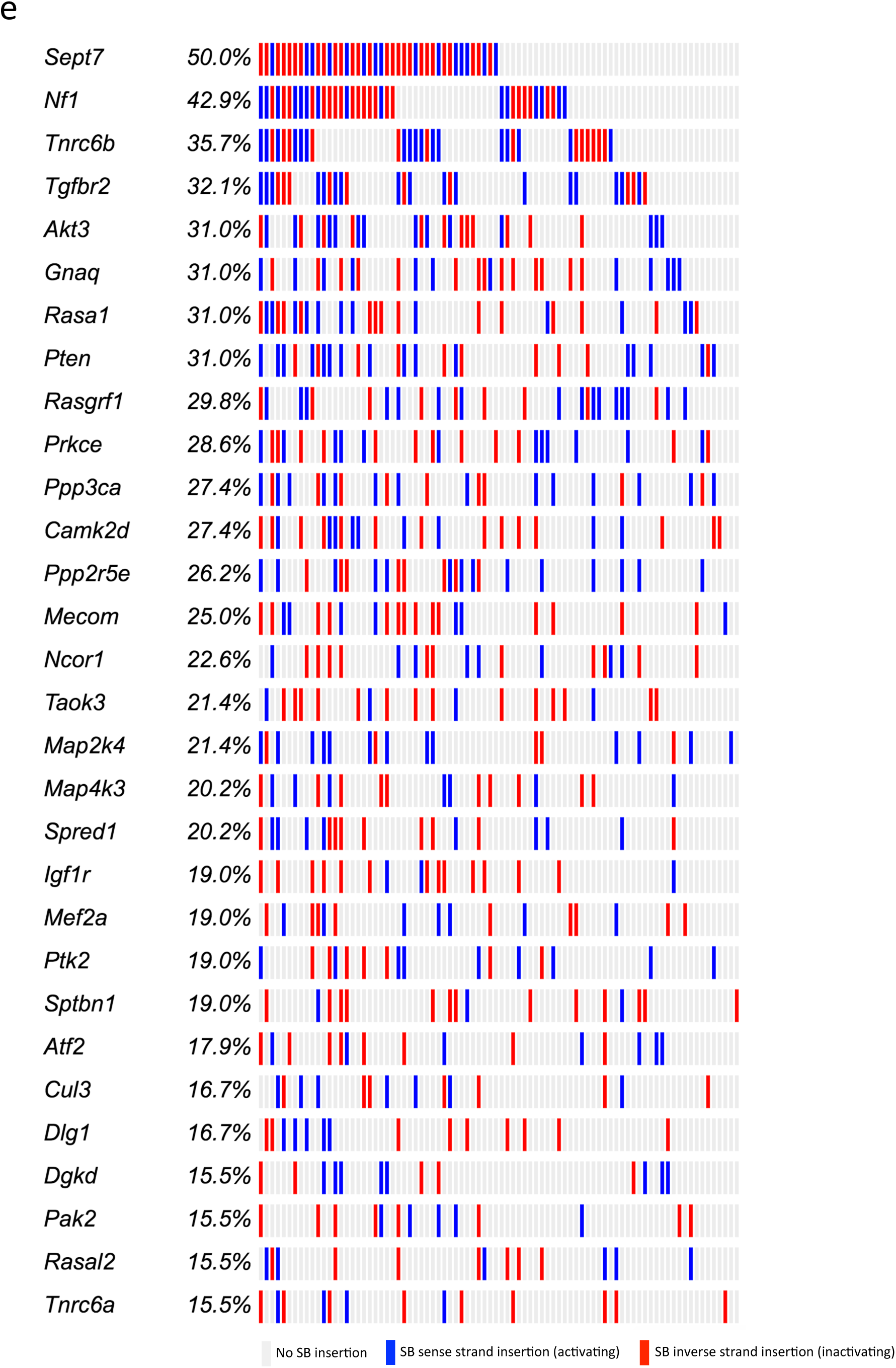
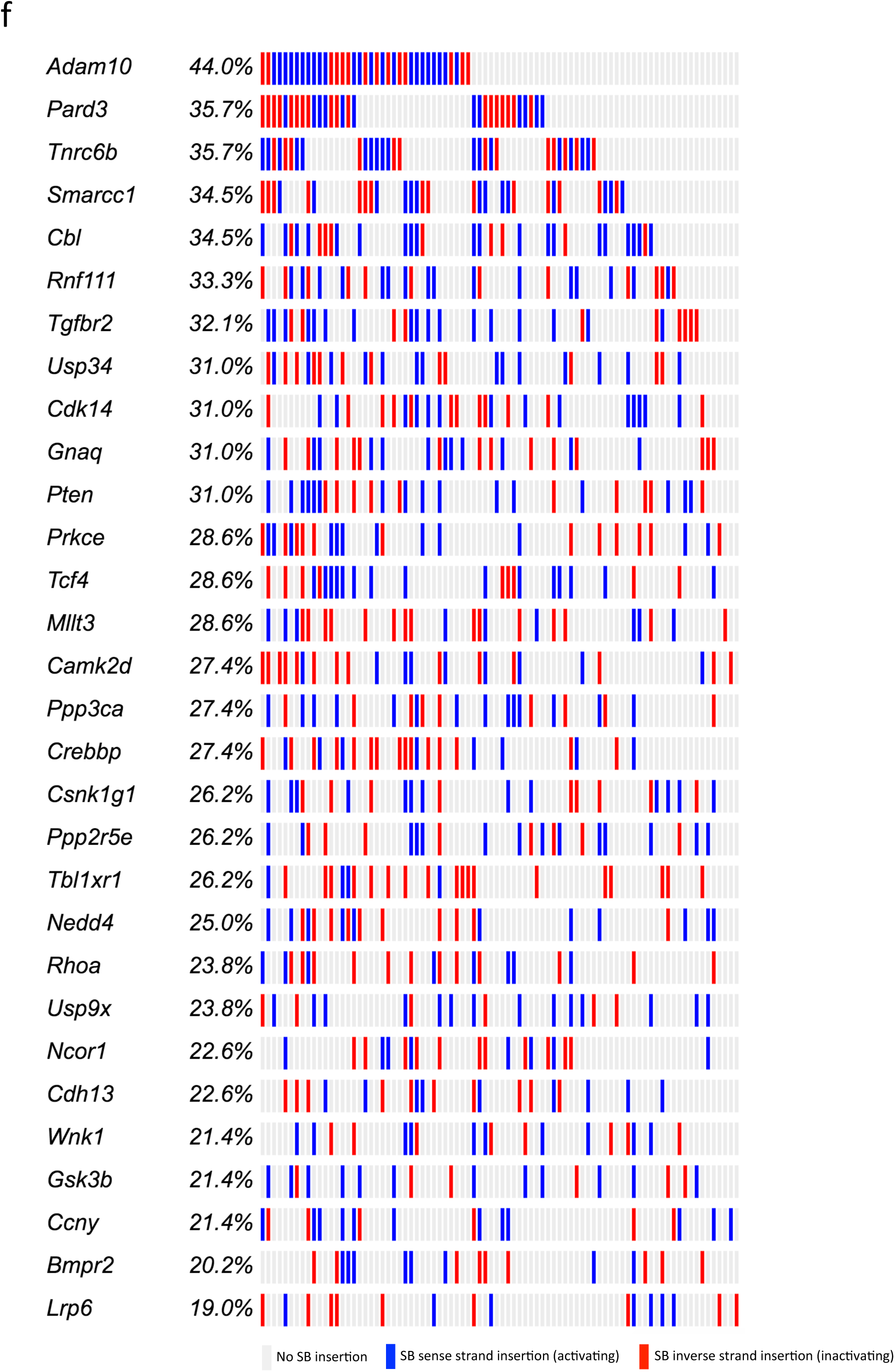
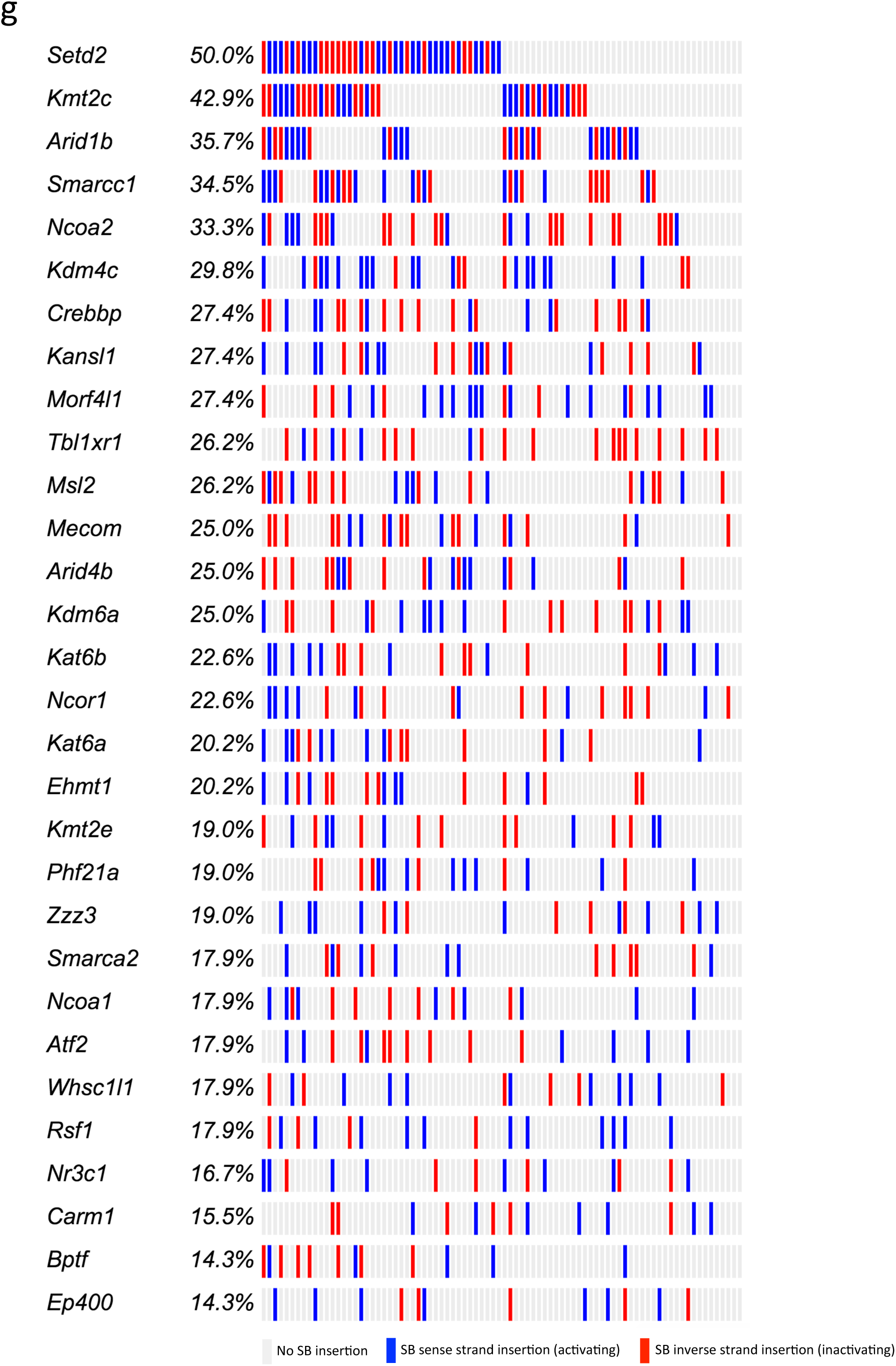
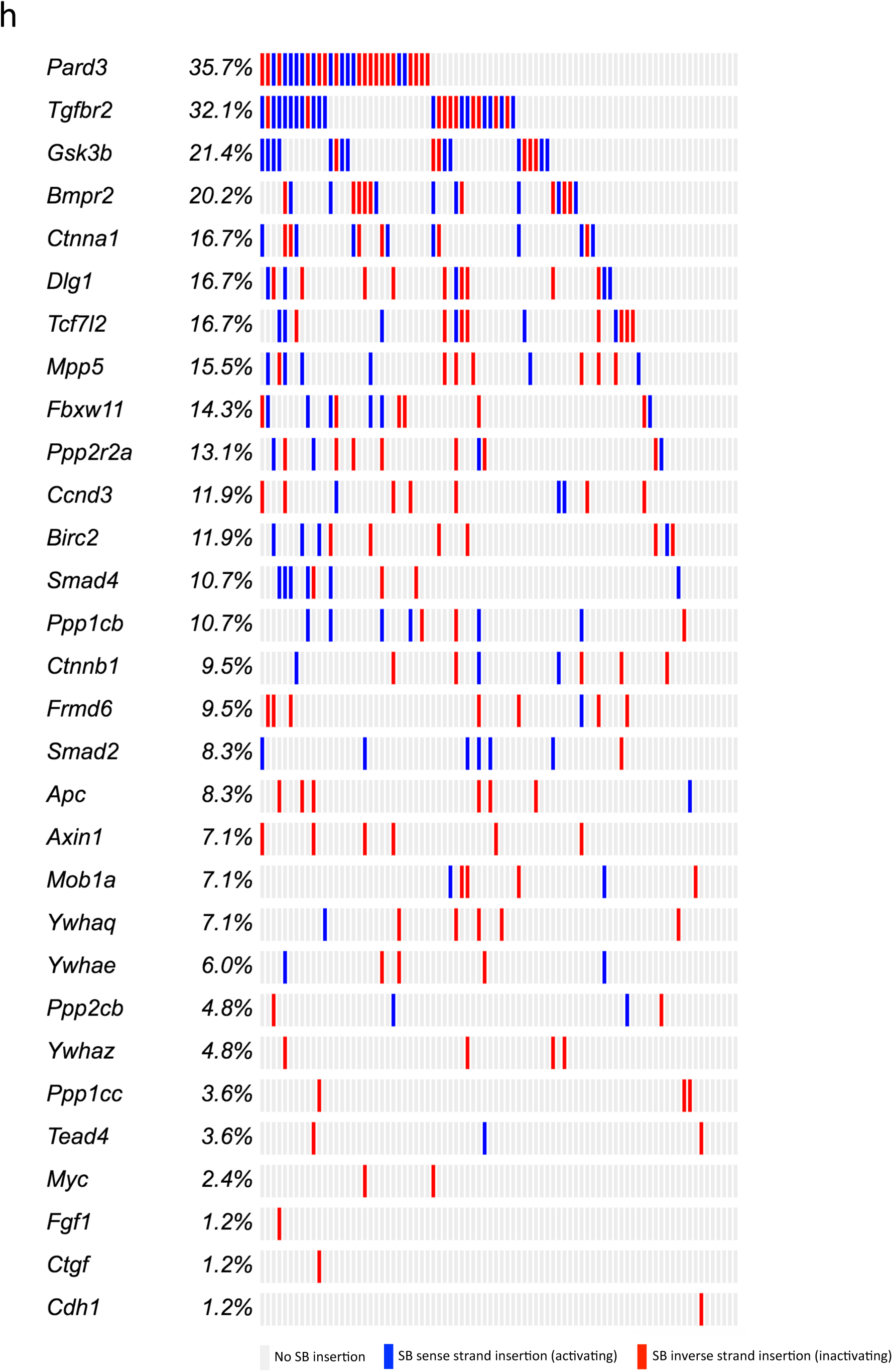
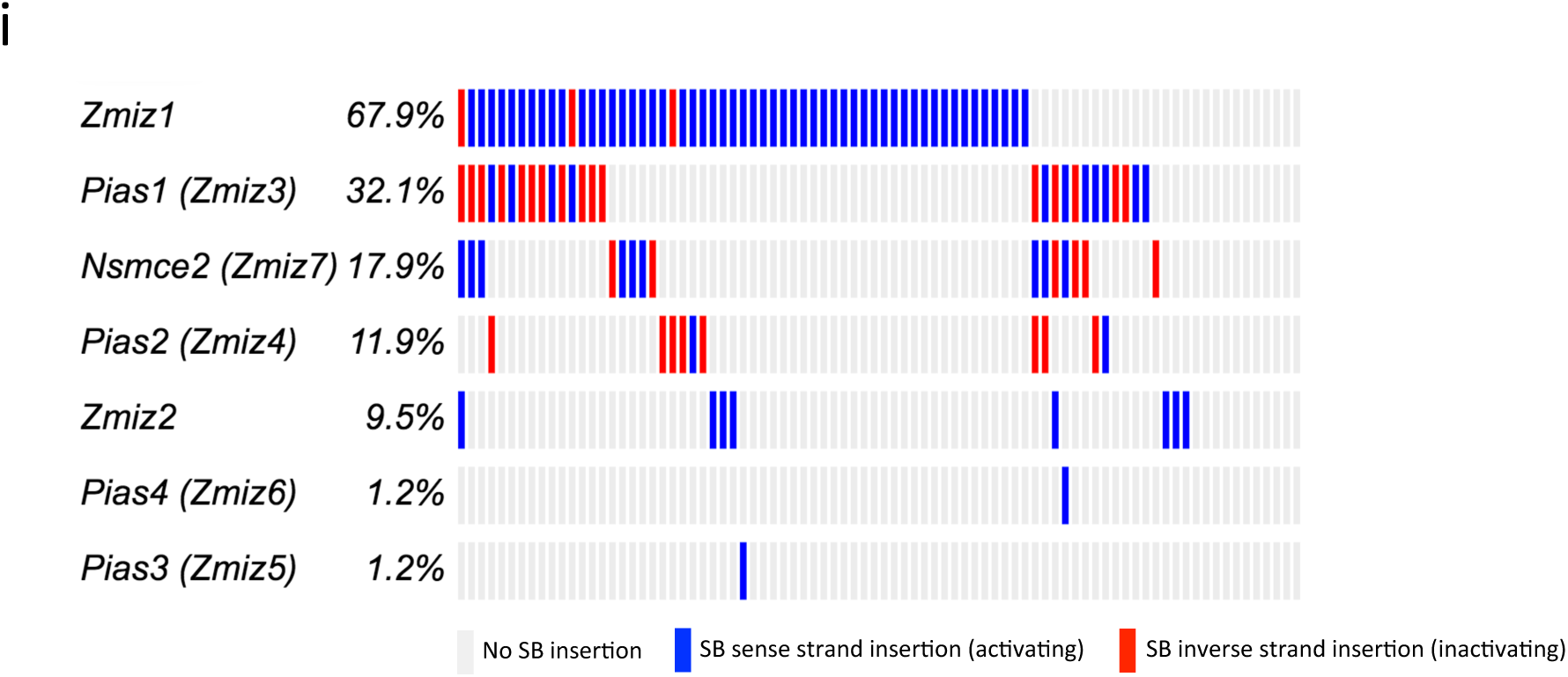
Curated biological pathways and processes enriched within SB-induced cuSCC. (**a**) Table of significant pathways collated from pathway enrichment categories in **Supplementary Table 28**. Oncoprints of the top 30 mutated cuSCC candidate drivers enriched in (**b**) Androgen Receptor Signaling, (**c**) NOTCH Signaling, (**d**) EGFR Signaling, (**e**) MAPK Signaling, (**f**) TGFβ-WNT Signaling, (**g**) Chromatin and Histone Modification, (**h**) HIPPO Signaling, and (**i**) ZMIZ paralog family. *(Panels b–i on next 8 pages)*

**Supplementary Figure 10:**
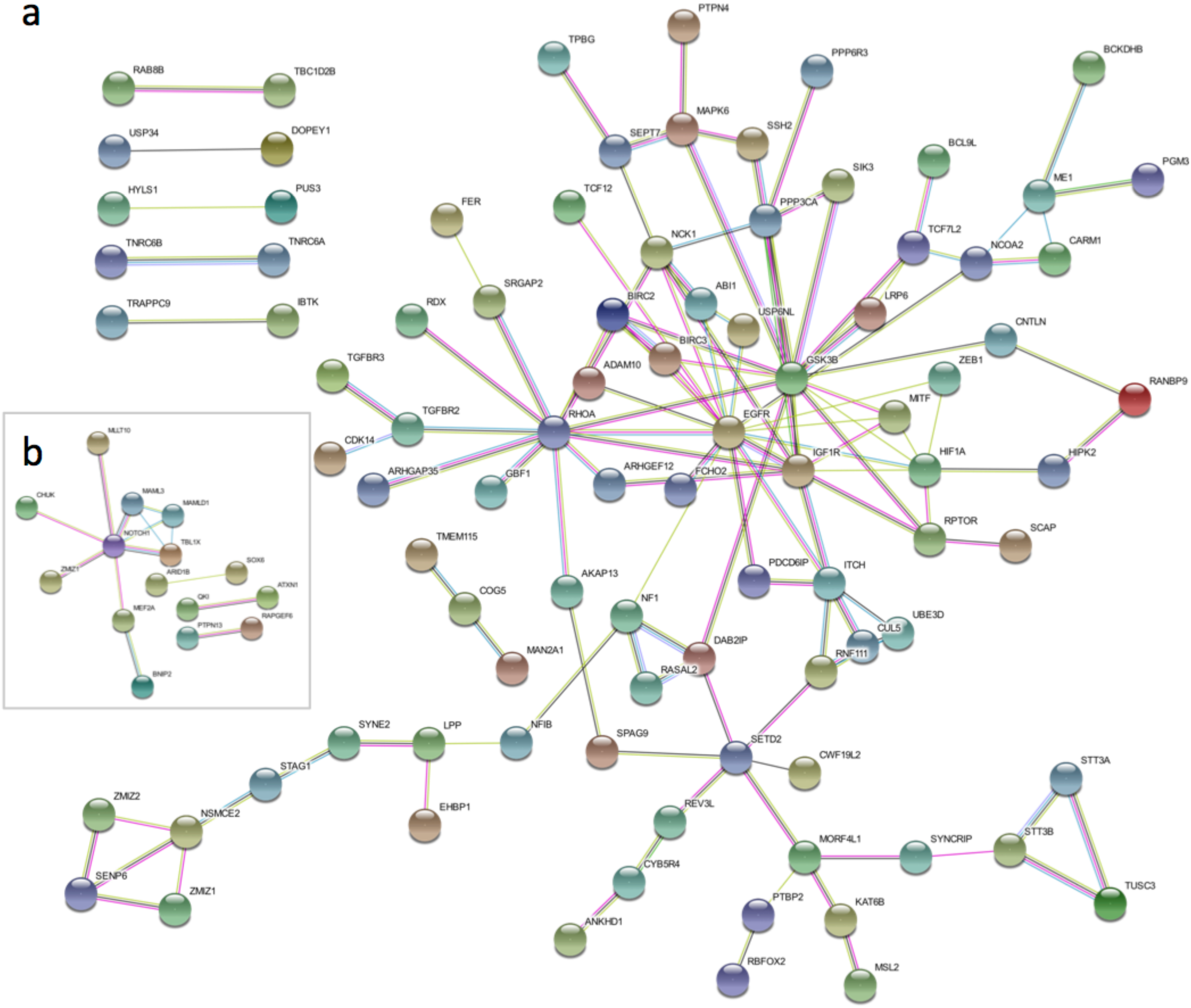
Trunk driver STRING network analysis. (**a**) The SB|cuSCC Trunk Driver (n=84) network has significantly more known protein-protein interactions than expected by chance (*P*=8.55 × 10^−6^8.55e-06, STRING enrichment analysis; number of nodes: 144, number of edges: 124, expected number of edges: 82 (**b**) The SB|cuKA Trunk Driver (n=62) network also has significantly more known protein-protein interactions than expected by chance (*P*=6.93 × 10^−5^, STRING enrichment analysis; number of nodes: 37, number of edges: 14, expected number of edges: 4.

**Supplementary Figure 11:**
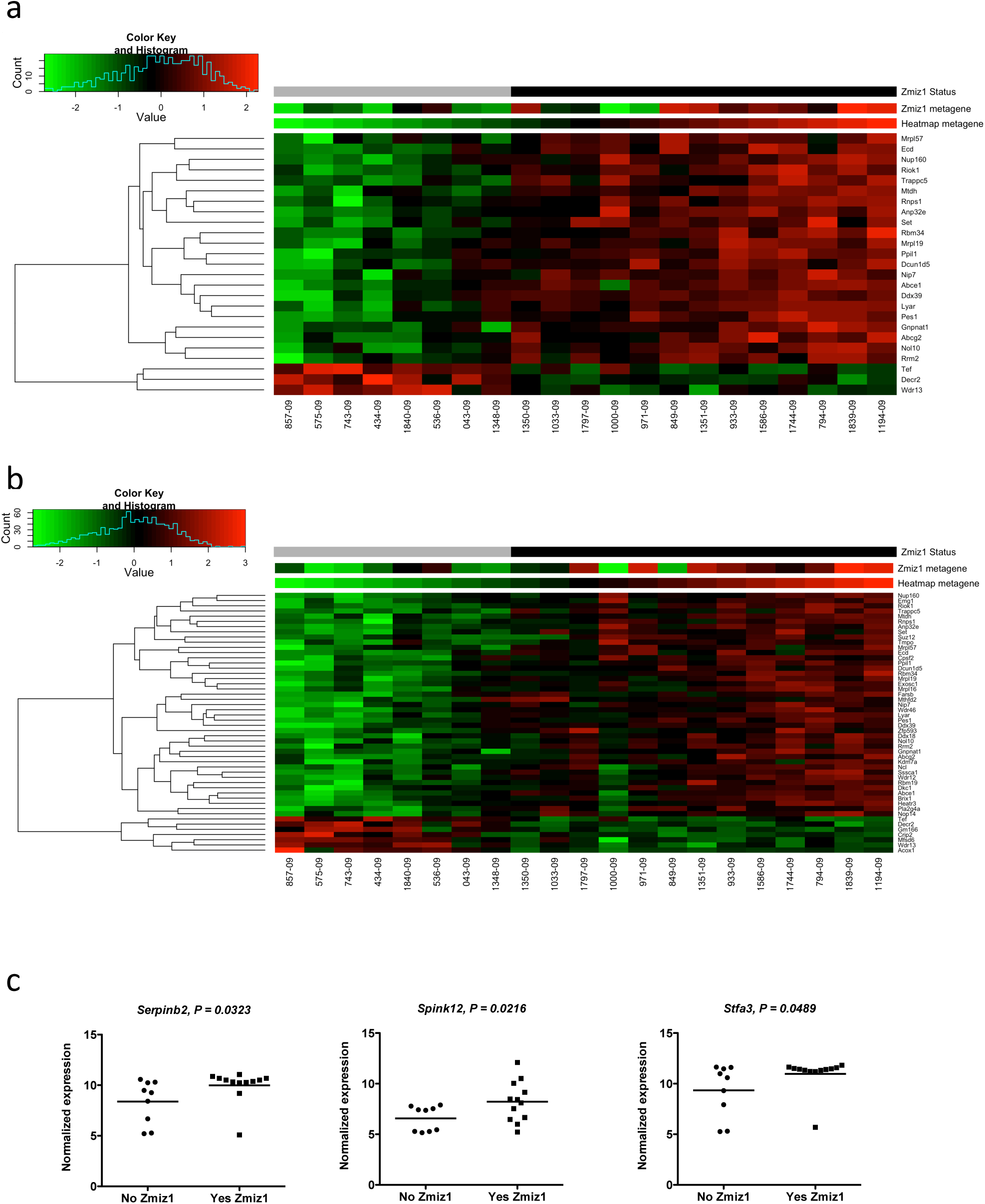
SB effect on gene expression by microarray analysis. (**a**) Top 25 DE genes: *Mtdh, Anp32e, Ecd* are cuSCC progression drivers. (**b**) Top 50 DE genes: *Mtdh, Anp32e, Ecd, Dcun1d5*, and *Kdm7a* are cuSCC progression drivers. (**c**) Differentially expressed genes reported between KA and normal skin after Zmiz1 ectopic expression (Rogers et al.), confirmed with microarray analysis from SB|cuSCC tumors.

**Supplementary Figure 12:**
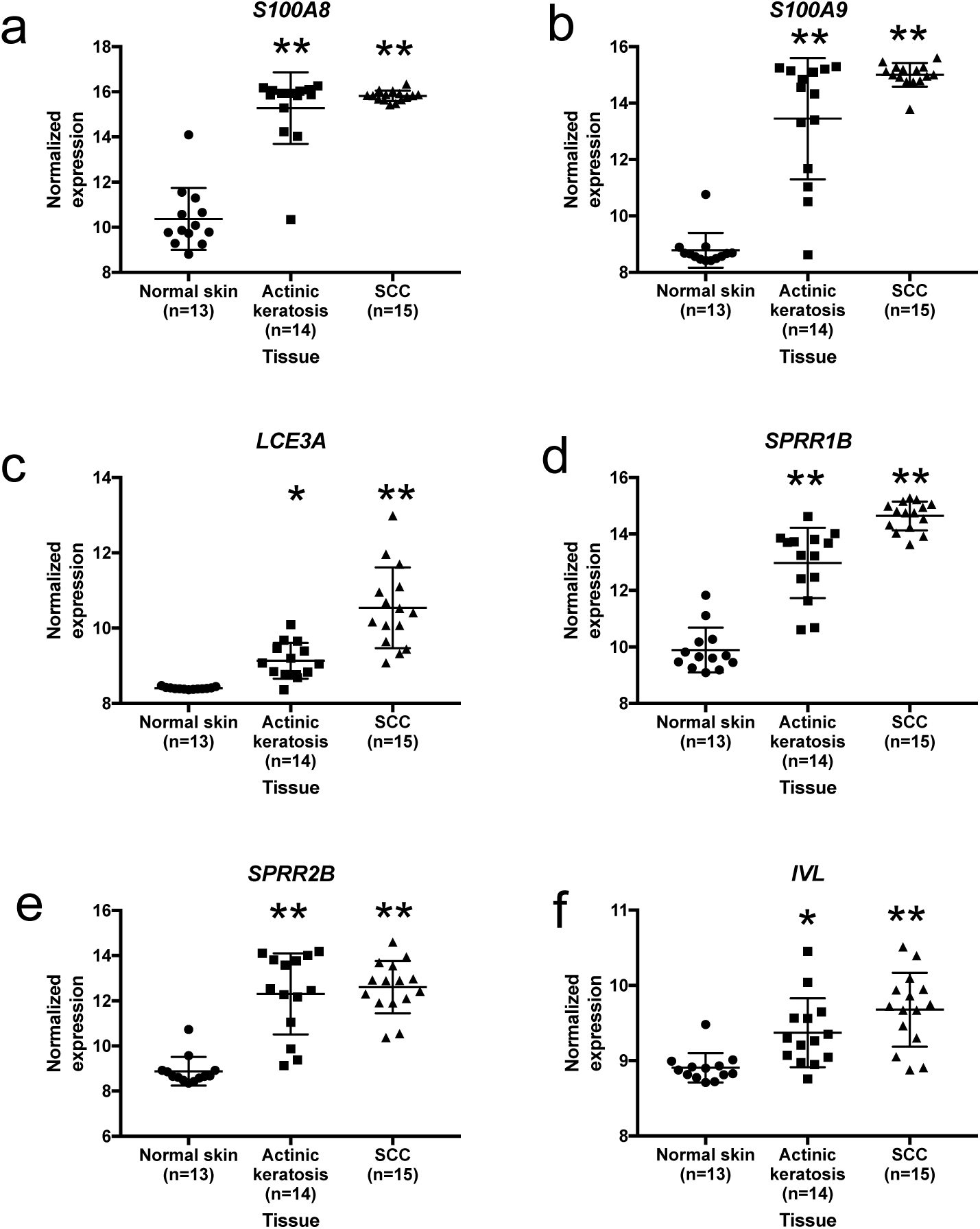
Zmiz1^ΔN185^ driven epidermal differentiation complex genes orthologs are significantly unregulated during human keratinocyte transformation and cuSCC progression. **(a-f)** Expression of selected EDC genes are significantly unregulated in human actinic keratosis (n=14) and cuSCC (n=15) tissues, relative to normal skin (n=13). * adjusted *P* <0.05 vs normal skin; **adjusted *P <* 0.0001 versus normal skin. GEO dataset: GSE32628 (Hameetman et al., 2013).

**Supplementary Figure 13:**
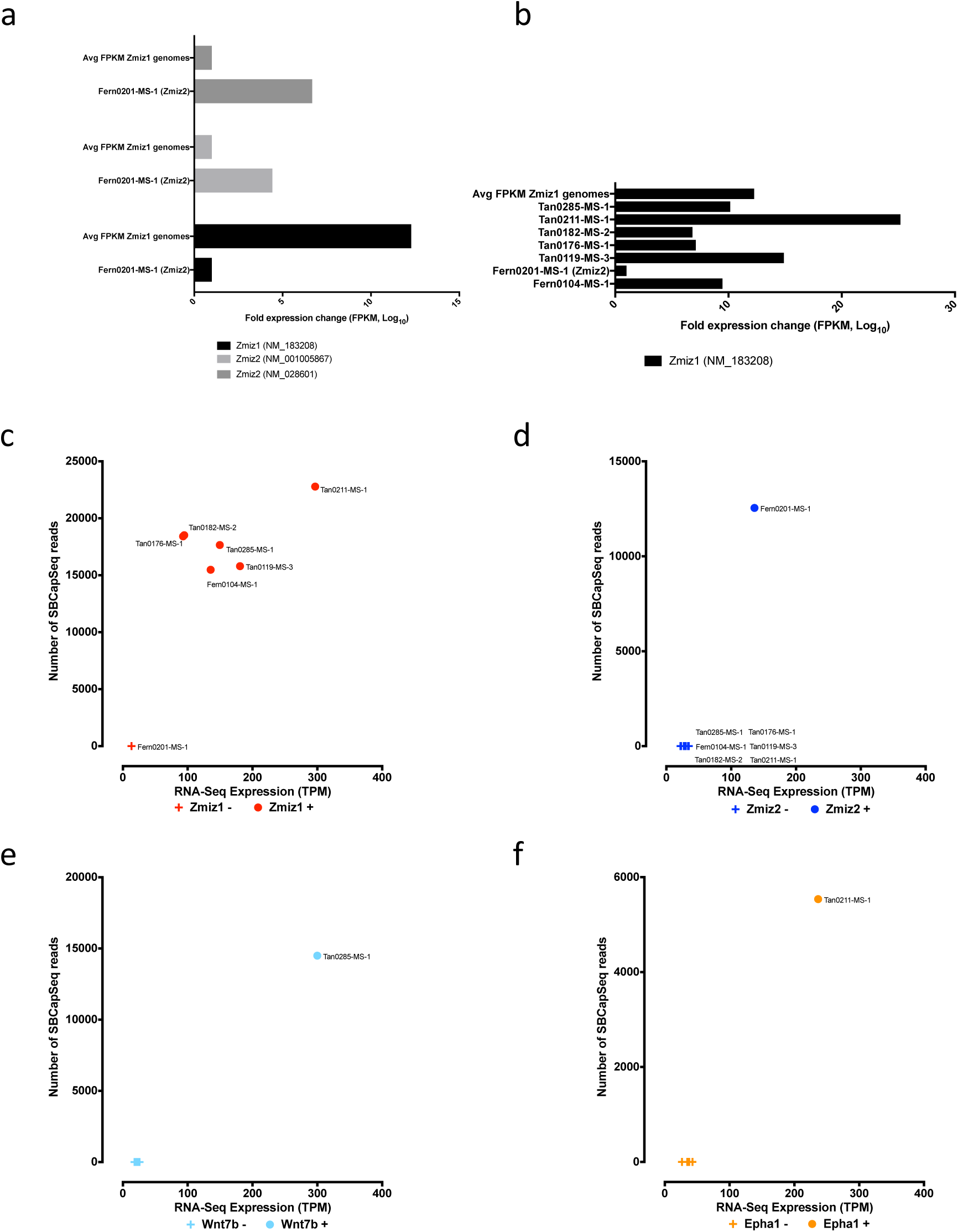
Clonally selected SB insertions affect trunk driver proto-oncogene expression in SB-cuSCC genomes. (**a**) Summary of *Zmiz1* and *Zmiz2* transcripts identified from bulk analysis of cuSCC cells by wtRNA-seq are represented as the log10 ratio of fragments per kilobase of transcript per million (FPKM) from the average of 6 genomes with SB insertions into *Zmiz1* compared to the transcripts from the single genome with *Zmiz2* SB insertion, defined by FPKM mapped reads from ribo-depleted RNA. (**b**) Transcripts identified from bulk analysis of cuSCC cells by wtRNA-seq are represented as the log10 ratio of SB insertion containing compared to wild-type transcripts, defined by fragments per kilobase of transcript per million mapped reads from ribo-depleted. (c-f) Individual panels from **Figure 5e**: multifold induction of gene expression in cuSCC masses with (**+**) high read depth activating SB insertion events among 4 candidate oncogenic drivers compared with normal gene expression levels in cuSCC tumors without (**–**) SB insertions.

**Supplementary Figure 14:**
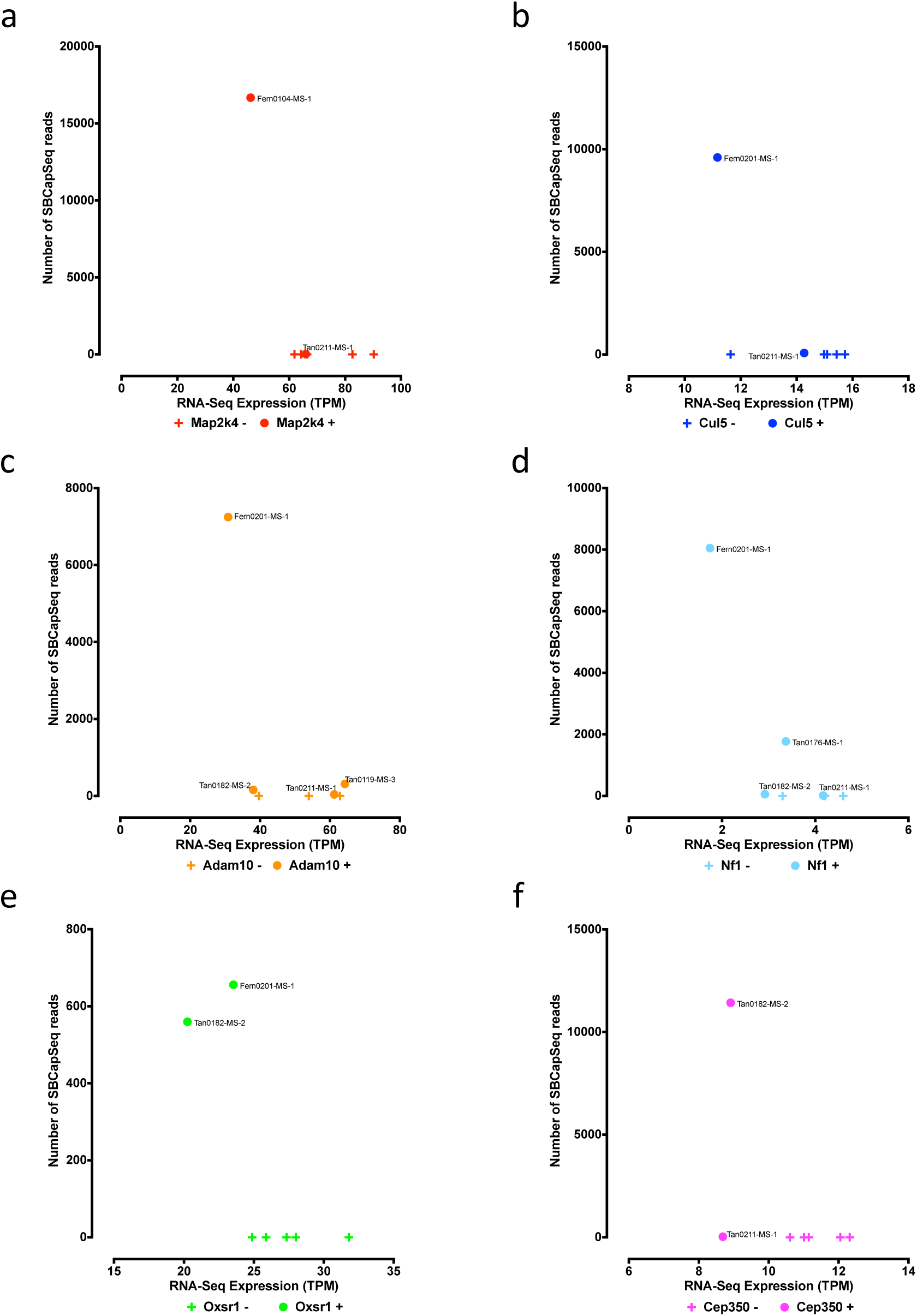
Clonally selected SB insertions affect trunk driver genes by inactivating expression in SB-cuSCC genomes. (**a-f**) Individual panels from **Figure 5f**: reduced gene expression in cuSCC masses with (**+**) high read depth inactivating SB insertion events among 6 candidate tumor suppressor drivers compared with normal gene expression levels in cuSCC tumors without (**–**) SB insertions.

**Supplementary Figure 15:**
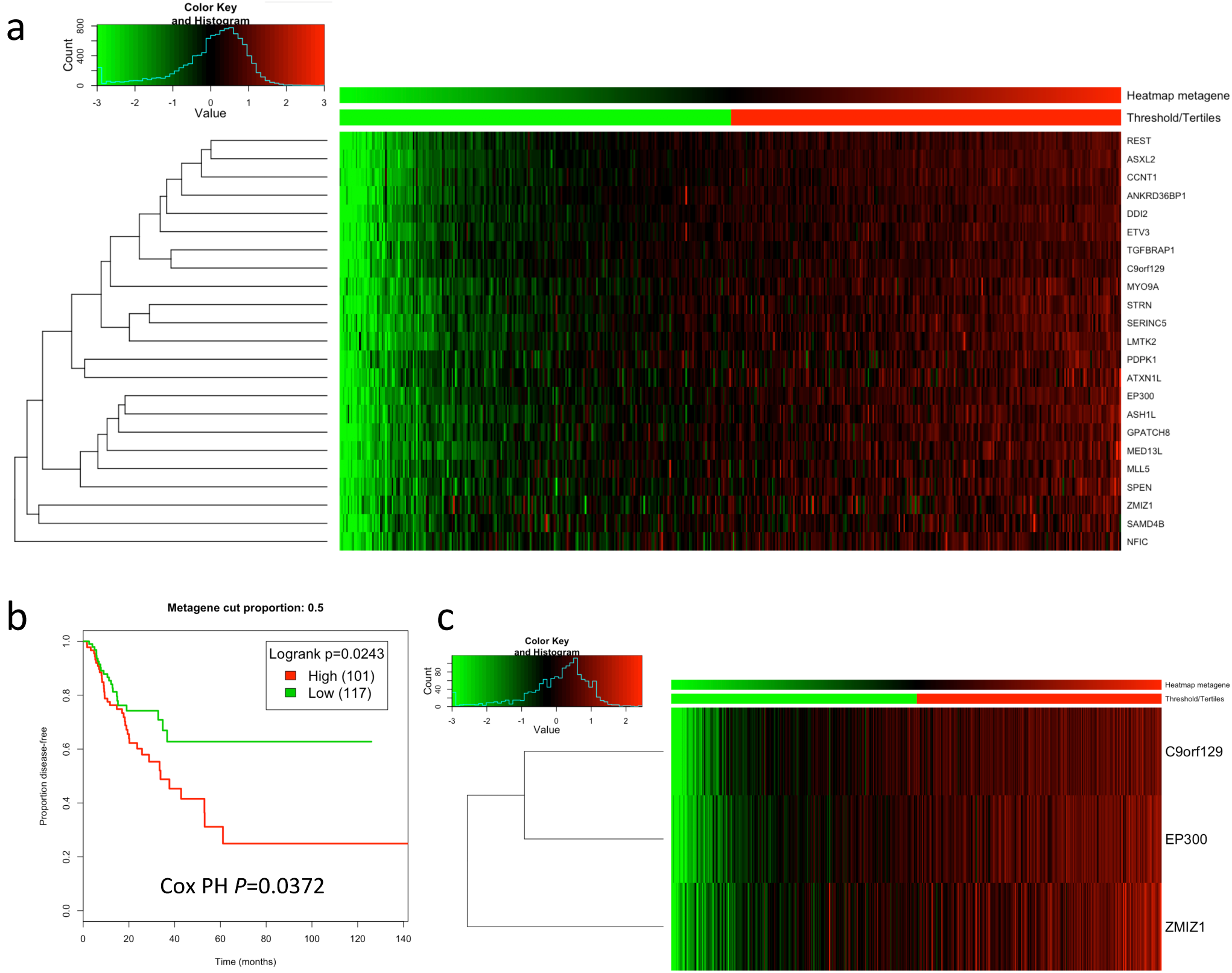
*ZMIZ1* metagene within the TCGA Head & Neck Squamous Cell Carcinoma (hnSCC) RNA-seq dataset. (**a**) *ZMIZ1* metagene heatmap constructed using singular value decomposition of human hnSCC RNA-seq dataset from TCGA consisting of a 23-gene signature from **Figure 5e**. (**b**) Survival plot of patient outcomes in human hnSCC (**b**) based on the expression of a *ZMIZ1*–centric metagene heatmap (**c**) constructed using singular value decomposition of human hnSCC RNA-seq dataset from TCGA, reveal poor patient survival outcomes predicted by increasing expression of *ZMIZ1* and closely regulated genes consisting of a 3-gene signature: *ZMIZ1, EP300*, and *C9orf129*.

**Supplementary Figure 16:**
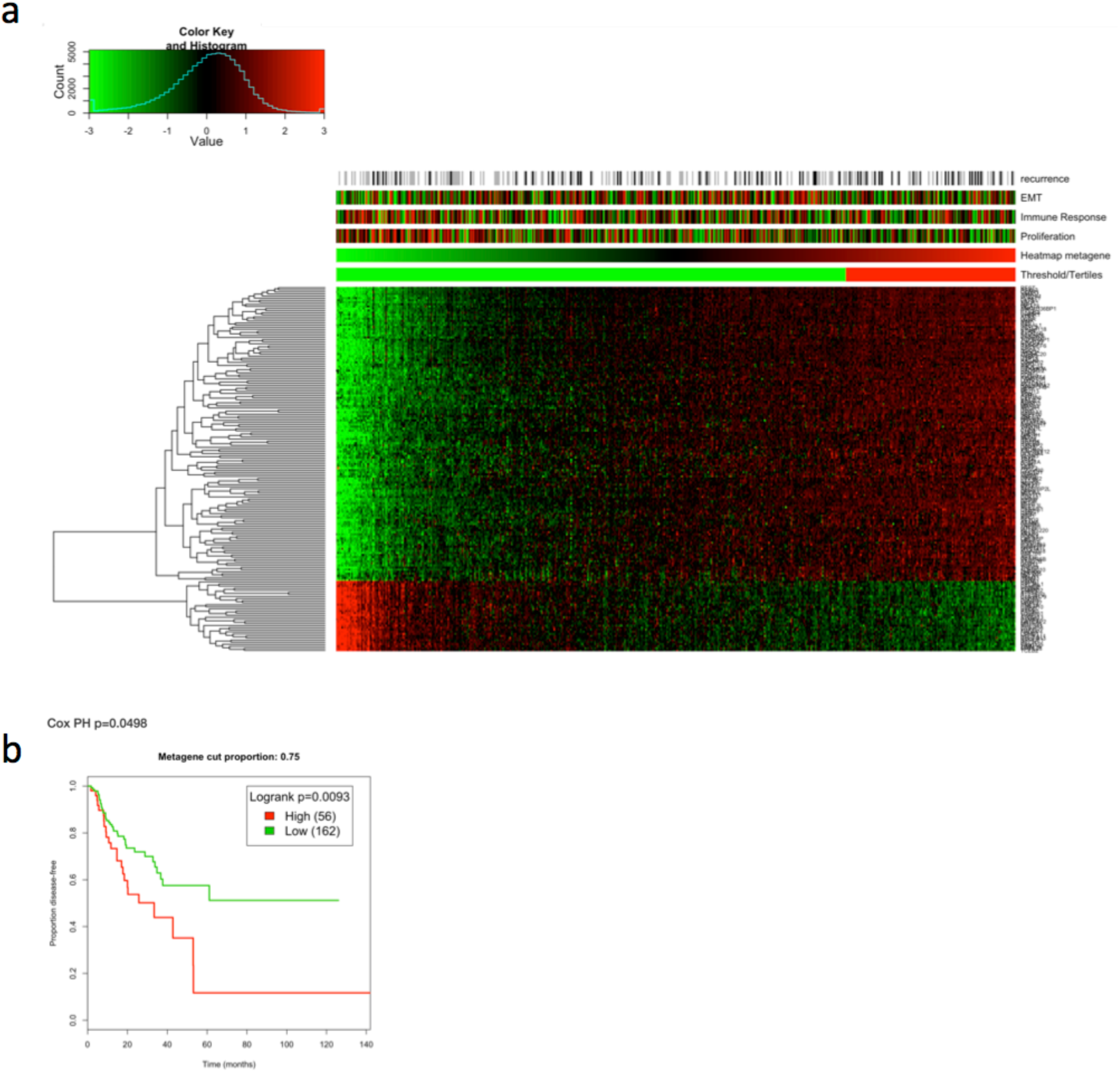
*KMT2C (MLL3)* metagene within the TCGA Head & Neck Squamous Cell Carcinoma (hnSCC) RNA-seq dataset. (**a**) *KMT2C* metagene heatmap and (**b**) patient survival outcomes.

**Supplementary Figure 17:**
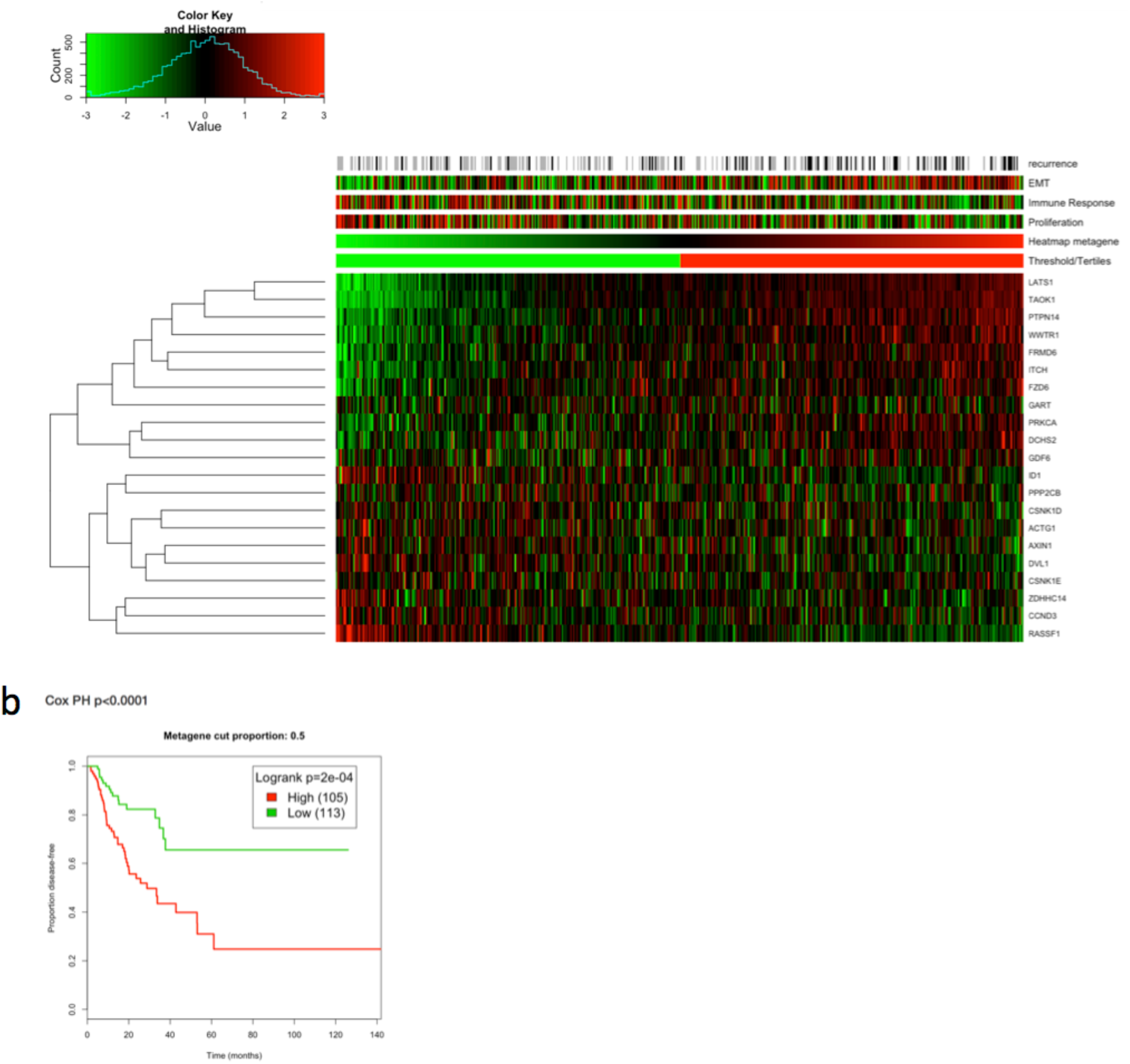
HIPPO pathway metagene within the TCGA Head & Neck Squamous Cell Carcinoma (hnSCC) RNA-seq dataset. (**a**) HIPPO pathway metagene heatmap and (**b**) patient survival outcomes.

**Supplementary Figure 18:**
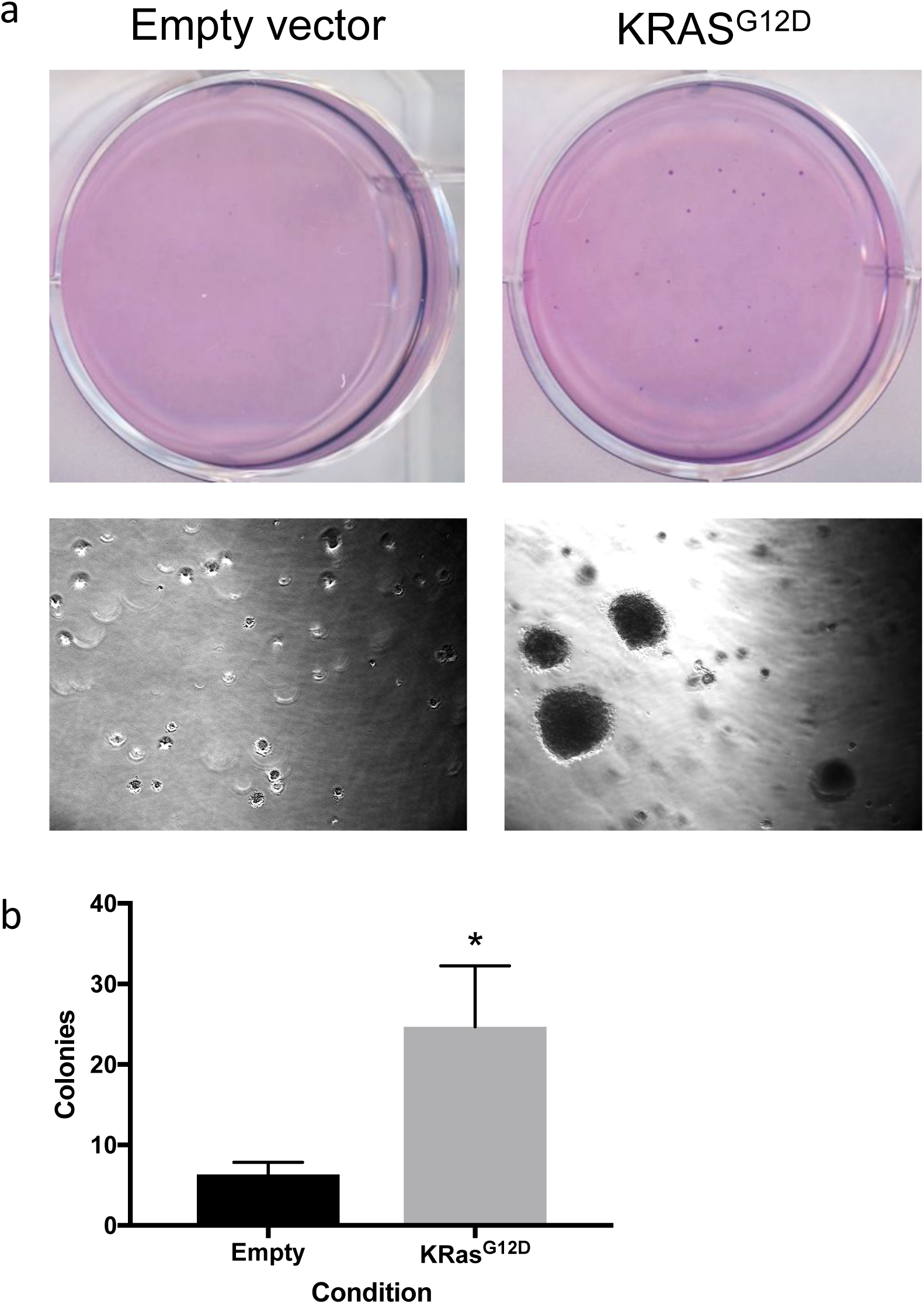
*In vitro* functional validation of oncogenic KRAS^G12D^ in immortal keratinocytes. (**a**-**b**) Assessment of keratinocyte transformation in response to expression of oncogenic KRAS^G12D^ by anchorage-independent soft agar assay (n=3). (**a**) Images of colonies stained with crystal violet solution. (**b**) Statistical significance was tested by unpaired t-test. \**P* = 0.015.

**Supplementary Figure 19:**
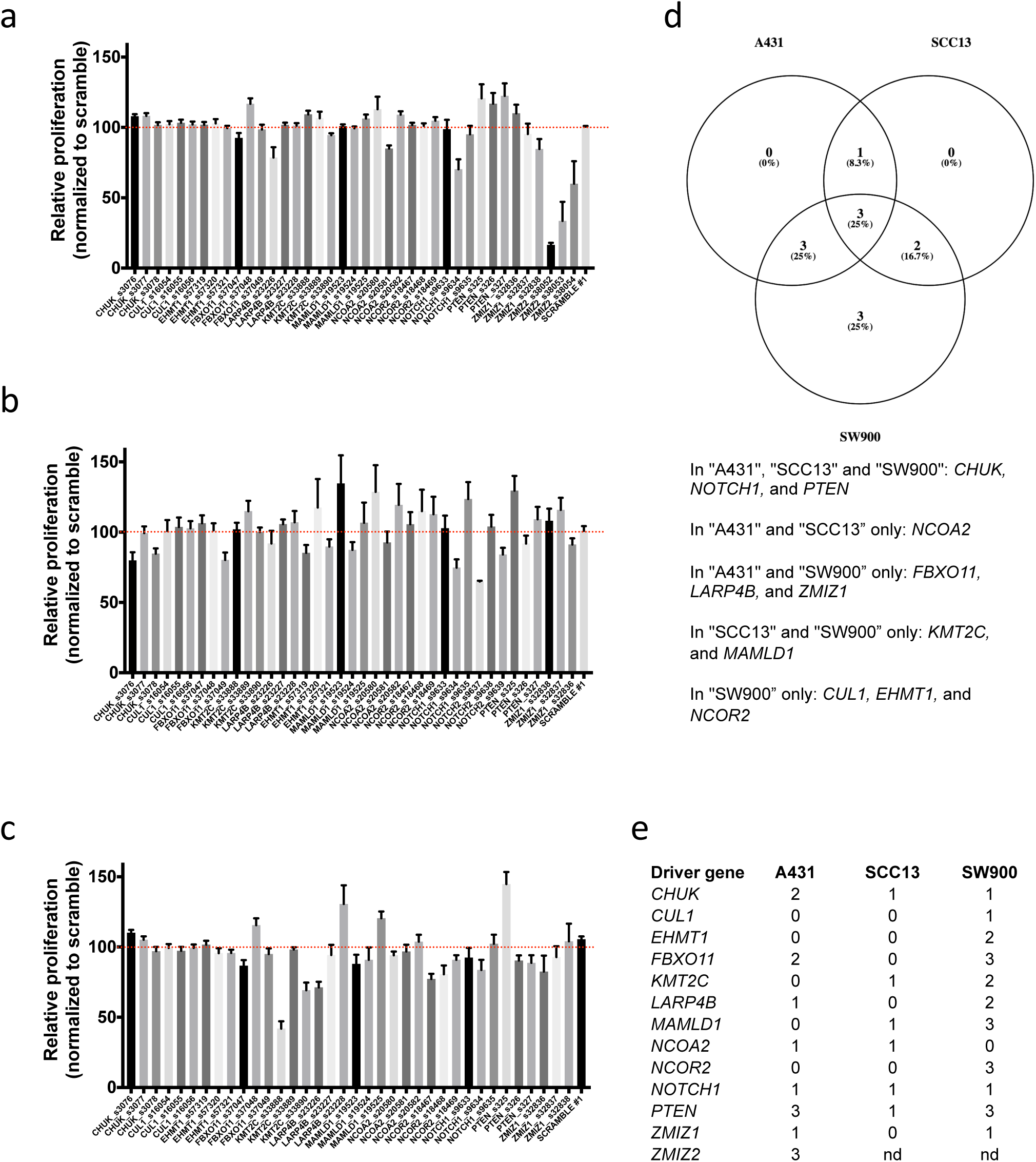
siRNA-mediated driver gene silencing in SCC cell lines. Cellular proliferation assay (WST-1) using an siRNA screen with 3 non-overlapping siRNAs from 13 genes and 2 non-targeting scramble controls (41 siRNAs, **Supplementary Table 42**) was conducted in (**a**) A431 cells (cuSCC), (**b**) SCC13 cells (cuSCC), (**c**) SW900 cells (lung SCC) at 72-hour time point. (**d**) Venn diagram of overlap between SCC cell lines, (**e**) SCC cell lines with number of statistically significant siRNAs (two-factor ANOVA, FDR adjusted-*P* <0.1).

**Supplementary Figure 20:**
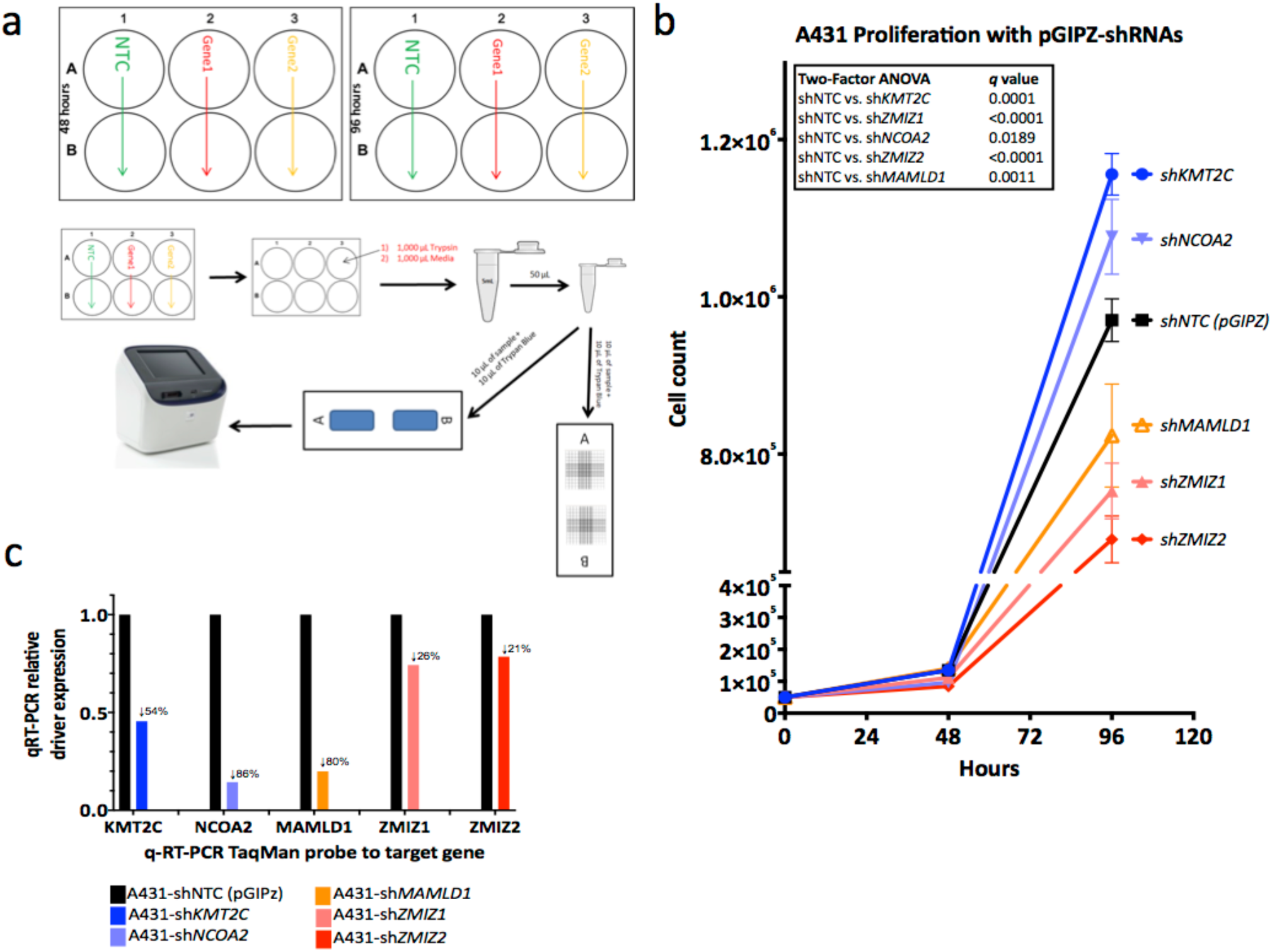
Lentiviral-shRNA-mediated driver gene silencing in cuSCC cell lines. (**a**) Experimental workflow for cellular proliferation in cuSCC cell line A431. Representative proliferation (**b**) and RNA abundance (**c**) of A431 cells with pools of 3 non-overlapping shRNAs directed against two tumor suppressor and three oncogenic drivers relative to control (shNTC) cells (two-factor ANOVA, FDR adjusted-*P* <0.1).

**Supplementary Figure 21:**
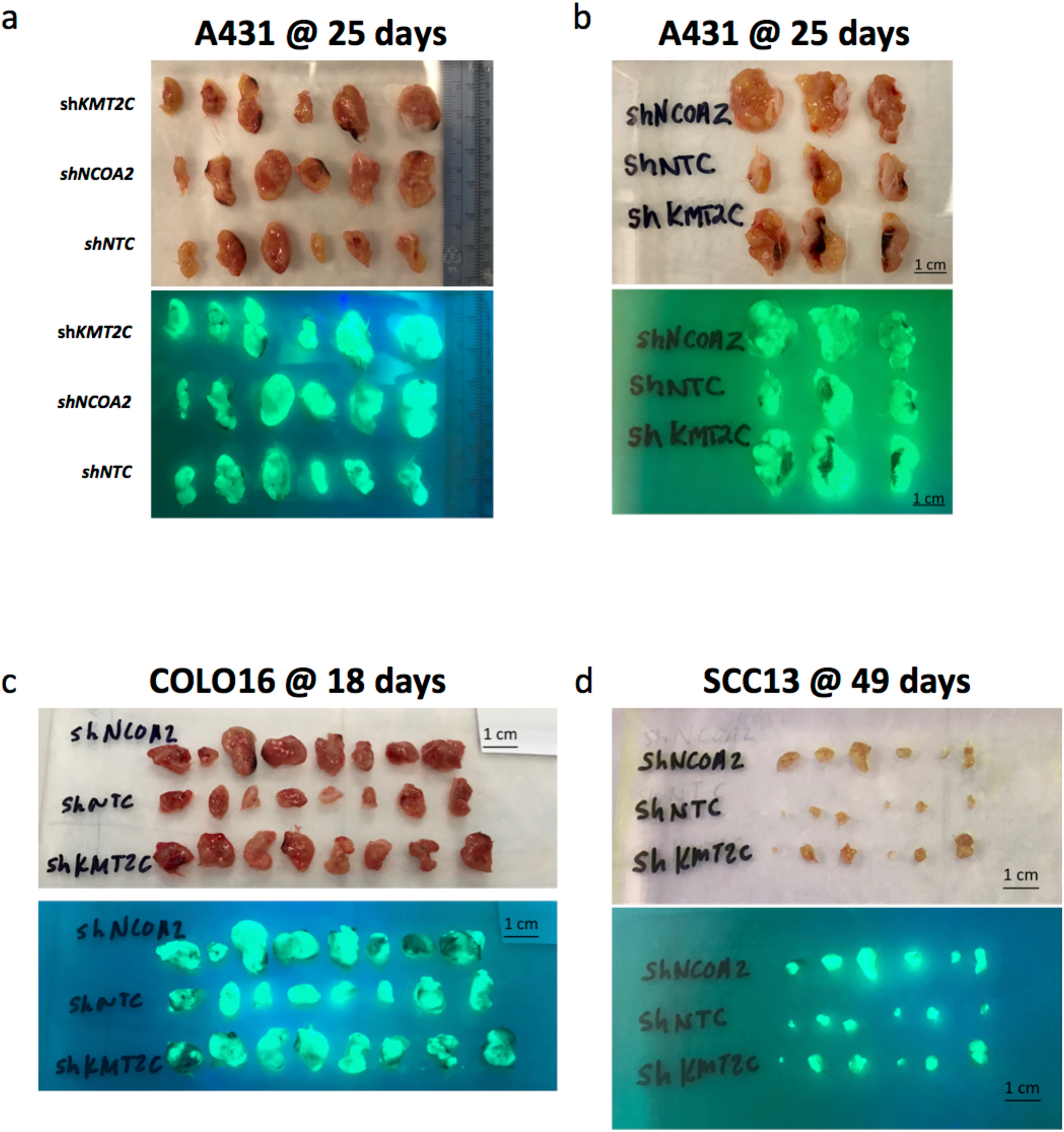
Gross photographs of cuSCC xenograft masses collected at necropsy showing robust TurboGFP expression. Xenografts of cuSCC human cell lines A431, COLO16 and SCC13 with stable lentiviral constructs expressing shRNAs directed knockdown of target drivers *KMT2C* (shKMT2C) or *NCOA2* (shNCOA2) relative to a non-targeting control (shNTC) grown in *NSG* immunodeficient mice.

**Supplementary Figure 22:**
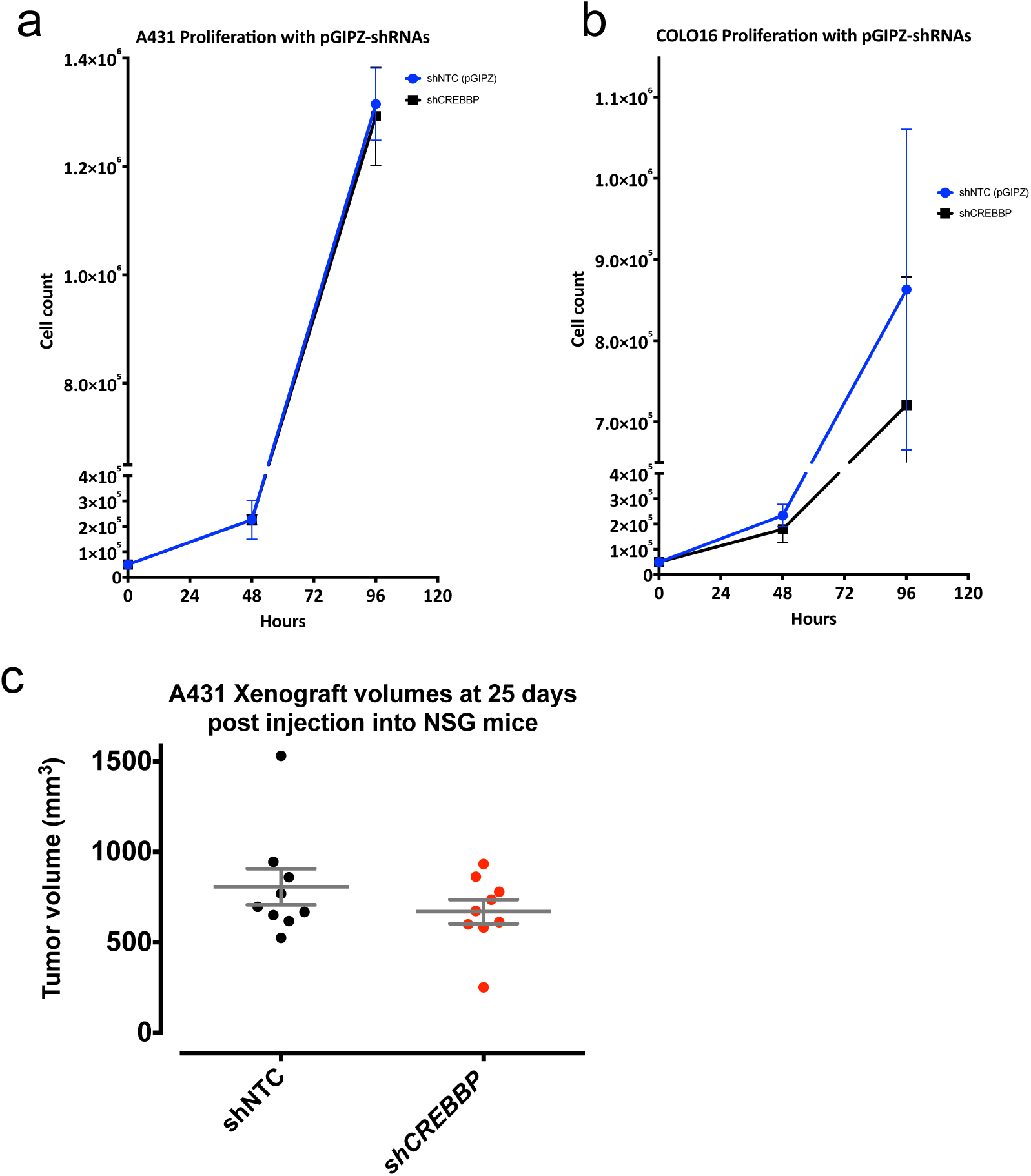
*CREBBP* knockdown does not alter proliferation rate in cuSCC cell lines. (**a**) 96-hour proliferation assay (n=3, error bars SEM). (**b**) *In vivo* xenograft assay of COLO16 shNTC or shCREBBP into NSG immunodeficient mice (n=9 per condition).

## Methods

### Mice used for SB screens

The following alleles were used to construct the SB-driven mouse model of multiple solid tumor histologies: *Actb-Cre* (FVB/N-Tg(ACTB-cre)2Mrt/J) (Lewandoski et al., 1997);*Trp53*^*flox/+*^ (FVB.129P2-*Trp53*^*tm1Brn*^/Nci) (Jonkers et al., 2001); *Trp53*^*LSL-R172H/+*^ (129S4-*Trp53*^*tm2Tyj*^/Nci) (Olive et al., 2004); T2/Onc2(TG.12740) (TgTn(sb-T2/Onc3)12740Njen) (Dupuy et al., 2009)); and *Rosa26*-LSL SBase or SBase^LSL^; (Gt(ROSA)26Sor^tm2(sb11)Njen^) (Starr et al., 2009)). The resulting cohorts of mice were on mixed genetic backgrounds consisting of C57BL/6J, 129, C3H and FVB. Genotyping by PCR assays with primers specific to the alleles was performed. No sample size estimate was used to determine the number of mice for aging. Mice were bred and maintained in accordance with approved procedures directed by the respective Institutional Animal Care and Use Committees in the National Cancer Institute Frederick National Lab, A*STAR Biological Resource Centre, Houston Methodist Research Institute, and Moffitt Cancer Center. Animals were co-housed for the duration the experiment, except in rare cases where single mice were separated based on vet recommendation due to fighting or near tumor burden end-point. Gross necropsies were performed and all masses were documented and prepared for subsequent analysis. See **Supplementary Figure 1** for details of mouse crosses. Both sexes were used for experiments. No randomization was performed; mice were assigned to groups based on genotype. No blinding was performed.

### Histological analysis

Histological analysis of spleen was performed on 5-μm sections of formalin-fixed, paraffin-embedded (FFPE) specimens stained with hematoxylin and eosin. As expected, robust nuclear staining of SB transposase (SBase) was confirmed by immunohistochemistry on FFPE tissues after antigen retrieval (pH 9) and endogenous peroxidase inhibition followed by overnight incubation with mouse antibody to SBase (anti-SBase; R&D Systems; pH 9; 1:200 dilution). After incubation with primary antibody, chromogen detection (with HRP polymer, anti-rabbit or anti-mouse, with Envision System from Dako) and hematoxylin counterstaining were performed per manufacturer’s instructions. Genomic DNA (gDNA) was isolated from flash frozen necropsy specimens using Qiagen Gentra^®^ Puregene^®^ DNA isolation kit protocol for tissue.

### Mapping transposon insertion sites using the splink_454 method

SB insertion reads were generated by 454 GS Titanium sequencing (Roche) of pooled splinkerette PCR reactions with nested, barcoded primers was performed(Mann et al., 2012; Mann et al., 2015; March et al., 2011). Pre- and post-processing of 454 reads to assign sample DNA barcodes, filter out local hopping events from donor chromosomes, and map and orient the SB insertion sites across the entire nuclear genome of the mouse was performed. All SB insertions from donor chromosomes were filtered out prior to identification of common insertion sites using the Gaussian kernel convolution (GKC) (de Ridder et al., 2006; March et al., 2011) and SB driver analysis (Newberg et al., 2018b) methods.

### Mapping transposon insertion sites using the SBCapSeq method

Full details for the SBCapSeq protocol (Mann et al., 2016b) optimized for sequencing from solid tumors, will be published elsewhere (Mann *et al*., in preparation; for general protocol and concept, see (Mann et al., 2016a; Mann et al., 2016b; Mann et al., 2019b)). Briefly, for selective SB insertion site sequencing by liquid hybridization capture, gDNA (0.5 µg per sample) of either bulk tumor specimens or single cell WGA genomes was used for library construction using the AB Library Builder™ System, including random fragmentation and ligation of barcoded Ion Xpress™ sequencing adapters. Adapter-ligated templates were purified by Agencourt^®^ AMPure beads^®^ and fragments with insert size of 200±30bp were excised, purified by Agencourt^®^ AMPure^®^ beads, amplified by 8 cycles of adapter-ligation-mediated polymerase chain reaction (aLM-PCR), and purified by Agencourt^®^ AMPure^®^ beads with elution in 50 µl of TE (1X Tris-EDTA [Ethylenediaminetetraacetic acid], pH8). Capture hybridization of single or multiplexed up to 12 barcoded libraries (60 ng per sample) was performed using custom xGen^®^ Lockdown^®^ Probes (IDT; full details available at https://doi.org/10.35092/yhjc.11441001.v1 (Mann et al., 2019b)). All 120-mer capture and blocking oligonucleotide probes were purchased from IDT as Ultramer DNA Oligos with Standard Desalting. Bar-coded single and multiplex captured library fragments were further amplified by 12 cycles of LM-PCR and run using an Agilent 2100 Bioanalyzer or TapeStation to estimate enrichment. Bar-coded single and multiplex captured libraries were quantified by Qubit^®^ Fluorometer and quantitative Real Time-PCR (qRT-PCR) were used to dilute libraries for template preparation and Ion Sphere™ Particle (ISP) loading using Ion Chef™ System and sequencing on the Ion Proton™ platform with PIv3 semiconductor wafer chips per manufacturers recommended instructions. High-throughput sequencing of up to 39 multiplex captured libraries was carried out per PIv3 chip to achieve at least 1.5 million reads per barcode. Reads containing the transposon IRDR element were processed using the SBCapSeq bioinformatic workflow as described(Mann et al., 2016b).

### SB Driver Analysis

BED formatted files containing SB insertions from each of the histologically verified SB-cuSCC, SB-cuKA, and SB-cuSK cohort specimens (**Supplementary Tables 2**,**4**,**7**,**11**,**14**,**17**,**21**) were used to perform SB Driver Analysis to identify statistically significant discovery, progression, and trunk driver genes that contain more SB insertions than expected by chance and were recurrently altered in three or more tumors (Newberg et al., 2018b). Discovery and progression SB Driver Analysis considered all SB insertion events; trunk SB Driver Analysis considered only insertions represented by 5 or more reads. Statistically significant discovery drivers were defined as adjusted *p*-values using false discovery rate (FDR) multiple testing correction (*q*-value<0.5). Statistically significant progression and trunk drivers were defined as adjusted *p*-values using family-wise error rate (FWER) multiple testing correction (FWER adjusted-*P*<0.05). Statistically significant drivers on donor and non-donor chromosomes analyses were performed separately and combined into a single driver lists (**Supplementary Tables 2**,**5-6**,**12-13**,**15-16**). Due to a local-hopping phenomenon known to occur with SB, where transposition events are biased to occur in cis along the donor chromosome more frequently than in trans to non-donor chromosomes in the genome, insertions from the donor chromosome are typically filtered away computationally before candidate cancer genes are identified. However, doing so in cohorts derived from mice with only a single SB donor allele (like our cuSCC cohorts) means that genome-wide driver analysis is not possible, because all genes on the donor chromosome are censored and are not reported even though they may contribute to the overall tumor burden. Since we obtained quantitative SBCapSeq datasets in this study, we evaluated whether all genes on the donor chromosome should be censored or if perhaps, as observed in *PiggyBac* transposon genomes, only a relatively small portion of the donor chromosome locus, occurring in close proximity to the donor concatemer insertion site, demonstrate significant bias compared with the frequency of SB insertions into non-donor chromosomes. In the cuSCC cohorts sequenced with SBCapSeq, we found that donor chromosome SB insertion site frequencies are significantly higher between chr9:79,000,000 and chr9:95,000,000 and drop to the genome wide average for non-donor chromosomes over the rest of chromosome 9 (**Supplementary Figure 2b**), suggesting that filtering all SB donor chromosome insertion events may not be warranted. To confirm and extend this observation, we used all tumors within the SBCDDB with chromosome 9 donor alleles and identified that the same region between chr9:79,000,000 and chr9:95,000,000 had a higher frequency of SB insertions than the rest of chromosome 9 loci, which again matched non-donor chromosome insertion frequencies (**Supplementary Figure 2a**). Thus, we ran SB Driver Analysis on tumors derived from mouse cohorts harboring an SB T2/Onc3 TG.12740 donor allele chromosome 9 (cuSCC83_454, cuSCC60_SBC, and cuKA11_SBC), by excluding the 81 genes that map between *Filip1* at chr9:79663368-79825689 and *Tfdp2* at 96096693-96224065 from the driver gene output files.

### Microarray gene expression analysis

Gene expression profiling of histologically confirmed SB-cuSCC masses selected by presence or absence of *Zmiz1* insertion by 454Splink sequencing was performed using Affymetrix microarrays, as described previously (Mann et al., 2016b). Briefly, 100ng of total RNA for each sample was extracted using a NORGEN Biotek Animal Tissue RNA Purification kit (Cat #25700) followed by labeled with an Affymetrix 3’ IVT Express kit (Cat # 901229) using the manufacturer’s instructions. Labeled samples were hybridized to Affymetrix GeneChip Mouse Genome 430 2.0 Arrays, and scanned at the University of Otago Genomics & Bioinformatics Facility. Raw data processing used R (version 2.15) (Team, 2013) with the “rma” function of the “affy” package (Gautier et al., 2004), including quantile normalization but no background correction. Quality assessment of the microarray data was performed using R (R_Core_Team, 2008; 2013) or the “affyQCReport” package (Parman et al., 2005). Data analysis and differential gene expression analysis were performed using Python, R (R_Core_Team, 2008; 2013) and visualized using the R Shiny package in RStudio.

### Transcriptome Sequencing

Total RNA was isolated from flash frozen necropsy specimens using *mir*Vana™ miRNA Isolation Kit (Ambion^®^ by Life Technologies, AM1560) from a shaved portion of each pathology verified cuSCC mass from 7 tumors for which sufficient tissues were available and the presence of a high read depth *Zmiz1* or *Zmiz2* trunk driver insertion was identified by SBCapSeq. Whole transcriptome RNA-Seq (wtRNA-Seq) libraries were prepared from total RNA (5 μg per sample) plus ERCC RNA Spike-In (Ambion^®^ by Life Technologies, 4456739) followed by selective ribosomal RNA (rRNA) depletion using the RiboMinus™ Eukaryote System v2 (Ambion^®^ by Life Technologies, A15026). rRNA-depleted total RNA (500 ng per sample) was used for whole transcriptome library construction according to the Ion Total RNA-Seq Kit for the AB Library Builder™ System (Life Technologies, 4482416) protocol for barcoded libraries, Ion Sphere™ Particle (ISP) loading using Ion Chef™ System and sequencing on the Ion Proton™ platform with PIv3 semiconductor wafer chips per manufacturers recommended instructions. Up to 4 RNA-Seq libraries were multiplex sequenced on a PIv3 chip twice to achieve >3 GB (∼40 million reads) per specimen. The Life Technologies Torrent Suite software was used to perform checking of the raw sequence data before the generation of sequencing read files in FASTQ format and minimally processed mRNA-Seq and wtRNA-Seq reads were processed using the Bowtie2 (Langmead and Salzberg, 2012) and Tophat (Trapnell et al., 2009) algorithms to align reads to a custom version of the mouse mm9+pT2/Onc genome, as described (Mann et al., 2016b). The detection of SB fusion events, defining novel SB fusion transcripts, and measuring transcript abundance in SB cuSCC tumor specimens from RNA-Seq data were performed as described (Mann et al., 2016b).

### Biological Pathway and Process Enrichment Analysis

Enrichr (Chen et al., 2013; Kuleshov et al., 2016), an online analysis tool for human and mouse gene-set enrichment, was used to identify specific signaling pathways and processes enrichment using the various cohort candidate cancer driver genes and/or their human orthologs (**Supplementary Tables 28**,**32-33**). The online STRING network enrichment analysis tool (Szklarczyk et al., 2019) was also used to determine functional connections between trunk drivers from the cuSCC and cuKA cohorts.

### Human cancer cell lines

The cutaneous squamous cell carcinoma A431(CRL–1555™) and grade IV lung squamous cell carcinoma SW900 (HTB–59™) cell lines, were purchased from American Type Culture Collection (ATCC^®^) and grown according to the manufacturer’s suggested conditions in complete medium (1× DMEM (ATCC, 30–2002) for A431cells or 1× ATCC-formulated Leibovitz’s L–15 Medium, Catalog No. 30–2008 medium (ATCC, 30–2001) for SW900 cells) supplemented with 10% FBS and 1× penicillin-streptomycin grown at 37 °C in 5% CO2. The cutaneous squamous cell carcinoma SCC13 cell line (Rheinwald and Beckett, 1981; Rheinwald and Green, 1977), were obtained from Harvard Skin Disease Research Center (HSDRC) and grown using the human keratinocyte culture methods provided by the HSDRC. The cutaneous squamous cell carcinoma COLO16 cell line (Vin et al., 2013) were a gift from Dr. K. Y. Tsai and grown in DMEM/Ham’s F12 50/50 media supplemented with cholera toxin (Sigma, C0852), insulin (Sigma, I5500-50MG), epidermal growth factor (Serotec/Bio-Rad, EGF-1), hydrocortisone (Sigma, H4881-1G), liothyronine (Sigma, T6397-100MG) and apo-transferrin (Sigma, T2252-100MG). All cell lines were verified human origin and free from pathogens and mycoplasma. Mycoplasma infection monitoring was performed using MycoAlert Detection Kit (Lonza) and only mycoplasma-free cultures were used.

### siRNA screening

Pre-designed siRNA oligonucleotides were purchased from Thermo Fisher Scientific and are listed in **Supplementary Table 41**. For each gene, three non-overlapping siRNA oligos were transfected using Lipofectamine RNAiMAX transfection reagent (Thermo Fisher Scientific) according to the manufacturer’s instructions. Briefly, 4 × 10^3^ cells were seeded into 96-well plates and cultured for 24 hours at 37 °C in complete culture medium. At 24 hours, single siRNAs were transfected and incubated at 37 °C. Cellular proliferation was measured with a WST–1 cell proliferation assay system (Takara). WST–1 reagent was added to each well and incubated for 2 hours at 37 °C. The colorimetric absorbance at 450 nm and 650 nm (reference) was measured with an Infinite 200 pro multimode reader (TECAN).

### Generation of stable shRNA-expressing cell lines

High titer lentiviral particles for pGIPZ-shRNA constructs targeting each of the 12 cutaneous candidate driver genes, and one non-targeting control, were purchased from Thermo Scientific Open Biosystems (**Supplementary Table 43**). Cells were plated at a density of 5 × 10^4^ cells per well in a 24-well plate in complete media 24 hours prior to infection. The following day, cells were plated with serum-free culture medium containing 8 µg/mL polybrene (Millipore), and transduced as pools of three independent shRNAs to different exons of each target gene, or individually for the control, at multiplicity of infection of 6. Puromycin selection was added the following day at concentrations of 1.0 to 3.0 µg/mL puromycin (Thermo Fisher Scientific) in complete media, and replaced every three until stable lines were achieved.

### Proliferation assay

To assess cuSCC progression, cuSCC cell lines were seeded in 6 well plates at a density of 5 × 10^4^ cells per well in complete media. Cell numbers for each condition were counted after 48- and 96-hour post-seeding. To assess transformation by proliferation, HaCaT cells were seeded at a density of 5 × 10^3^ cells per well and cell counts performed after seven days.

### Soft agar assay

To assess anchorage-independent growth, HaCaT cells were seeded at a density of 1 × 10^4^ cells per well in a six well plate in 0.3% agar, over a layer 0.6% agar. Complete media was added to each well the following day, and replaced every two days for four weeks. At end point, cells were fixed in 20% methanol and 0.0025% crystal violet. Washes were done with milliQ water until low background was achieved, and plates were scanned, and colonies counted for each condition. KRAS^G12D^ expression vector (Origene cat# RC400104) was a kind gift from Dr Karen Mann. The empty vector (pCMV6-Entry) was generated by removing the KRAS^G12D^ cDNA using the *SgfI* and *MluI* restriction enzymes, gel purified, and ligated to obtain an empty plasmid. HaCaT cells were transfected with the expression plasmids and selected with 800ug/mL G418 to obtain stable cell lines. For the soft agar assay was performed as described above.

### qRT-PCR

Total RNA was purified and DNase treated using the RNeasy Mini Kit (Qiagen). Synthesis of cDNA was performed using SuperScript VILO Master Mix (Life Technologies). Quantitative PCR analysis was performed on the QuantStudio 12K Flex System (Life Technologies) or 7900HT Sequence Detection System (Applied Biosystem). All signals were normalized to the levels of GAPDH TaqMan probes. TaqMan probes were obtained from Life Technologies (**Supplementary Table 41**).

### Xenografts

One million cells were prepared for injection into the left flank of randomized selected male and female immunodeficient NSG (NOD.Cg-Prkdcscid;Il2rgtm1Wjl/SzJ; JAX, 005557; 6–15 weeks old) mice. Allocation to study groups was random. Xenograft measurements were taken twice weekly using digital calipers while mice were conscious but restrained by an experimenter familiar with collecting caliper measurements of xenografts and blinded to the experimental group designations. GFP fluorescence was visualized at the time of caliper measurement. Ellipsoid tumor volumes were calculated as volume (mm3) = 0.52(length (mm) × width2 (mm2)), where the two longest axes, length and width, were the major and minor diameter measurements, respectively; width2 represents an assumption that the xenograft depth was equivalent to the diameter of the minor axis. Statistical significance of tumor volumes at defined time points was determined by either one-way t test for all cohorts relative to shNTC or by two-way, repeated-measures ANOVA with Bonferroni correction for multiple comparisons, as indicated.

### Western blot

Whole cell or tissue lysates were prepared in RIPA buffer supplemented with protease and phosphatase inhibitors, and samples sonicated to achieve optimum lysis. Protein concentrations were quantitated using the BCA assay, and 20 µg of lysates were loaded into 8% SDS-polyacrylamide gels. Gels were resolved at 80V for 120 mins, transferred onto nitrocellulose membrane for 70 mins at 0.35A with transfer buffer containing 20% methanol, and blocked for one hour at room temperature with 5% w/v BSA in 0.1% TBST. Membranes were incubated with primary antibodies in 5% w/v BSA in 0.1% TBST. Antibodies used in this study are: Total ERK (1:1000, Cell Signaling #9102), phospho ERK (1:1000, Cell Signaling #4695), GAPDH (1:5000, Santa Cruz sc-69778), anti-mouse and anti-goat secondary antibodies (1:10000, LiCOR IRDye 680/800 Cat. 925-68070/926-32211). Blots were imaged using the LiCOR Odyssey Fc system.

### Software

Unless otherwise noted, bioinformatic analysis pipelines, report generation, and figure visualization performed using bash, R (version 3.5.2), RStudio (Version 1.1.463), Python scripts, and GraphPad Prism 8 software (Version 8.1.1). Hierarchical clustering was performed in Python 2.7.10 with the scipy 0.13.0b1 toolbox using the Hamming distance metric with Ward’s linkage method. The R packages ggplot2 (Wickham, 2009) and Gviz (Henare et al., 2012) were used to generate the graphics in various figure panels.

### URLs

TgTn(sb-T2/Onc3)12740Njen; Gt(ROSA)26Sortm2(sb11)Njen double mutant mice, https://ncifrederick.cancer.gov/Lasp/MouseRepository/MouseModels/StrainDetails.aspx?StrainNum=01XGB&g=ROSA26; FVB/N-Tg(ACTB-cre)2Mrt/J mice, http://jaxmice.jax.org/strain/003376.html; Cancer Gene Census, http://cancer.sanger.ac.uk/census/; Mouse Genome Informatics (MGI database, ftp://ftp.informatics.jax.org/pub/reports/index.html; Enrichr, http://amp.pharm.mssm.edu/Enrichr/, STRING, https://string-db.org/, The Cancer Genome Atlas (TCGA), http://cancergenome.nih.gov/; FVB.129P2-Trp53tm1Brn/Nci mice, https://ncifrederick.cancer.gov/Lasp/MouseRepository/MouseModels/StrainDetails.aspx?StrainNum=01XC2&g=Trp53; 129S4-Trp53tm2Tyj/Nci mice, https://ncifrederick.cancer.gov/Lasp/MouseRepository/MouseModels/StrainDetails.aspx?StrainNum=01XM2&g=Trp53.

### Accession Codes

NCBI BioProject accessions: PRJNA580460 for whole-transcriptome RNA-Seq data; PRJNA580462 for microarray expression data.

**Supplementary Tables**. Supplementary Table datasets may be downloaded from NIH.figshare.com at (http://dx.doi.org/10.35092/yhjc.11441130) (Mann et al., 2019a). Brief descriptions of each table are provided for reference.

**Supplementary Table 1**: Tumor incidence and subgroup classifications by cohort

**Supplementary Table 2**: Sequencing projects summary by cohort

**Supplementary Table 3**: Specimen metafile data for projects sequenced using Splink_PCR protocol with 454 GS-FLX Titanium pyrosequencer

**Supplementary Table 4**: BED file of SB insertions for cuSCC83_454

**Supplementary Table 5**: Discovery and progression SB Driver Analysis for cuSCC83_454

**Supplementary Table 6**: Trunk SB Driver Analysis for cuSCC83_454

**Supplementary Table 7**: BED file of SB insertions for cuKA51_454

**Supplementary Table 8**: Discovery and progression SB Driver Analysis for cuKA51_454

**Supplementary Table 9**: Trunk SB Driver Analysis for cuKA51_454

**Supplementary Table 10**: Specimen metafile data for projects sequenced using SBCapSeq protocol with Ion Torrent Proton sequencer

**Supplementary Table 11**: BED file of SB insertions for cuSCC60_SBC.

**Supplementary Table 12**: Discovery and progression SB Driver Analysis for cuSCC60_SBC.

**Supplementary Table 13**: Trunk SB Driver Analysis for cuSCC60_SBC.

**Supplementary Table 14**: BED file of SB insertions for cuKA11_SBC

**Supplementary Table 15**: Discovery and progression SB Driver Analysis for cuKA11_SBC

**Supplementary Table 16**: Trunk SB Driver Analysis for cuKA11_SBC

**Supplementary Table 17**: BED file of SB insertions for cuSK32_SBC

**Supplementary Table 18**: Discovery and progression SB Driver Analysis for cuSK32_SBC

**Supplementary Table 19**: BED file of SB insertions for 4 cuSCC genomes selected for multi-region resequencing because they had intermixing of cuSCC and cuKA histologies

**Supplementary Table 20**: SBCapSeq read depth and analysis for 4 cuSCC genomes selected for multi-region resequencing because they had intermixing of cuSCC and cuKA histologies

**Supplementary Table 21**: BED file of SB insertions for cuSK10_SBC

**Supplementary Table 22**: Discovery and progression SB Driver Analysis for cuSK10_SBC

**Supplementary Table 23**: Contingency Tables comparing Cancer Gene Census (GRCh38 COSMICv86) driver enrichment with cuSCC cohorts

**Supplementary Table 24**: Venn diagram for overlap of Cancer Gene Census (GRCh38 COSMICv86) driver enrichment with all SB|cuSCC Trunk Driver and SB|cuKA Trunk Driver lists

**Supplementary Table 25**: Venn diagram for overlap of Cancer Gene Census (GRCh38 COSMICv86) driver enrichment with all SB|cuSCC Progression Driver and SB|cuKA Progression Driver lists

**Supplementary Table 26**: Venn diagram for overlap of Cancer Gene Census (GRCh38 COSMICv86) driver enrichment with cuSCC60_SBC Trunk Driver, cuSCC83_454 Trunk Driver, and cuKA51_454 Trunk Driver lists

**Supplementary Table 27**: Venn diagram for overlap of Cancer Gene Census (GRCh38 COSMICv86) driver enrichment with cuSCC60_SBC Progression Driver, cuSCC83_454 Progression Driver, and cuKA51_454 Progression Driver lists

**Supplementary Table 28**: Enrichr gene set pathway enrichment analysis of cuSCC drivers

**Supplementary Table 29**: Normalized microarray values per gene from RNA isolated from cuSCC genomes with and without Zmiz1 insertions

**Supplementary Table 30**: Normalized microarray values per probe from RNA isolated from cuSCC genomes with and without Zmiz1 insertions

**Supplementary Table 31**: Top 96 genes with differential expression analysis from microarray data from RNA isolated from cuSCC genomes with and without Zmiz1 insertions

**Supplementary Table 32**: Enrichr gene set pathway enrichment analysis of the top 96 differentially expressed genes from microarray data from RNA isolated from cuSCC genomes with and without Zmiz1 insertions

**Supplementary Table 33**: Enrichr gene set gene ongology term enrichment analysis of the top 96 differentially expressed genes from microarray data from RNA isolated from cuSCC genomes with and without Zmiz1 insertions

**Supplementary Table 34**: All 289 genes with differential expression analysis from microarray data from RNA isolated from cuSCC genomes with and without Zmiz1 insertions with FDR adjusted-*P*<0.0001

**Supplementary Table 35**: Summary of 7 cuSCC transcriptomes selected for whole transcriptome RNA-seq analysis

**Supplementary Table 36**: BED file of SBfusion insertions in 7 cuSCC genomes by whole transcriptome RNA-seq analysis

**Supplementary Table 37**: Venn diagram for overlap of genes with SBfusion reads detected by whole transcriptome RNA-seq analysis and cuSCC60_SBC discovery drivers

**Supplementary Table 38**: Venn diagram for overlap of genes with SBfusion reads detected by whole transcriptome RNA-seq analysis and all cuSCC drivers

**Supplementary Table 39**: Transcripts per million (TPM) normalized whole transcriptome RNA-seq values per gene from RNA isolated from cuSCC genomes with and without Zmiz1 insertions

**Supplementary Table 40**: Fragments Per Kilobase of Transcripts per Million (FPKM) normalized whole transcriptome RNA-seq values per gene transcript from RNA isolated from cuSCC genomes with and without Zmiz1 insertions

**Supplementary Table 41**: siRNA and TaqMan probes used in this study

**Supplementary Table 42**: siRNA primary screen results

**Supplementary Table 43**: Lentiviral vectors containing shRNAs used in this study

## Acknowledgments

M.B.M. and K.M.M. are supported by start-up funds provided by the H. Lee Moffitt Cancer Center and Research Institute, a National Cancer Institute-designated Comprehensive Cancer Center through the Cancer Center Support Grant (NCI P30-CA76292; PI: J. Cleveland). M.B.M. was also supported by Career Enhancement Program Grant Award from the NCI Moffitt Skin Cancer SPORE (5P50CA168536-05; PI: V. Sondak). N.G.C. and N.A.J. were supported by the Biomedical Research Council, Agency for Science, Technology, and Research, Singapore and as CPRIT Scholars in Cancer Research from the Cancer Prevention Research Institute of Texas. The authors wish to thank the Copeland–Jenkins labs in Singapore and Houston for helpful discussions; we thank Tyler Jacks (MIT) for P53 LSL R172H mice; Anton Berns (NKI) for P53 floxed mice; Gail Martin (UCSF) for Actin-beta-Cre mice; James G. Rheinwald and the Harvard Skin Disease Research Center for the SCC13 cell line; David Adams, Theo Whipp, Richard Rance, Alistair Rust, and the Wellcome Trust Sanger Institute sequencing and informatics teams for 454 sequencing; the Elsa Flores Lab for use of the LiCOR imaging system; Sean Yoder (Moffitt Cancer Center) and Dr. Chaomei Zhang (Moffitt Cancer Center) for providing sequencing services; Pearlyn Cheok, Nicole Lim, Dorothy Chen and Cherylin Wee for assistance with tumor monitoring and animal husbandry at IMCB (Singapore, Republic of Singapore), Hubert Lee and Erik Freiter for assistance with animal husbandry at HMRI (Houston, TX, USA), and Crystal Reed and Bethanie Gore for assistance with animal husbandry at MCC/USF (Tampa, FL, USA). Necropsy and histology were performed by the Advanced Molecular Pathology Laboratory, Institute of Molecular and Cell Biology (A*STAR), Singapore. Necropsy and histology were also performed by the Tissue Core Facility and Ion Torrent sequencing was performed by the Molecular Genomics Core at the H. Lee Moffitt Cancer Center & Research Institute, a comprehensive cancer center designated by the National Cancer Institute, and funded in part by Moffitt’s Cancer Center Support Grant (P30-CA076292; PI: J. Cleveland).

